# Cellular Factors Involved in Transcriptome Dynamics in Early Zebrafish Embryogenesis

**DOI:** 10.1101/2022.09.29.510050

**Authors:** Han Rauwerda, Johanna F.B. Pagano, Wim C. de Leeuw, Wim Ensink, Marina van Olst, Ulrike Nehrdich, Martijs J. Jonker, Herman P. Spaink, Timo M. Breit

## Abstract

At gastrulation in the zebrafish embryogenesis, the embryonic genome is switched on to produce transcripts that are used for the maintenance and development of the embryo. In a previous study from late blastula to mid gastrula on the transcriptomes of 179 individual embryos, we capture the transcriptome dynamics via ten gene-expression types. Here we study the factors that regulate these transcriptome dynamics by in extensive silico analyses and two small-RNA sequencing experiments. We analyzed mechanisms that would make it possible for the embryo to achieve the tight regulation of gene expression that was observed, not only during development, but also when individual embryos were compared. We found that many of the gene-expression regulatory factors that are available to the embryo are operational in the different gene-expression types and act concurrently with not one mechanism prevailing in this developmental phase. We also saw that at least one of the regulatory mechanisms, the expression of members of the miRNA-430 family again is very tightly regulated, both during development as well as when miRNA expression from individual embryos is compared.

## Introduction

Gene-expression regulation is a key process in the physiology of a cell. Through it a cell is able to develop and respond to the wide array of intracellular and extracellular signals to which it is exposed in life. The means available to a cell to regulate gene expression are manifold: the production or the rate of production mRNAs in the nucleus are for instance controlled by chromatin remodeling, nucleosome organization at promoters [1], transcription factors (TFs) interacting with transcriptionfactor binding sites (TFBSs), interaction of the transcription machinery with promoters and enhancers, pol II pausing [2] and also by structural components in the nucleus, i.e. nuclear laminae [3]. Export of mRNAs to the cytoplasm is well-regulated [4] and in the cytoplasm transcripts can be silenced and degraded via deadenylation and interaction with miRNAs. For instance, in early embryogenesis maternal mRNA can be activated via cytoplasmic polyadenylation. These processes are deployed at different timescales: chromatin remodeling can take place within seconds [5], pol II transcription takes place at a speed of tens of bases per second (minutes per transcript) [6] and nucleosome turnover can be in the order of minutes to hours [7]. Even more, transcription regulation still can have a stochastic component, which in some cases is used to create a binary switch [8]. In general, in a multicellular organism singular cell’s responses are integrated at the level of tissue and, upwards at the level of the organ and the organism, the modeling of which is reviewed by [9]. In the developing embryo, transcripts drive development [10,11], which makes transcriptome-regulation research an important topic in embryology as well. The early zebrafish embryo is primarily dependent on maternally provided transcripts that can be activated by cytoplasmic polyadenylation [12,13]. From the 64-cell stage onwards, miR-430 genes are zygotically transcribed and between the 128-cell and 512-cell stages a wider range of relatively short zygotic transcripts are observed [14,15]. Maternal transcripts are, at least partially cleared by miR-430 [16]. From the gastrulation onwards the amount of polyadenylated mRNA molecules increases steadily[17]. It is likely that this increase has a zygotic origin [15,18,19].

Until recently, transcriptome dynamics in embryogenesis was studied using pools of eggs or embryos taken at time points rather far apart during development. Nowadays, comprehensive studies on individuals and/or at a high time-resolution are feasible. Our previous study of the transcriptome in individual zebrafish eggs revealed an overall strict regulation of the maternal transcriptome, including a maternal effect that introduced mother-specific differences [20]. Recently two high-resolution time-course studies along embryonic development were carried out: in Xenopus the developmental period from egg to 65 hours post fertilization (hpf) was covered by pools of minimally 10 embryos and time points along the entire development that are minimally 30 minutes apart [17] and in a previous study of us (Figure 1B) [21] we profiled transcriptomes of 180 individual zebrafish embryos that are on average one minute apart in a period from late blastula to mid gastrula. Both studies indicate that the gene-expression fold changes during gastrulation are modest and transcriptome dynamics are described by half periods (off - on - off) of hours rather than minutes. In our study we showed that this tight gene-expression regulation also applies when various individual embryos are compared. We identified in this developmental period, ten types of distinct geneexpression profiles (Figure 1D) and determined the exact expression starting and stopping points for a number of genes. Noticeable, several genes showed expression at two or three distinct levels that strongly related to the spawn an embryo originated from (Figure D). In the current study, we focus on several cellular factors that might be involved in the dynamics of the gene-expression types. We investigated: i) the genome location of genes in these types, because nucleosome organization is localized on chromosomes; ii) the gene size since in early development, the size of the expressed genes is increasing [14]; iii) the gene promoter regions for specific TFBSs; iv) the 3’UTRs for cytoplasmic polyadenylation motifs; v) the 3’UTRs for miRNA targets; vi) overrepresentation of specific pathways and gene ontology (GO) categories in the gene-expression types. In all these analyses, we also included the set of genes that showed a distinct multi-level expression pattern (Figure 1D) that correlates with spawn. To obtain insight in the regulation of the gene-expression regulating factors, we investigated the miRNA expression of individual embryos at approximately 50% epiboly (Figure 1C) using the same spawns as in our high-resolution gene-expression time course, plus miRNA expression of individual embryos spanning the entire zebrafish embryogenesis including unfertilized eggs and adult zebrafish (Figure 1A).

**Figure 1.**
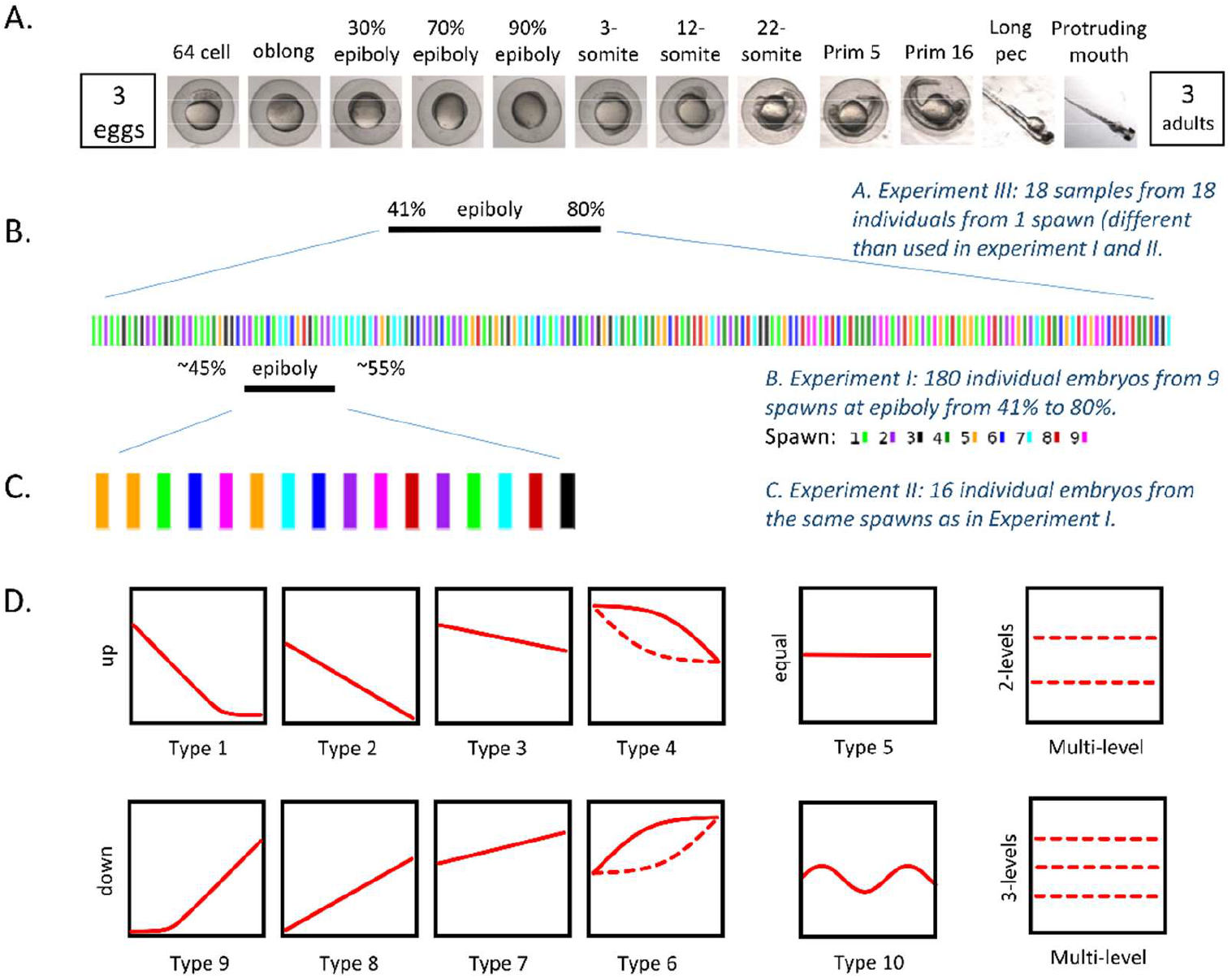
Graphical representation of the experiments. A. Small-RNAseq time course experiment over embryonic development from unfertilized egg to adult (male). Photos represent the individual embryos that were sequenced. B. High-resolution time course microarray experiment on 180 individual embryos from late blastula to early gastrula, which study is described in detail in [21]. C. Small-RNAseq experiment of individual embryos from the same spawns (except spawn 4) as in B, indicated by the same color legend. D. Schematic representation of the 10 gene-expression types from the high-resolution time course microarray experiment (B) plus multi-level gene expression types [21]. Not all multi-level genes show level expression; some also show an increasing or decreasing expression at one or more levels.

## Results and Discussion

### Experiment set-up

To investigate the factors that are involved in regulation of transcriptome dynamics during early zebrafish embryogenesis, we started our analyses from results of our gene-expression series in an extremely high temporal resolution on 180 individual zebrafish embryos [21], (Figure 1B). These embryos were obtained from nine different spawns, developmentally ordered and profiled from late blastula to mid-gastrula stage (approximately 40 - 80% epiboly). The individual profiles of the genes in this high-resolution time course can, together with the maternal-transcriptome results from another study of individual unfertilized zebrafish eggs [20], be queried through the Zebrafish Expression Browser (https://github.com/rauwerda/ZF_expression_browser). The transcriptome dynamics in the embryonic time course were captured in ten distinct gene-expression profiles, i.e. Types (Figure 1D). These ten types can be summarized in four categories of gene-expression dynamics: Down (Types 1-4), level (Type 5), UP (Types 6-9), and oscillating (Type 10), which leaves only one atypical type: Multi-Level (Figure 1D). Here we set out to link the observed gene-expression profiles to cellular factors that possibly can explain the observed transcriptome dynamics, such as chromosome location, TFBSs and miRNA expression. As an example of regulation of regulators, we investigated miRNA expression during the complete zebrafish embryogenesis by two additional small-RNAseq experiments (Figure 1A and 1C).

### Linking genome location and gene size to gene-expression types

The ten distinct gene-expression types of the time series (Figure 1 B and D) are based on the dynamic behavior of 6,734 Ensembl-defined expressed genes. Since chromosome organization, e.g. by nucleosome organization at promoters [22] plays a role in gene-expression regulation, we investigated the genome distribution of the associated genes per type (Supplemental File S1, pages 1-10). The fold enrichment of the number of genes of a specific gene-expression type on a chromosome, are distributed in a non-random fashion (Figure 2). Especially the fold enrichments of the Starting Type (Figure 1D, Type 9) and of the Discontinuously-Increasing Type (Type 6) show a distinct pattern on the genome and differ from the other types. Type 6 and Type 9 genes are, on chromosome 4 and chromosome 9 respectively, significantly overrepresented with a more than twofold enrichment (Supplemental File S2). We also plotted the position of the Multi-Level Type genes on the genome. The Multi-Level Type genes show an even wider distribution of fold enrichments on chromosomes as compared to the gene-expression Types 6 and 9 (Supplemental File S1, page 11), however no chromosome showed a significant overrepresentation of Multi-Level Type genes. Altogether, this means that, although the link appears to be particularly strong, chromosome location seems to be a factor that is involved in the transcriptome dynamics during early embryogenesis.

**Figure 2.**
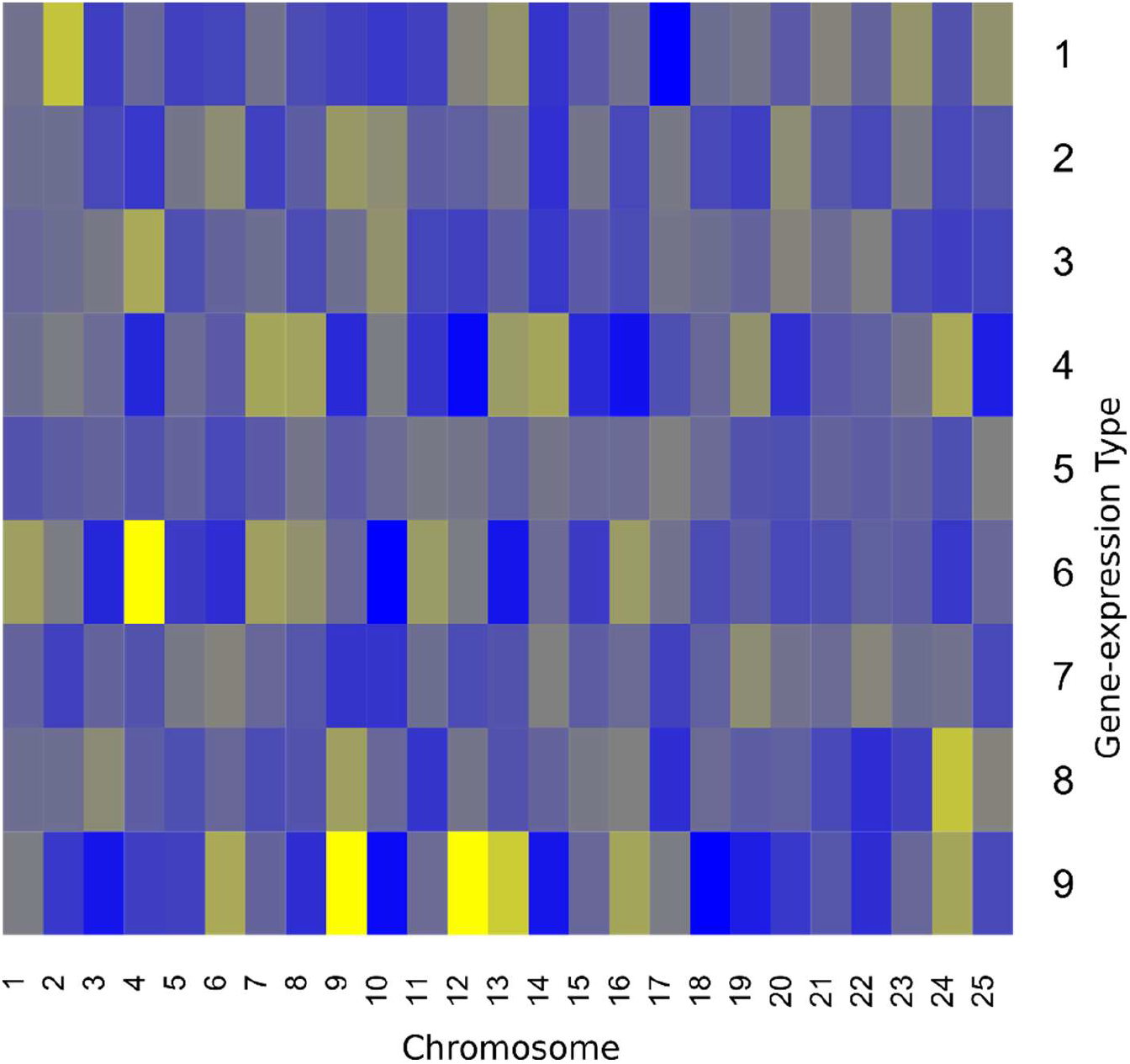
Relative number of genes in a gene-expression type per chromosome. Fold enrichment of number of genes of gene-expression types on chromosomes. The enrichment is calculated against all expressed genes.

In early embryogenesis, gene size is strongly related to gene expression [14]. In an attempt to link gene-size to gene-expression we found that gene-size distributions of expressed and non-expressed genes are different (Supplemental File S3). On average, expressed genes are shorter but very short genes of less than 4 kb are overrepresented in the non-expressed set. Comparing sizes of the genes in the Up, Level, and Down Categories with that of expressed genes, the Up and the Down Category distributions are significantly different: the Up Category genes are shorter while the Down Category genes are longer. This is in line with the fact that the earliest transcribed zygotic genes are relatively short [14]. It also suggests that the genes in the Down Category at least partly have a maternal origin. The size distribution of Multi-Level Type genes did not significantly differ from the size distribution of the present genes (results not shown).

### Linking promoter TFBSs to gene-expression types

TFs have an obvious role in gene-expression, hence we investigated the possible binding sites for TFs in the sets of expressed and non-expressed genes. TFBSs are typically located in the promoter, however defining promoters is a challenge and hence we arbitrarily took 300 bp regions upstream and downstream of the start site of each gene. Thus, we detected some remarkable patterns in our dataset. In the set of expressed genes, 307 TFBS motifs were found and in the set of non-expressed genes 165 (Supplemental File S4) were found. Both in the expressed and non-expressed gene sets, the number of TFBS motifs found in the upstream regions was larger than that in the downstream regions (respectively 1.9 and 1.4 times more). Three TFBS motifs that were enriched in the downstream regions of the non-expressed gene set were also enriched in the expressed genes upstream regions; two of them being POU domains (Pou2F2 and Pou3F3, for sequence logos, cf. Supplemental File S5). In the upstream analysis of the non-expressed gene set and the downstream analysis of the expressed gene set an overlap was found of 8 TFBS motifs, two of these belonging to the IRF-class (Irf6 and Irf4).

The overrepresented TFBS motif with the lowest p-value in the upstream analysis of the expressed gene set was the TATA-box (Tbp_primary). By considering the TFBS motif descriptions, some patterns appear (Table 1) that link homeobox domains (*Hox, Homez, Hlx* or *lmx*) and POU domains to gene expression in this analysis, whereas SRY (SOX) boxes and Zinc motifs relate to non-expression. One of the two uniquely enriched motifs in the non-expressed downstream set is Pou5f1::Sox2, a protein that is implicated in the maintenance of pluripotency, the activation of zygotic genes and in the regulation of miR-430, a miRNA responsible for the clearance of maternal mRNAs [23]. E2F TFs are involved in regulation of cell cycle progression [24].

**Table 1.** TFBSs in 300 bp regions upstream and downstream of the start sites of genes in the Expressed, Non-expressed, Down, Equal and Up Type gene sets. Percentages are given as the ration of TBFSs found in a set and number of TFBSs in the database (JASPAR 2014 and UniProbe Mouse).

| TFBS motifs |  |  |  |  |  | Expressed gene set |  |  |  |  |  |
| --- | --- | --- | --- | --- | --- | --- | --- | --- | --- | --- | --- |
|  |  | Expressed gene set |  | Non-expressed gene set |  | Down Type gene set |  | Equal Type gene set |  | Up Type gene set |  |
|  | JASPAR<br>2014 + UP | upstream | down-<br>stream | upstream | down-<br>stream | upstream | down-<br>stream | upstream | down-<br>stream | upstream | down-<br>stream |
| Homeobox domains | 52 | 92% | 25% | 0% | 0% | 23% | 21% | 0% | 0% | 56% | 0% |
| POU domain | 12 | 83% | 0% | 0% | 33% | 0% | 0% | 0% | 0% | 0% | 0% |
| Forkhead (FOX) boxes | 24 | 58% | 63% | 0% | 0% | 4% | 0% | 0% | 0% | 0% | 0% |
| SRY (SOX) boxes | 33 | 18% | 9% | 55% | 3% | 6% | 0% | 0% | 0% | 0% | 0% |
| E2F | 8 | 75% | 88% | 13% | 0% | 0% | 0% | 0% | 0% | 0% | 0% |
| Zinc related motifs | top 20 sign. | 10% | 15% | 65% | 70% | 5% | 0% | 0% | 0% | 30% | 0% |

We also tested the gene-expression categories; Down, Level, Up, and Type Multi-Level for TFBSs (Table 1). The absence of any overrepresented binding motifs in the Type 5 set probably implies that the genes in this set are not one group and thus are regulated by many different factors. Most (56%) of the overrepresented TFBSs were found upstream of the Up Category genes, which is understandable given that they all change expression in the examined embryonic developmental period. The overrepresented motifs in the gene sets with increasing gene expression are in majority zinc finger motifs. In the Up-Category 5 TFAP motifs and 10 zinc finger motifs were found.

The 11 TFBSs found in the downstream sequences of the Down Category only and explicitly overlap with TFBSs in the non-expressed genes. The TFBS set contains 4 MEIS homeoboxes, 2 PBX homeoboxes and 2 TALE family homeoboxes (TGFB) with remarkably low p-values. These 4 MEIS boxes, together with 1 PBX box and 2 TALE boxes are also overrepresented in the downstream sequences of the gene-expression Type 1 set, while 1 MEIS box is overrepresented in the Type 2 set as well. Since Types 1 and 2 contain genes which expression actually stops during the investigated embryogenesis period, our data indicate that processes involving Meis are apparently switched off. The Meis cofactor has been reported to bind directly to Pbx, and to displace histone deacetylases from Hoxb1 regulated promoters in zebrafish embryos [25].

Noticeably, in the set of Multi-Level Type genes, we did not find any overrepresented TFBS motifs.

### Linking 3’UTR polyadenylation motifs to gene-expression types

At the late blastula stage in zebrafish embryogenesis, which coincides with the start of our high-resolution time course (Figure 1B), there is extensive zygotic transcription. Whether at the early gastrula stage all maternal mRNA has been cleared, is still a question. Maternal mRNA can be polyadenylated in the cytoplasm, a process that uses polyadenylation motifs in the 3’ UTR of transcripts [12,18]; hexamer motifs (hex-1 and hex-2) form the actual polyadenylation signal and determine the actual start of the poly(A)-tail; cytoplasmic polyadenylation elements (eCPE, CPE-1, CPE-2, and CPE-3) can bind to the CPE-binding protein and to other factors and not only support polyadenylation but also act as regulatory elements in cytoplasmic polyadenylation as well[12]; and Pumilio binding elements (PBEs) facilitate the binding of the Pumilio protein, which in its turn can stabilize the binding of CPEB [26]. We analyzed overrepresentation of such motifs in the geneexpression categories and types (Supplemental File S6). In the Expressed genes all motifs but one hexamer (hex-2) and the Pumilio Binding Element are highly overrepresented. Within the expressed genes the hex-2 motif is overrepresented in the Type 2 set. Furthermore, two motifs are overrepresented in the Type 5 set, among which is the embryonic cytoplasmic polyadenylation element [27]. The hex-1 motif is overrepresented in the Up Type, which is largely due to its highly significant overrepresentation in Type 7 genes. These results indicate that a process in which maternal mRNAs are reused via cytoplasmic polyadenylation cannot be ruled out.

### Linking pathways to gene-expression types

Genes operate in cellular pathways. Hence pathway analysis is a common approach in omics experiment analysis. Here, we tested which biologically important sets, such as KEGG pathways were overrepresented in any of the gene-expression types and categories (Table 2). The top five KEGG pathways that are overrepresented in the Expressed gene set are also overrepresented in the gene-expression Types 5 and 7. These two types are characterized by an *expressed* status throughout the studied embryogenic period. Other overrepresented pathway categories in the Types 5, 6 and 7 also point at transcriptional, translational processes, whereas the starting Types 8 and 9 show morphogenic terms and an involvement of the homeobox domain (Supplemental File S7). In the Down Category gene-set, a number of signaling pathways are overrepresented that are also overrepresented in the Non-Expressed gene set. Remarkably, also some embryonic developmental gene ontology terms are overrepresented in the gene-expression Type 4. Three pathways from the expressed set are also overrepresented in the Down Category, but with a lower rank than the pathways in the Level and Down Category. With three genes, the iron-sulfur cluster binding set was found to be the only overrepresented category in Multi-Level Type genes. The fact that in such analysis mostly high-level pathways are identified is because genes in pathways are not expressed at the same time and in the same matter. Hence, genes of a pathway are likely to be scattered over the gene expression types and categories, preventing their discovery by this naïve analysis.

**Table 2.** Overrepresented KEGG Pathways in the gene-expression categories. The gene-expression types are organized in three categories: Down (Types 1-4), Level (Type 5) and Up (Types 6-9) Categories (Figure 1D) and overrepresentation is determined (p-value) against the background of the expressed genes. Shown are the p-values of the overrepresentation of the Down, Level and Up Categories. Geneexpression types that are overrepresented individually are indicated by their type number between brackets. *; no overrepresentation was found in any category, but that a gene-expression type was overrepresented individually. In that case p-values are shown in italics and represent the overrepresented gene-expression type. The columns marked with e. and *n.e.* respectively indicate the rank by which a pathway is overrepresented in the expressed genes and non-expressed genes. Ranks are called by p-values, without a fold-enrichment criterion.

| Pathway | Down | Level | Up | e. | n.e. |
| --- | --- | --- | --- | --- | --- |
| dre04010:MAPK signaling pathway | 5.42E-03 (1) |  |  | - | 6 |
| dre04622:RIG-I-like receptor signaling pathway | 2.28E-02 |  |  | - | - |
| dre04070:Phosphatidylinositol signaling system | 2.28E-02 |  |  | - | 8 |
| dre04144:Endocytosis | 2.47E-02 (2) |  |  | - | - |
| dre03018:RNA degradation | 3.37E-02 |  |  | 8 | - |
| dre04621:NOD-like receptor signaling pathway | 3.66E-02 |  |  | - | - |
| dre04620:Toll-like receptor signaling pathway | 3.98E-02 (2) |  |  | - | 10 |
| dre00071:Fatty acid metabolism* | 2.73E-02 (2) |  |  | 20 | - |
| dre04114:Oocyte meiosis* | 9.78E-03 (3) |  |  | 14 | - |
| dre03040:Spliceosome |  | 2.84E-09 (5) |  | 1 | - |
| dre00280:Valine, leucine and isoleucine degradation |  | 3.47E-03 (5) |  | 15 | - |
| dre00640:Propanoate metabolism |  | 1.09E-02 (5) |  | 19 | - |
| dre04110:Cell cycle |  | 2.05E-02 (5) |  | 3 | - |
| dre04144:Endocytosis |  | 4.57E-02 (5) |  | - | - |
| dre03010:Ribosome |  |  | 5.36E-29 (7) | 2 | - |
| dre00190:Oxidative phosphorylation |  |  | 1.38E-03 (7) | 4 | - |
| dre03050:Proteasome |  |  | 1.11E-02 (7) | 5 | - |
| dre00260:Glycine, serine and threonine metabolism* |  |  | 1.14E-02 (9) | - | - |
| dre04260:Cardiac muscle contraction* |  |  | 1.90E-02 (7) | - | - |
| dre04512:ECM-receptor interaction* |  |  | 3.83E-02 (6) | - | 5 |

### Linking 3’UTR miRNA target sites to gene-expression types

Over the last decades it has become clear that non-coding small-RNAs, such as miRNAs play an important role in gene-expression regulation. We therefore examined possible regulation of genes in the gene-expression types by miRNAs by estimating the overrepresentation of miRNA target sites in the 3’UTRs of the expressed genes and of the genes in the gene-expression types (Table 3, Supplemental File S8). The gene sets in the Up Category do not have overrepresented miRNA targets, neither do the sets in the Down Category have any underrepresented targets. There is a large overlap in target sites between the Up Category underrepresented set and the Down Category overrepresented set, although miR-128-3p, miR-20a-3p and miR-20a-5p are specific for the Up Category underrepresented set and miR-181a-2-3p, miR-124, miR-16a, miR-228 and miR-150 are specific for the overrepresented targets in the Down Category genes. No overrepresented target sites were found for the Level Category and Multi-Level Type genes. The observations that no miRNAs are expressed that target genes with increasing expression and that genes with a deceasing expression have miRNA targets of actually expressed miRNAs are in line with the general miRNA mechanism. MiR-430c-3p, miR-19a-3p, miR-19a-5p and miR-430c-5p are the 1^st^, 6^th^, 7^th^ and 8^th^ most expressed miRNAs and target genes in the Down Category. However, other highly abundant miRNAs, such as miR-430b and miR-20a do not specifically target a gene-expression category nor type.

**Table 3.** Number of miRNA targets [18] that are overrepresented or underrepresented in the set of expressed genes and in the types of the high-resolution time course experiment [21] and the intersect with these targets and the actually expressed miRNAs in embryos taken from the same spawns as in [21] at approximately 50% epiboly (fig. 1 C). The types are organized in three categories: Down (Types 1 to 4), Level (Type 5) and Up (Types 6 to 9). Overrepresentation for the expressed genes is calculated against the background of all queried genes and analysis of the genes in the gene-expression types is performed against the background of the expressed genes. Since TargetScanFish, from which the targets were retrieved often does not discriminate between the 3p and 5p mature miRNA forms, the number of corresponding occurrences of expressed miRNAs and targets is indicated between brackets.

| Class/Category | Number of targets<br>($p_{\text{corrected}} < 0.15$) | Intersect of targets and expressed miRNAs | Intersecting expressed miRNAs |
| --- | --- | --- | --- |
| Expressed | 28 | 5 | miR-124, miR-462, miR-23a, miR-365, let-7a |
| Non-expressed | 10 | 8 (5) | miR-93, miR-430a-3p, miR-430a-5p, miR-733, miR-430c-3p, miR-430c-5p, miR-430b-3p, miR430b-5p |
| Up *(overrepresented)* | 0 | 0 |  |
| Up *(underrepresented)* | 26 | 13 (8) | miR19a-3p, miR19a-5p, miR-128-3p, miR-148, miR-430c-3p, miR-430c-5p, miR-122, miR-16a, miR-20a-3p, miR-20a-5p, miR-17a-3p, miR-17a-2-3p, miR-17a-5p |
| Level *(overrepresented)* | 0 | 0 |  |
| Level *(underrepresented)* | 0 | 0 |  |
| Down *(overrepresented)* | 20 | 14 (10) | miR-17a-3p, miR-17a-2-3p, miR-17a-5p, miR-148, miR-19a-3p, miR-19a-5p, miR-181a-2-3p, miR-124-3p, miR-122, miR-16a, miR-228, miR-150, miR-430c-3p, miR-430c-5p |
| Down *(underrepresented)* | 0 | 0 |  |
| Multi-Level *(overrepresented)* | 0 | 0 |  |
| Multi-Level *(underrepresented)* | 0 | 0 |  |

### Analyzing miRNAs in zebrafish embryogenesis

Although using miRNA target-sites in 3’ UTRs we were unable to establish a strong link between our gene-expression categories or types and miRNAs, it is obvious that miRNAs are an important factor in gene-expression regulation. To estimate the value of the results of the overrepresentation, we checked which miRNAs are actually present during early gastrulation by a small-RNAseq experiment of 16 individual zebrafish embryos that originated from the same spawns as the high-resolution time course and are at around 50% epiboly (Figure 1C and Supplemental File S9). Of the overrepresented miRNA targets in the expressed set, a sobering less than 20% show actual expression of the associated miRNA in early gastrulation, whereas the five most abundantly expressed miRNAs, miR-430c-3p (1), miR-430b-3p (2), miR-430a-3p (3), miR430b-5p (4), miR-430a-5p (5) correspond with overrepresented targets in the non-expressed set.

Further analysis revealed that the correlation of total number of miRNA reads and the relative amount of synthetic spike-in reads of 0.79 does not suggest big differences in the absolute amount of expressed miRNAs between these staged embryos. Moreover, we found 11 miRNAs with over 5,000 reads in all 16 samples together (high-abundant) and 84 miRNAs with less than a total of 5,000 reads, but more than 50 reads in at least one sample (low-abundant, Supplemental File S10). Remarkably, the top 5 miRNAs accounts for 91% of all miRNA, whereas the top 11 for 96%. We analysed the high- and low-abundant miRNAs separately (Figure 3, Supplemental File S10). In our previous high-resolution time series experiment, we observed that the gene-expression of genes in the Multi-Level Type correlated with the spawn from which the embryos originated. We questioned whether miRNAs would also show a spawn-specific expression effect. In the high-abundant miRNA set only the samples from spawn 7 cluster together and in the low-abundant miRNA set this is the case for spawn 7 and spawn 1. Hence, in contrast to mRNA expression, spawn seems no major determinant in relative miRNA expression.

**Figure 3.**
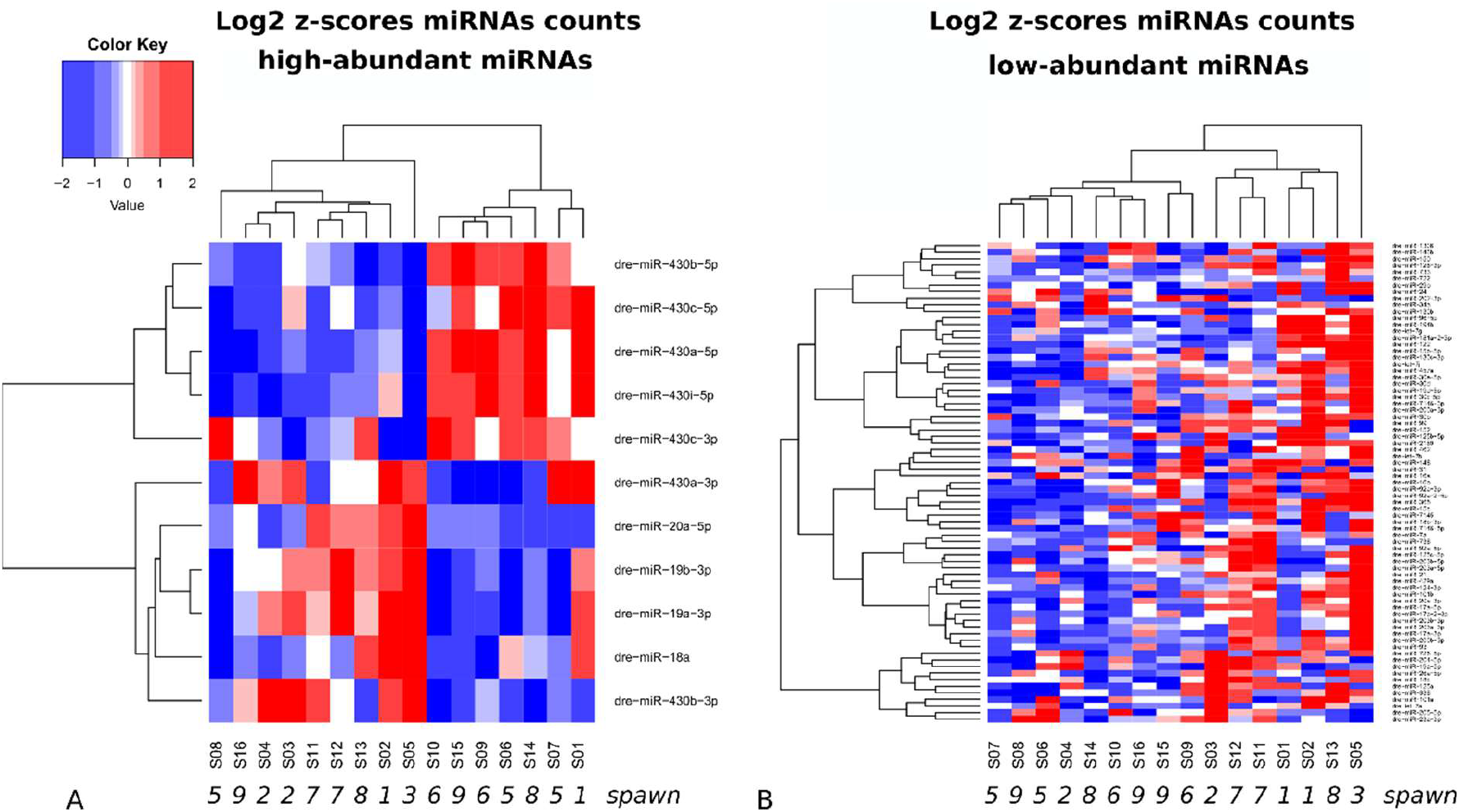
Heatmaps of high- and low abundant miRNAs. Heatmaps of scaled and centered log2 miRNA counts. The spawn number is indicated below the sample-id. A. Heatmap of high-abundant miRNAs with a total read count > 5,000. B. Heatmap of low-abundant miRNAs with a read count of at least 50 in one of the samples but with less than 5,000 reads in all samples.

To gain a better insight in the regulation of miRNAs during the embryogenesis, we additionally sequenced small-RNA along the entire zebrafish embryonic development (Figure 1A). In contrast to the small RNA samples taken at early gastrulation, over embryonic development the correlation of the small RNA read counts with the amount of synthetic spike-in reads is low, i.e. 0.24. Also, the number of miRNA reads in the 64-cell stage is substantially lower than in later stages (Supplemental File S11). This indicates that the absolute amount of miRNA is changing during embryogenesis, an observation that was also made by others [28,29]. We therefore used the relative abundances of miRNAs to characterize miRNA expression over embryogenesis. To evaluate how miRNAs are relatively expressed during embryogenesis, we partitioned the developmental time in seven stages (egg, maternal, gastrulation, somite, pre-hatching, post-hatching and adult) (Figure 4H). It is clear that there are several distinct miRNA expression profiles during embryogenesis, which shows an extensive regulation of the expression of these miRNA during embryogenesis. As these expression profiles are indicative for their functioning in embryogenesis, they can be used to elucidate the roles of these miRNAs. The miR-430 serves as an example, being involved in maternal mRNA degradation and being expressed with a peak at the gastrulation (Figure 4C).

**Figure 4.**
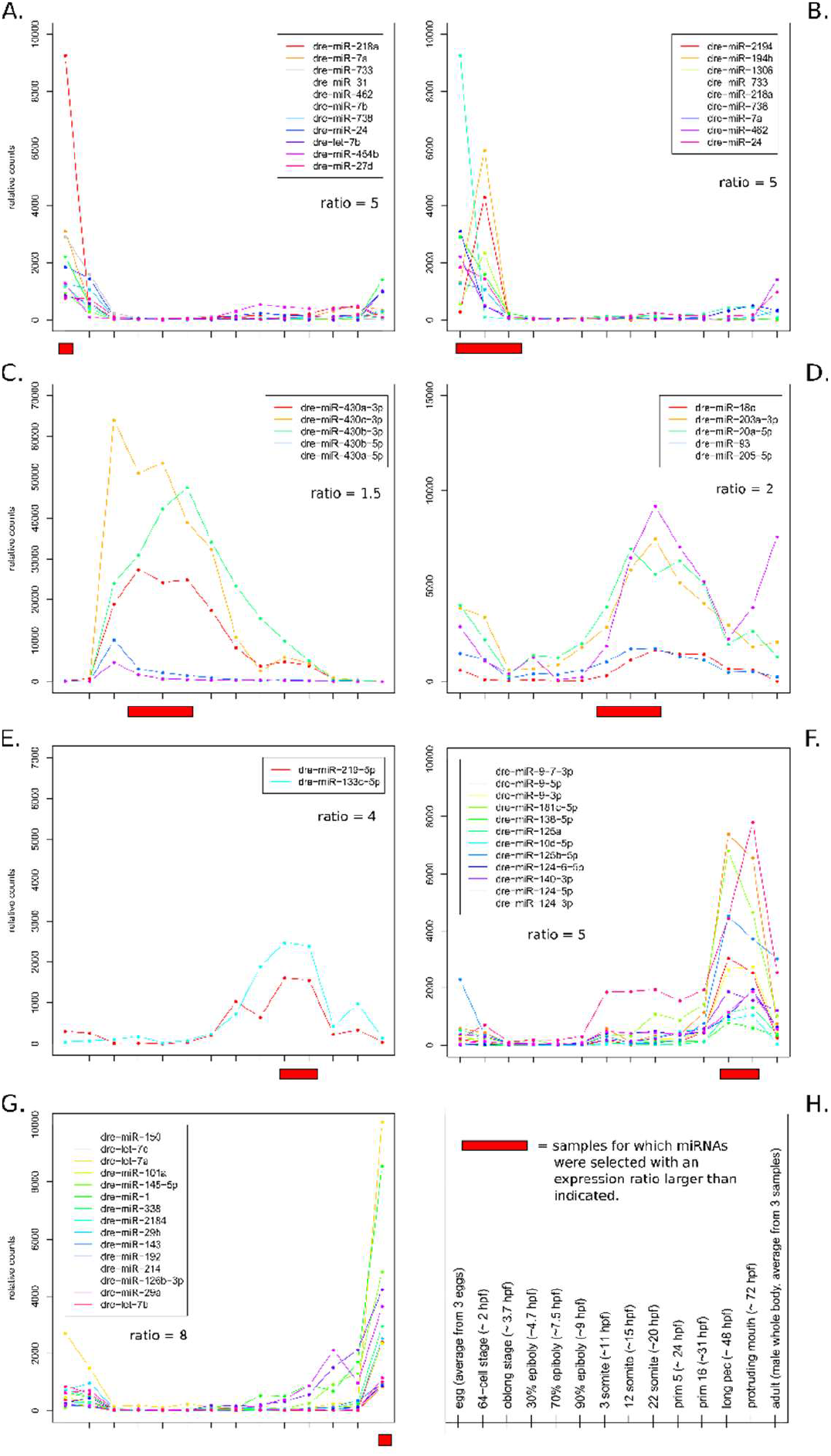
Relative miRNA abundance in zebrafish development. Relative counts of miRNA abundances in an embryonic developmental time course. The X-axis represents developmental stage as specified in panel H. The Y-axis represents relative counts. A-G miRNAs specific for a developmental stage. All miRNAs have a relative abundance of at least 700 (approximately 0.5% of all mapped miRNA reads) and are present in one or more specific stages relative to all other stages at a ratio indicated in the figure. A. Egg; B. Maternal: egg, 64 cell + oblong stage; C. Late Blastula + gastrula: 30% epiboly + 70% epiboly + 90% epiboly; D. Somite: 3-somite + 12-somite + 22-somite; E. Pre-hatching: prim-5 + prim-16; F. Post-hatching: Long-pec + protruding mouth; G. Adult (male). H. Legend of the X-axis.

Pushing forward, we analysed the 11 high-abundant miRNAs that we previously identified at early gastrulation more extensively. These miRNAs display a wave-like expression pattern over the entire embryogenesis (Figure 5B); some that are maternally deposited show an increase in relative abundance by late gastrula stage and have a decreased relative abundance in adult tissue. Others are zygotically expressed from the oblong stage onward, with relative abundance peaking at different stages. Yet in the early gastrulation experiment (Figure 1C), two subsets of miRNAs could be identified along with two groups of samples, that showed an opposite behaviour (Figure 3A). Upon closer inspection, it became clear that, at 50% epiboly, whenever dre-miR-430c-3p is expressed at a higher level, dre-miR-430b-3p is expressed at a lower level (correlation −0.90, Figure 5A). These miRNAs with this striking anti-correlating expression in the gastrulation show, measured from oblong to 90% epiboly, a decreasing and an increasing relative abundance, respectively (Figure 5B). It therefore is possible that the anti-correlating behaviour originates from the fact that the analysed embryos were in a slightly different embryonic stage.

**Figure 5.**
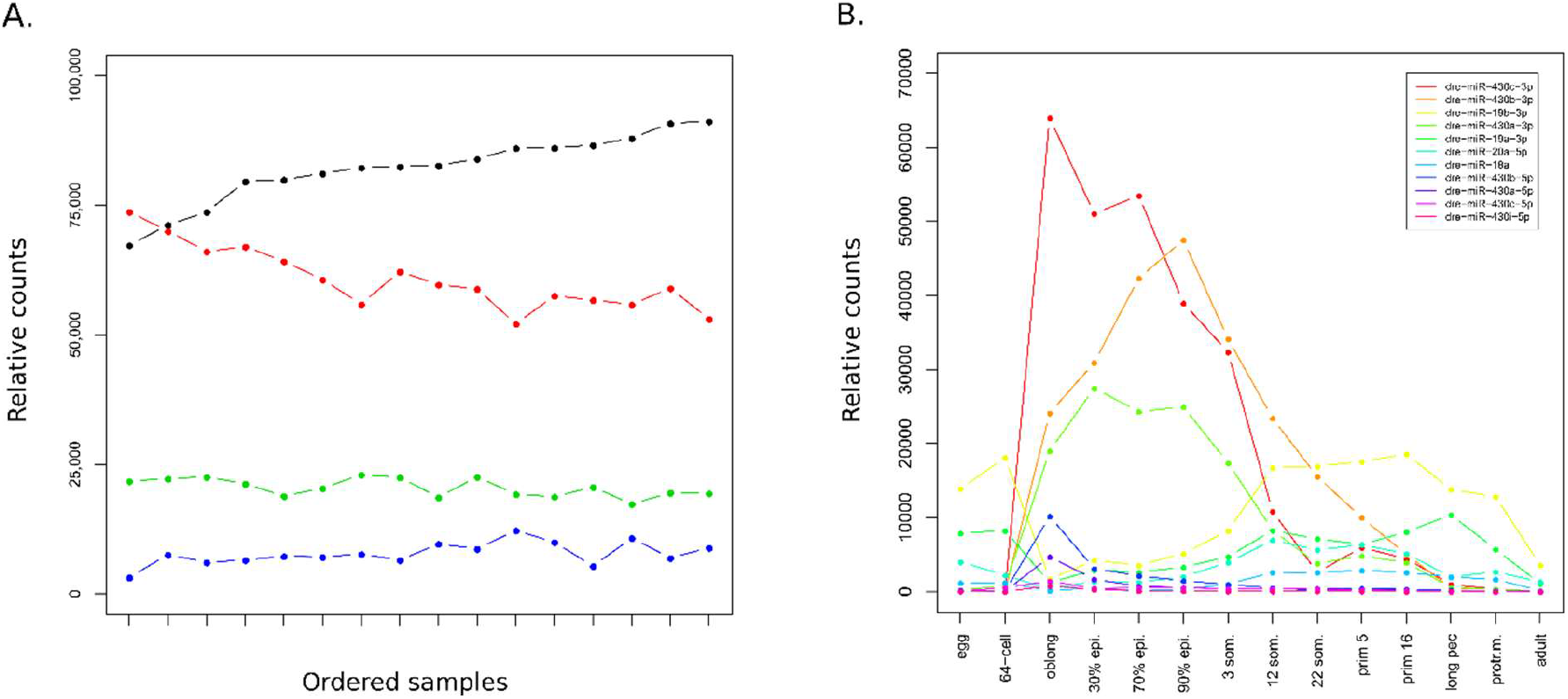
Comparing miRNA expression. Results from two early gastrulation small-RNAseq experiments (Figure 1C and A). A. Expression of the four most high-abundant miRNAs (dre-miR-430c-3p, black; dre-miR-430b-3p, red; dre-miR-430a-3p green; and dre-miR-430b-5p, blue) in the early-gastrulation experiment (Figure 1C) with the samples ordered by dre-miR-430c-3p. B. Relative abundances of the top-11 miRNAs during embryonic development as measured in the whole embryogenesis time course (Figure 1A).

To a lesser extent, dre-miR-430a-3p and dre-miR-430b-5p also show anti-correlated expression (correlation −0.6). In total, we found in the combined high- and low-abundant (n=95) miRNAs, eight sets with a correlation higher than 0.9 and two sets of miRNAs with a correlation lower than −0.85 (Supplemental File S12). A correlation of −0.85 was found between dre-miR-430b-5p and dre-miR-17a-5p, which are the 4^th^ and 21^th^ most abundantly expressed miRNAs. A decreasing, respectively increasing relative expression similar to dre-miR-430c-3p and dre-miR-430b-3p pair is observed with the anti-correlating pair miR-430b-5p and miR-17a-5p (data not shown). While the genomic clusters of miR-430a and miR-430b lie on chromosome 4 at around 28 Mbase, with many variants (e.g. miR-430a-10.1, miR-430b-20.1 and miR-430a-17.2) interspersed with one another, miR-17a has two variants, one on chromosome 1 and on chromosome 9. So chromosome location alone does not explain this strong correlation. Collectively, it shows that the expression of mRNA-expression regulating miRNAs is quite strictly regulated.

## Concluding Remarks

The embryonic transcriptome is tightly regulated during development. This is not only established for pools of embryos [17], but is also true when individual embryos are compared to one another over time. Here we analysed a number of cellular factors that might be involved in this regulation and found that all gene-expression regulation strategies studied here are used by the developing zebrafish embryo. The picture that emerges is that a large number of different regulatory events take place in the short developmental time from late blastula to gastrula. For instance, the overrepresented motifs in the Up Category were almost exclusively found in genes of that category, but these motifs are only found in the promoters of a subset of these genes; the most highly overrepresented motif in this category, ESR1 was found in 18 % of the genes of this category (data not shown, cf. Material & Methods). Hence ESR1 is a cis element that is important in the upregulation of genes during this embryonic phase, but it only one of the many factors involved in the up-regulation. Another example that illustrates the diversity of regulatory mechanisms that is at work is the co-occurrence of miR-430 as the predominant miRNA in the blastula and gastrula stages and the presence of cytoplasmic polyadenylation motifs in many expressed genes. MiR-430 is widely implicated in the clearance of the maternal message [16] while the presence of cytoplasmic polyadenylation motifs still keeps open the possibility for the embryo to activate and use maternally provided transcripts. Looking at predominant pathways and Gene Ontology categories we see that a number of these sets are overrepresented in the gene-expression types; however, the percentage of genes that is called to be present in a set is with 19% highest in the Type 6 for GO:0045449 (regulation of transcription), but in general these percentages are below 5% (Supplemental File S7). Rather than to characterize early gastrulation as to be regulated by a handful of mechanisms, we observe an overwhelming amount of processes involved in regulation of this developmental stage.

Not only the resulting gene expression is tightly regulated, also at least some of the regulators themselves are tightly controlled. This is illustrated by the anti-correlating expression of miR-430c-3p and miR-430b-3p (Figure 5A), but also by the wave-like expression cascade of the different members of the miR-430 family Figure 5B).

We conclude that genes in gene-expression types are active in specific pathways. These temporal expression profiles might also be referred to as synexpression groups, a term coined by Niehrs and Pollet [30] and recently again used to characterize the embryonic transcriptome [17]. Compared with the latter study our study zooms in on a much shorter period and has an even higher sampling density. But also in this shorter developmental period of 3 hours, the underlying regulation of the strict temporal behaviour of the transcripts in the Up, Level and Down Categories, is extremely complex on the level of the individual transcriptome. Novel approaches in single cell analysis [31], RNA tomography [32] and RNA tracking in live cells [33] will add to the resolution in time and space at which the developing embryo can be studied in the future.

## Materials and Methods

### High-resolution time course experiment late blastula to mid-gastrula

The high-resolution time course experiment in early zebrafish embryogenesis and specifically the handling of the zebrafish embryos, image processing, RNA extraction, amplification & labeling, hybridization, scanning, data preprocessing & normalization, determination of the sample order, gene categorization, identification of starting and stopping points and identification of multi-level genes is described in [21]. 3’ UTR sequences, sequences upstream of genes and gene locations were retrieved using Biomart Zv9 (Ensembl release 79). From the same experiment 16 previously not used samples from 8 spawns were taken (Supplemental File S11) and their small RNA content was sequenced using the procedure described below. The 16 embryos were in early gastrulation with an epiboly ranging from 43% to 58%.

### Embryonic developmental course

A number of samples were taken along the zebrafish’ development (Supplemental File S11). Zebrafish were handled and maintained according to standard protocols (http://ZFIN.org). The local animal welfare committee (DEC) of the University of Leiden, the Netherlands specifically approved this study. All protocols adhered to the international guidelines specified by the EU Animal Protection Directive 86/609/EEC. Embryos were obtained by placing a female and a male, both of genotype ABTL, overnight in a tank separated by transparent, but watertight division. Approximately 1 hour after the start of the light period the division was removed after which spawning and fertilization took place, generally in one or more pushes. Embryos were retrieved by transferring both animals to a different tank and sieving the remaining water. Embryos, which were obtained from one spawn were maintained at 28.5 °C in egg water (60 μg/ml Instant Ocean sea salts). The embryos were processed, from approx. 10 minutes’ post fertilization (hpf) to 72 hpf. During this period, embryos were, while kept at a temperature of 28.5 °C, one by one taken out of the medium and positioned along the anteroposterior axis under a stereo microscope (Leica MZ16 FA) in a 1.5% agarose gel (Sigma Aldrich IX-A) composed with egg water. After an image was taken the embryo was transferred to a 1.5 ml tube and was snap-frozen in liquid Nitrogen. Unfertilized eggs (oocytes) were collected by following a similar approach as described above. However, approximately 1 hour after the start of the light period the female now was removed from the tank and anesthetized briefly in another tank with egg water containing 0.02% buffered ethyl 3-aminobenzoate methane sulfonate (Tricaine, Sigma-Aldrich). Subsequently, the eggs were obtained by gently squeezing the female. Whole body male-adult zebrafish samples and eggs were flash-frozen in liquid nitrogen and stored at −80°C. Before freezing, fish were put under anesthesia using 0.02% buffered Tricaine.

### Small-RNA isolation

Tubes with single zebrafish embryos were kept in liquid nitrogen and processed using a pre-chilled metal micro-pestle (Carl Roth) and Qiazol (Qiagen) as described by de Jong et al 2010 [34]. Synthetic spike-ins [35] were added to the Qiazol (equal amounts for each single embryo) and subsequently the phase separation was performed according to the manufacturer’s instructions. For the samples in the time course experiment (Figure 1A) the aqueous phase was precipitated to first isolate total RNA, followed by an enrichment of small RNAs using the mirVana miRNA Isolation Kit (Thermo Fisher Scientific). For the samples taken at gastrulation (Figure 1C) the aqueous phase was directly applied to a mirVana spin-column and further processed according to the manufacturer’s instructions. Small RNA concentration and quality were assessed on a NanoDrop ND-2000 (Thermo Fisher Scientific) and 2200 TapeStation System with Agilent RNA ScreenTapes (Agilent Technologies), respectively.

### Small-RNA-sequencing

Bar-coded small RNA sequencing libraries were generated using the Ion Total RNA-Seq Kit v2 and the Ion Xpress RNA-Seq bar coding kit (Thermo Fisher Scientific). For the developmental course samples (Figure 1A), the two-step bead-based size selection that directly follows reverse transcription and cDNA amplification was replaced with a single bead-based purification step using ethanol at a final concentration of 45% and 40% ethanol, respectively. Together, these modifications increase the size limitation of the small RNAs that will be part of the small RNA sequencing library from below ~50 to below ~200 nucleotides. Size distribution and yield of the resulting barcoded libraries were assessed using the 2200 TapeStation System with Agilent D1K ScreenTapes (Agilent Technologies). Libraries were prepared for sequencing on the Ion Chef System (Thermo Fisher Scientific). Sequencing was performed on an Ion Proton System using Ion PI v3 chips (Thermo Fisher Scientific).

### Genome location and gene size

The visualization of genes on the assembly was done using the ‘View karyotype’ link on the Ensembl website. The fold enrichment of the number of genes of a gene-expression type *i* on chromosome *j* was calculated according to the following formula:

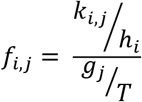

with:

g_x_ = number of expressed genes on chromosome x
h_y_ = number of genes of Type y
k_xy_ = number expressed genes on chromosome x of Type y
T = total number of expressed genes

Testing the similarity of distribution of gene sizes was performed with the implementation of the Kolmogorov Smirnov test in R (kstest).

### Enrichment analysis of TFBSs

Promoters were retrieved as 300 nucleotides upstream and downstream stretches of DNA sequence relative to the transcription start site of a gene. These sequences were tested against 591 motifs in the JASPAR Vertebrates 2016 [36] and the UniPROBE Mouse [37] databases using AME version 4.11.2, which is part of the MEME suite [38]. The background model was established once using all upstream and downstream sequences of genes that are interrogated by the microarray [21]. A one-tailed Wilcoxon rank-sum test was applied on the average odds motif score, after which a correction for the multiple testing of 591 motifs was applied using a Bonferroni procedure. A corrected p-value of 0.05 was used. AME was used with a fixed partition: the non-expressed set was tested with the expressed set as control sequences, the expressed set was tested with the non-expressed set as control sequences and all other sets were tested with the sequences from that set with the remainder of the sequences from the expressed set as control sequences. The number of motifs occurring on a set of promoter sequences was established with the FIMO tool version 4.11.2 [39], which is part of the MEME suite, with a single match-p value cut off of < 0.001.

### Enrichment analysis of Polyadenylation motifs

Hexamer motifs, poly(A) sites (PASs) and Pumilio Binding elements (PBEs) in the 3’ UTRs were determined using the AME tool version 4.11.2 of the MEME suite [38]. The sequence motifs were retrieved from [19,27,40,41]. The background model was established once using the sequences of genes that are interrogated by the microarray [21]. A one-tailed Wilcoxon rank-sum test was applied on the average odds motif score, after which a correction for the multiple testing of 16 motifs was applied using a Bonferroni procedure. A corrected p-value of 0.05 was used. AME was used with a fixed partition and all sets were tested in a similar fashion as with the TFBS analysis.

### Set analysis

Overrepresentation analysis on pathways and Gene Ontology categories was carried out using the web version of DAVID Bioinformatics Resources 6.7 [42]. Backgrounds were specified by choosing an appropriate subset of all genes that are queried by the microarray. The criteria to call a category (i.e. GO category, chromosome, KEGG pathway) as overrepresented are an EASE-score < 0.05 and a fold enrichment (calculated in a similar fashion as before) larger than 1.5.

### miRNA target site analysis

The Ensembl genes that have putative targets of miRNAs in their 3’ UTRs were obtained from TargetScanFish, release 6.2 [18], using those targets with context scores smaller than −0.2. The probability of a found overlap of a gene set containing the targets for a specific miRNA in a set of interest was calculated using a hypergeometric test in R by using an appropriate background (cf. Results and Discussion). A correction for multiple testing was applied using the Benjamini Hochberg procedure from the multtest R-library [43].

### miRNA mapping, normalization and expression analysis

Reads were mapped to zebrafish miRNA hairpin and mature sequences from miRBase version 21 [44] by the following procedure; all reads larger than 14 nucleotides but shorter than 44 nucleotides were mapped against the annotated mature miRNAs augmented with 3 nucleotides on the 5’ and 3’ end, while allowing for 10% mismatches or indels. For the mature miRNAs that lack either a 5p or 3p counterpart, reads were mapped against the half of the premature hairpin opposite to the half that contains the annotated mature miRNA. These mappings were annotated as ‘putative mature miRNAs’. Since a number of our analyses uses the miRNAs that are defined by TargetScanFish, miRBase annotations were condensed into the annotations ofTargetScanFish by aggregation of the miRBase identifiers that point to an identical mature miRNA.

A normalization factor was estimated using the assumption of an equal amount of small-RNA molecules in all samples. This factor was checked using the read counts of the synthetic spike-ins [35]. However, since the assumption that the number of small-RNA molecules is the same throughout development is evidently not correct [28,45], we used relative counts in our analyses. A relative count is defined as the ratio of reads mapping in a sample to a miRNA and the total number of reads in that sample mapping to miRNAs times the average number of reads mapping to miRNAs in the entire experiment. Clustering of relative counts was performed on log transformed counts that were subsequently scaled and centered over all samples. Heatmaps were drawn with the heatmap.2 function in R using ward.D2 as clustering method and using a Euclidean distance.

## Data availability

All experimental data sets are available online. High resolution gene expression profiles from late blastula to mid-gastrula stage (Figure 1B): accession number GSE83395 (http://www.ncbi.nlm.nih.gov/geo/query/acc.cgi?acc=GSE83395). Sequencing data are deposited at (www.ncbi.nlm.nih.gov/bioproject), small-RNA-seq data developmental time course (Figure 1A): PRJNA347637; small-RNA-Seq data at gastrulation (Figure 1C): PRJNA350377

## Acknowledgments

The University of Amsterdam has provided the funding for this work, but had no role as such in the study design, data collection and analysis, decision to publish, or preparation of the manuscript. We would like to thank Prof. Dr. Annemarie Meijer and Dr. Marcel Schaaf for their hospitality and kind assistance in obtaining the zebrafish samples at the Leiden University.

## Supplemental file descriptions

### Supplemental_File_S1.pdf

Chromosomal location of expressed genes. Page 1 displays the number of expressed genes on each chromosome. The pages 2-10 display per page the position of these genes in a gene-expression type on the genome. The Types 1 to 9 are shown. Page 11 displays the position of the Multi-Level Type genes on the genome. Page 12 shows the distributions of the fold enrichment of the number of genes per type on a chromosome.

### Supplemental_File_S2.xlsx

Overrepresentation of the gene-expression types, of the Down and Up Category and of the Multi-Level Type genes. Shown are the following categories: *General:* Chromosome, *Gene ontology:* Biological Process, Molecular Function & Cellular Compartment, *Functional Categories:* Uniprot Sequence Feature, PIR (Protein Information Resource), *Protein Domains:* Interpro, SMART, *Pathways:* KEGG, *Tissue:* Uniprot Tissue. Each tab shows the set for which overrepresentation was tested and displays all categories that have an EASE value (indicated by *PValue*) smaller than 0.05 and a fold enrichment larger than 1.5.

### Supplemental_File_S3.xlsx

Tab A. Table of average gene size in expressed and in the Up, Level and Down Categories compared with the average gene size of the remaining genes as indicated in the table. The p-value indicates probability that the gene sizes in the comparison are drawn from the same distribution. Tab B. Visualization of the distributions of gene size in the expressed, Up, Level and Down gene-expression Categories.

### Supplemental_File_S4.xlsx

Enriched motifs in stretches of DNA 300 nt upstream and downstream of the transcription start site. From the 4^th^ column onward the overrepresented motifs from JASPAR2014 and Uniprobe Mouse are given for each gene-expression type, first by a column enumerating the possible motifs in the upstream sequences and then by a column indicating the possible motifs in the downstream sequences. All values are ordered by corrected p-value from low to high, unless a motif already occurred in a previous column, in which case the p-values are unordered.

### Supplemental_File_S5.pdf

All enriched motifs per gene-expression type, p-values, corrected p-values and sequence logos. Gene-expression types without any enriched motifs are omitted.

### Supplemental_File_S6.xlsx

Overrepresentation of hexamer motifs, cytoplasmic polyadenylation elements, embryonic cytoplasmic polyadenylation elements and Pumilio binding element per gene-expression type. Corrected p-values above 0.05 are indicated by a dash.

### Supplemental_File_S7.xlsx

Overrepresentation of KEGG pathways, Gene Ontology (GO) categories, Chromosomes, Interpro protein ids, SMART ids (http://smart.embl.de/), PIR keywords (Protein Information Resource), Uniprot Sequence Feature and Tissue (http://www.uniprot.org/uniprot) in expressed genes, gene-expression types and Multi-Level Type genes against an appropriate background (cf. Material & Methods) with an EASE-score < 0.05 and a fold enrichment larger than 1.5.

### Supplemental_File_S8.xlsx

Genes carrying specific miRNA targets in their 3’UTR that are overrepresented and underrepresented in specific categories. Tab A. Counts of overrepresented and underrepresented miRNA targets in the set of expressed genes, their p-values and corrected p-values. Actual expression of the miRNA at gastrulation is indicated by a ‘1’. Tab B. All miRNA targets overrepresented in gene-expression types. Only types that contain overrepresented targets are shown. The yellow shading indicates the specific miRNA target - dynamic type combination with a called overrepresentation (with a corrected p-value < 0.15). Tab C. All miRNA targets underrepresented in gene-expression types and presented similar to tab B. Tab D.

Overrepresentation and underrepresentation in the set of expressed genes of all miRNA targets. Tab E and F. Respectively Overrepresentation and underrepresentation of all miRNA targets in all dynamic categories.

### Supplemental_File_S9.pdf

Total number of miRNAs normalized with the assumption of equal amounts of small RNA in the 16 samples taken at gastrulation (Figure 1C).

### Supplemental_File_S10.pdf

Relative counts of read mapped to miRNAs in 16 samples taken at gastrulation (Figure 1C).

### Supplemental_File_S11.xlsx

Experiment information on the small RNA sequencing experiments: 18 samples in a developmental time course (upper panel) and 16 samples of embryos taken at gastrulation around 50% epiboly.

### Supplemental_File_S12.pdf

Anti-correlating miRNA expression (page 1 and 2) with a correlation coefficient < −0.8 and correlating miRNA expression (page 3 to 10) with a correlation coefficient > 0.9 in 16 samples taken at gastrulation (Figure 1C). Expression is given as log 2 relative counts. Samples without any mapping are indicated as non-expressed.

