## Supplemental File S1 for "Cellular Factors Involved in Transcriptome Dynamics in Early Zebrafish Embryogenesis"

### Number of Expressed Genes per Chromosome

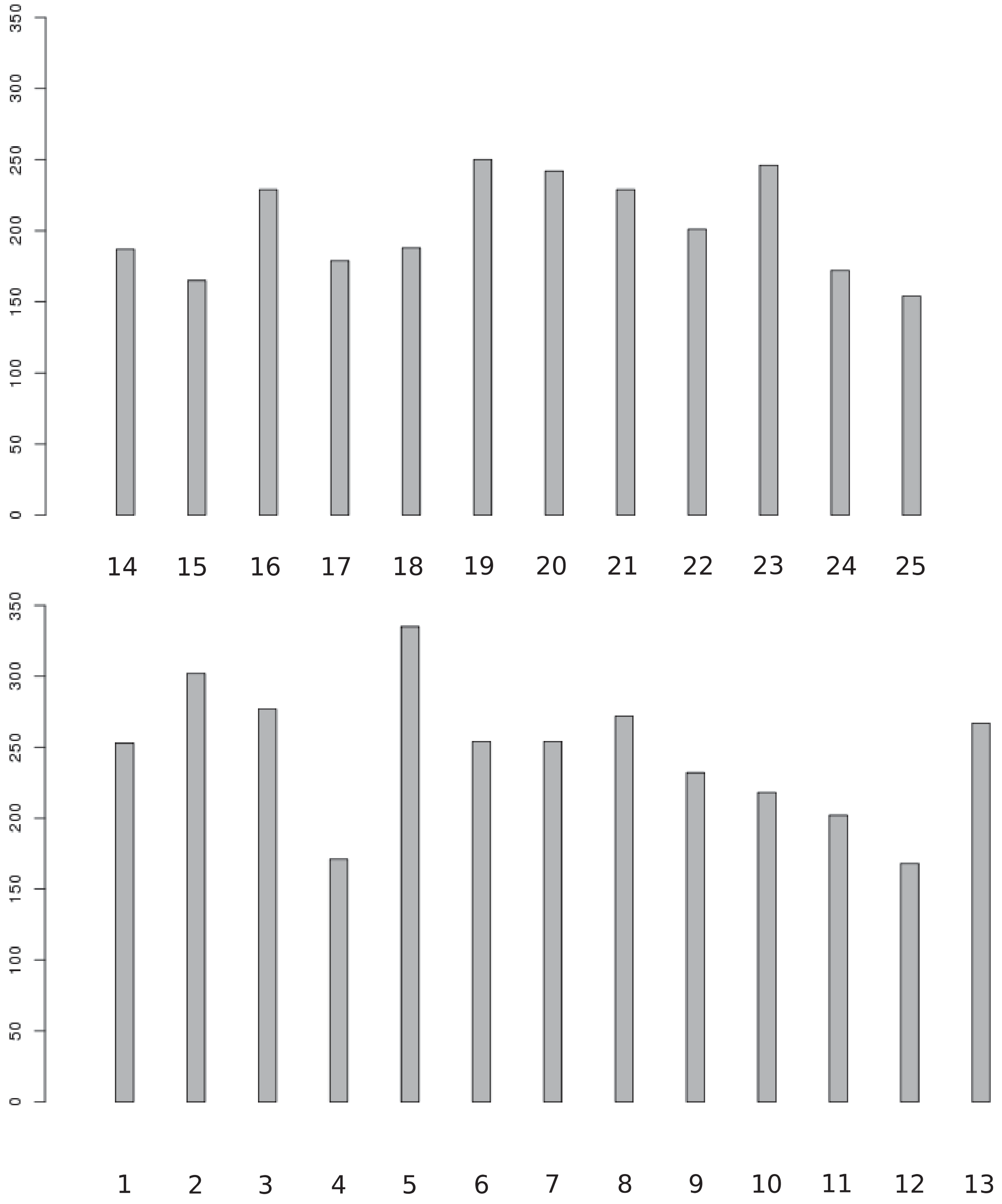

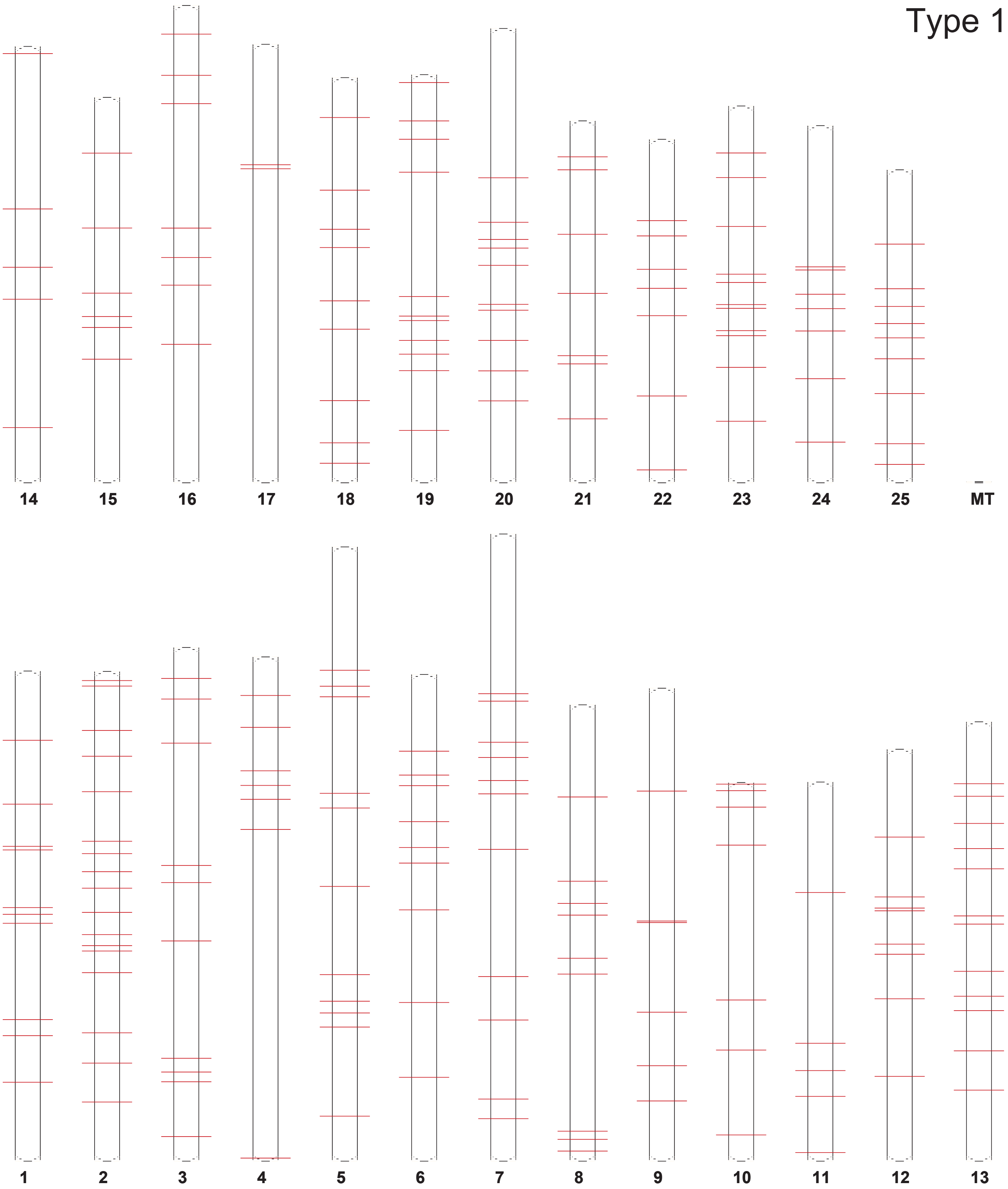
Type 1
MT
1
2
3
4
5
6
7
8
9
10
11
12
13

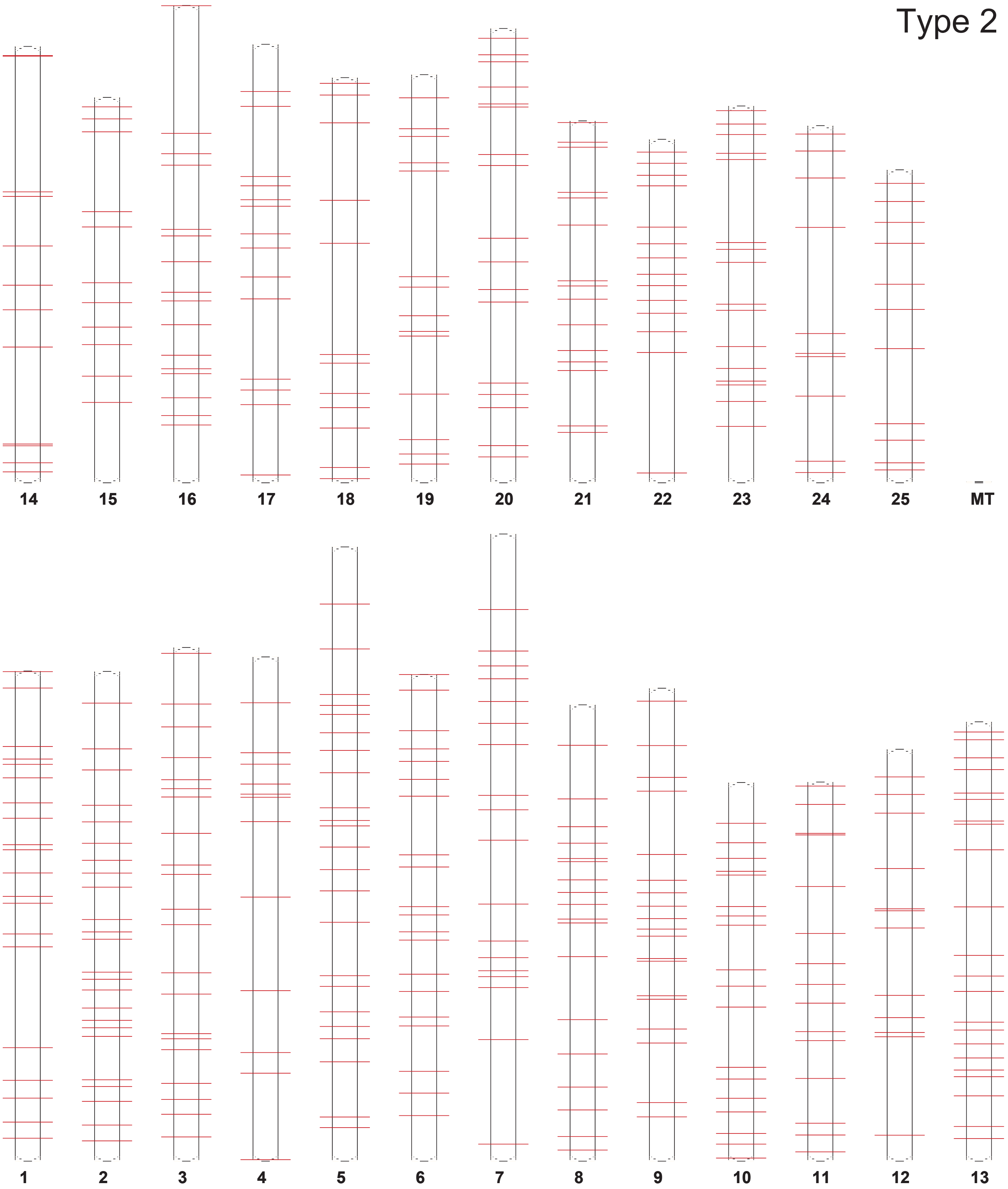
Type 2
MT
1
2
3
4
5
6
7
8
9
10
11
12
13

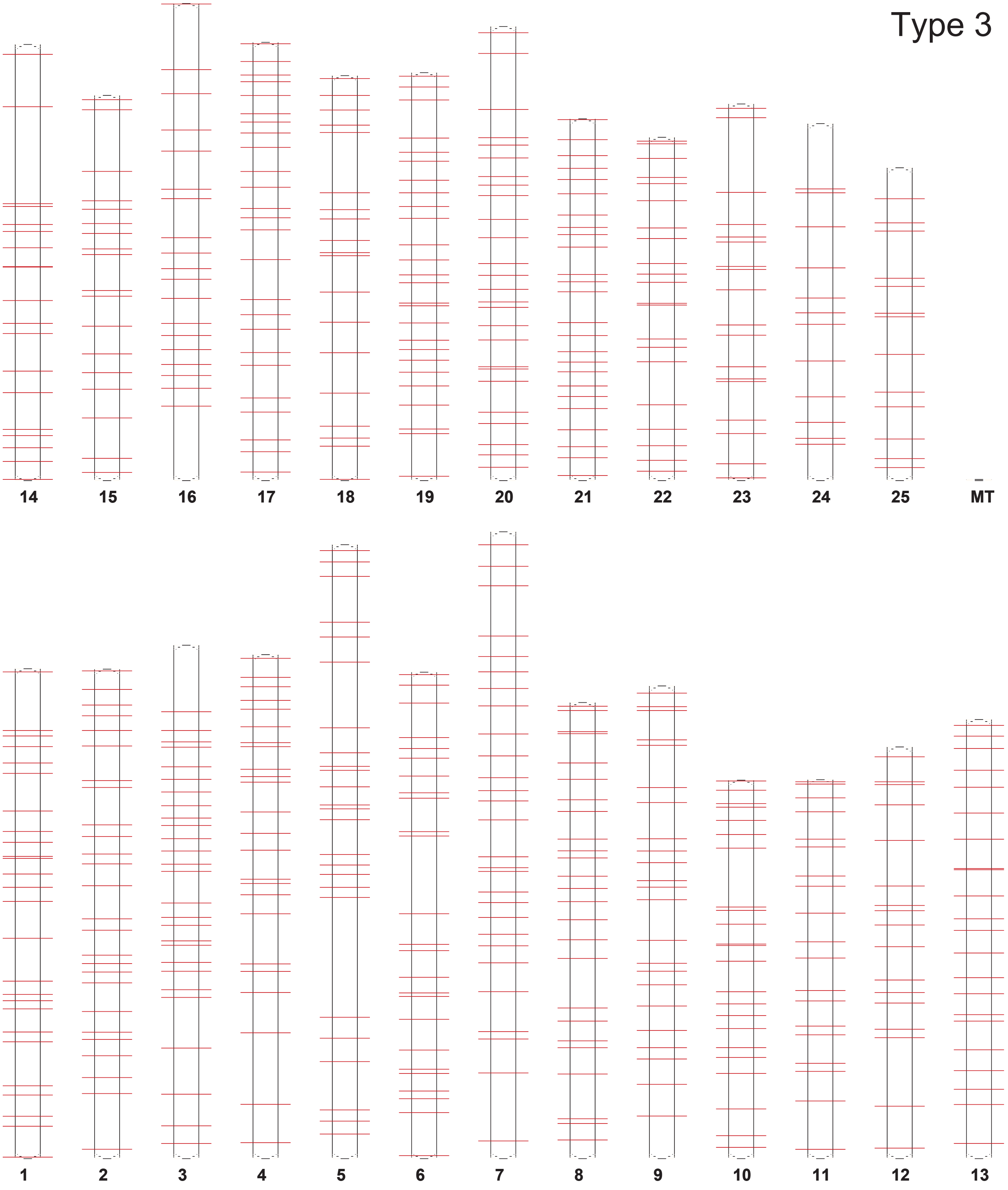
Type 3
MT
1
2
3
4
5
6
7
8
9
10
11
12
13

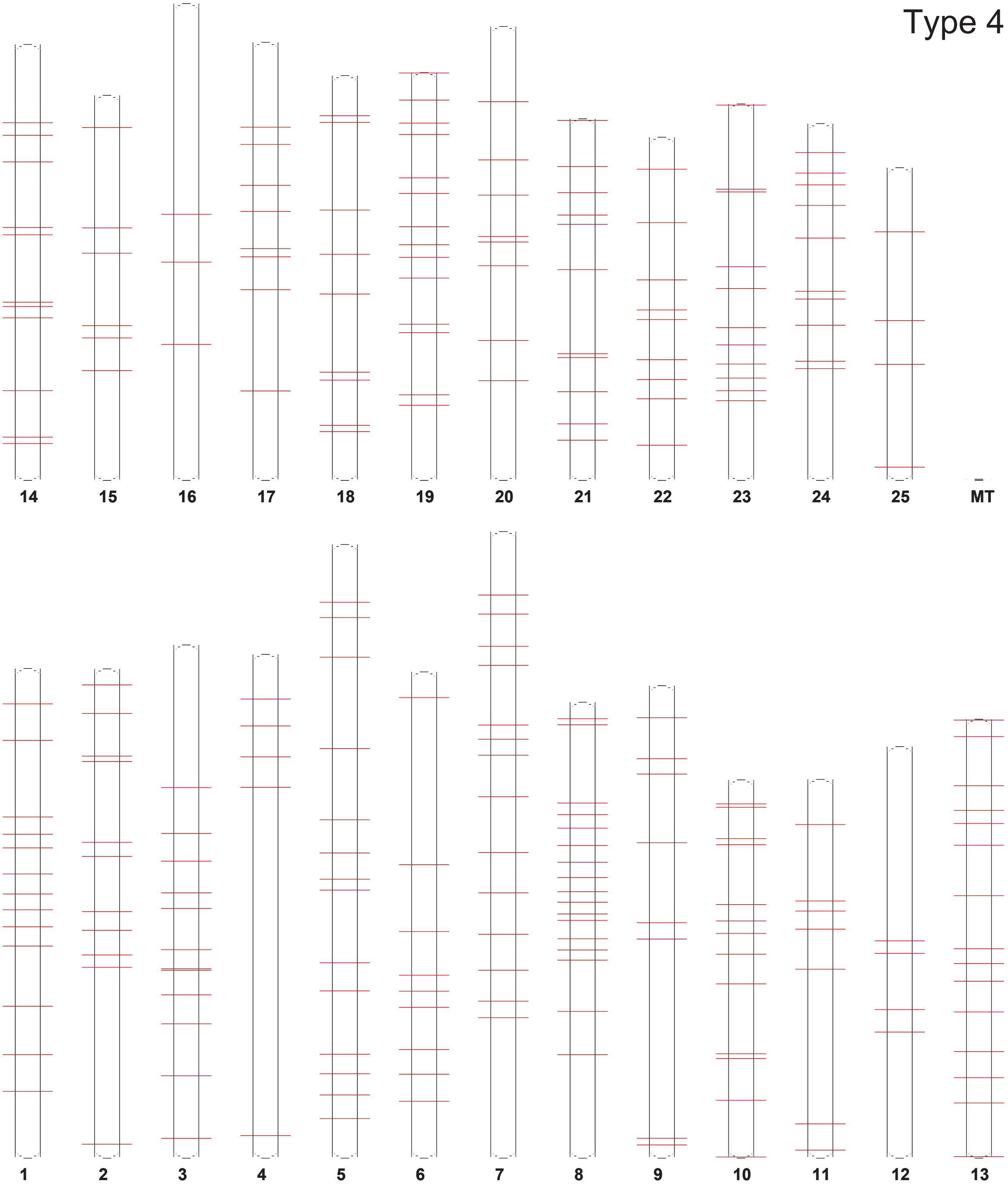
Type 4
MT
1
2
3
4
5
6
7
8
9
10
11
12
13

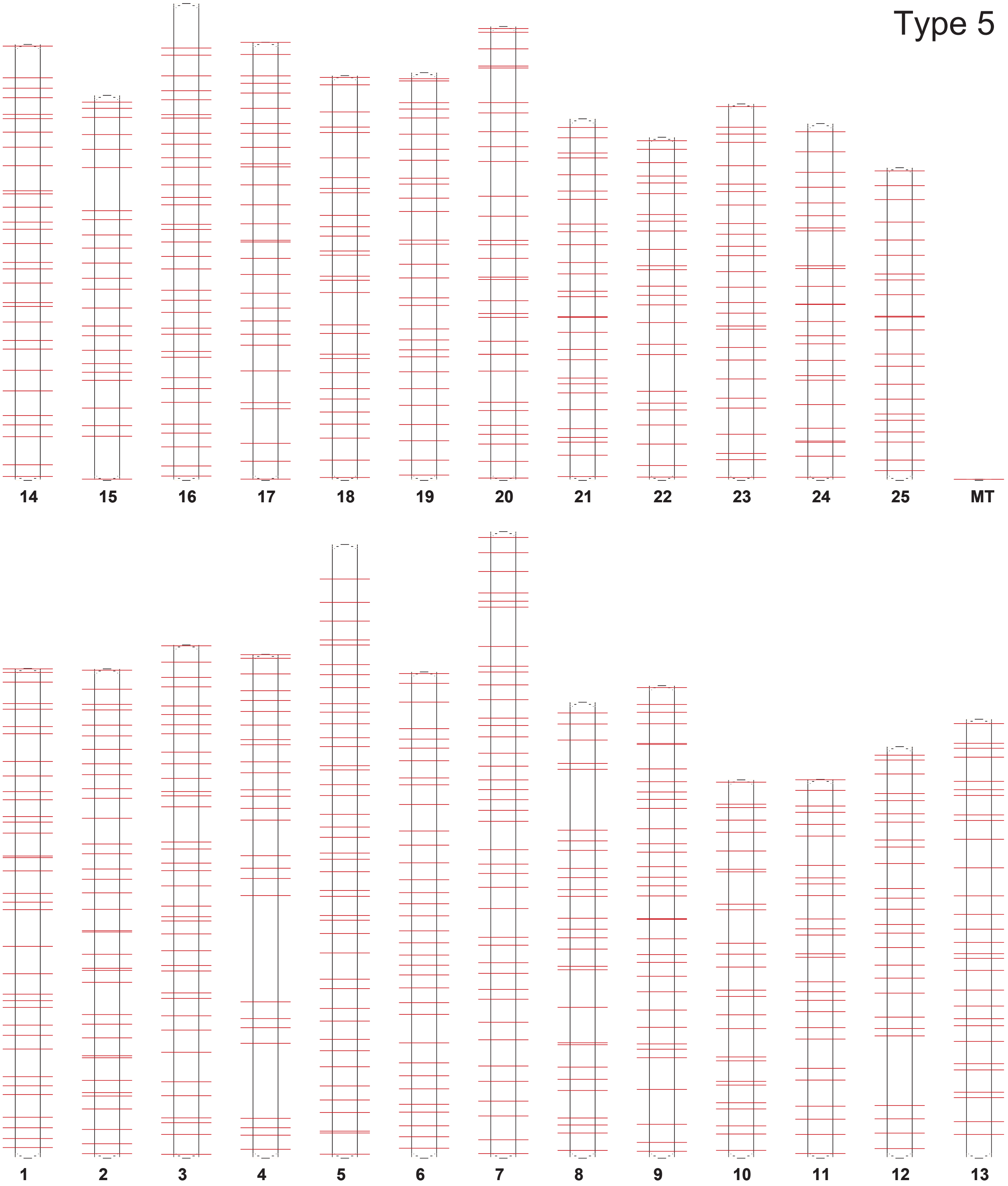
Type 5
MT
1
2
3
4
5
6
7
8
9
10
11
12
13

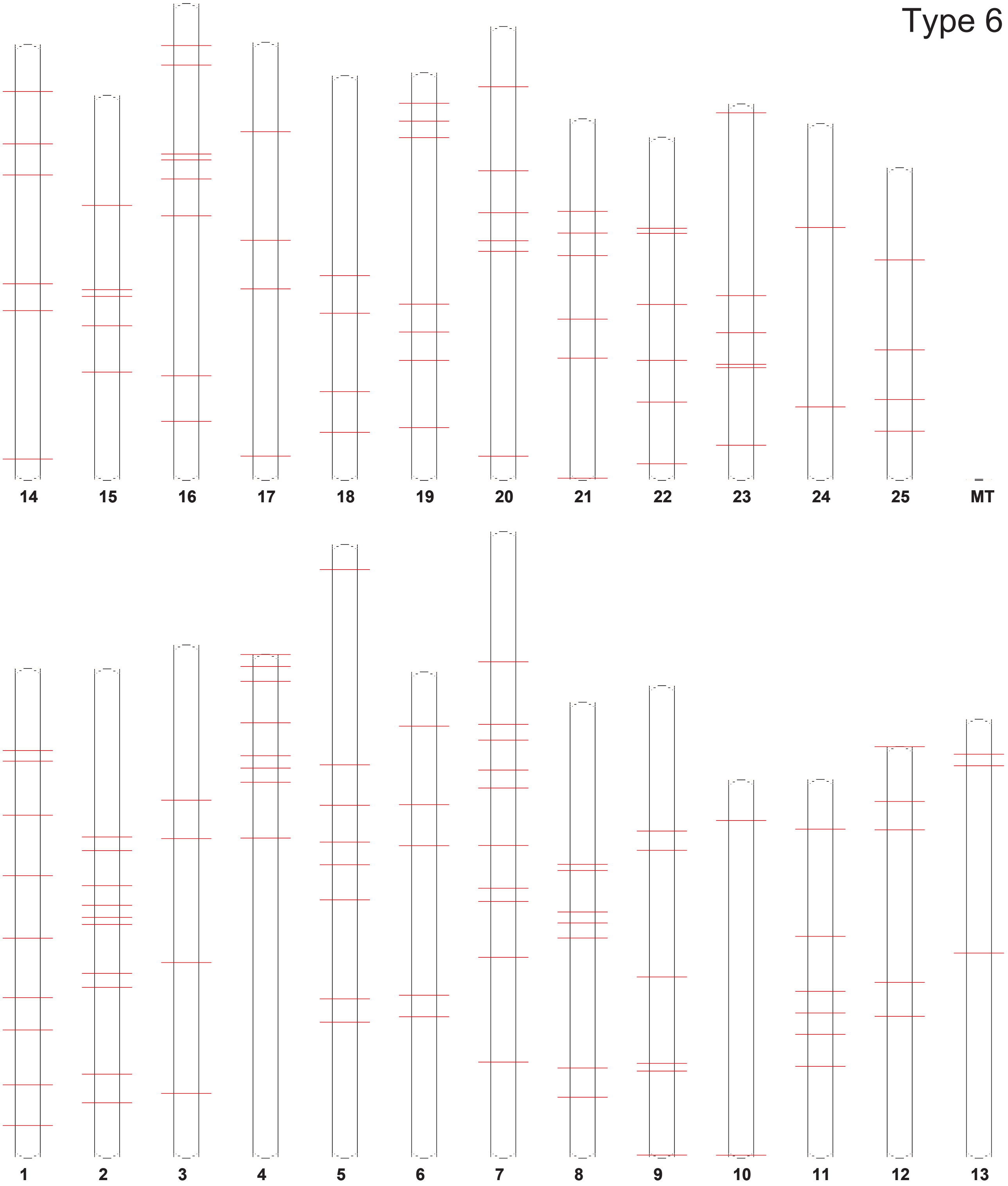
Type 6
MT
1
2
3
4
5
6
7
8
9
10
11
12
13

Type 7

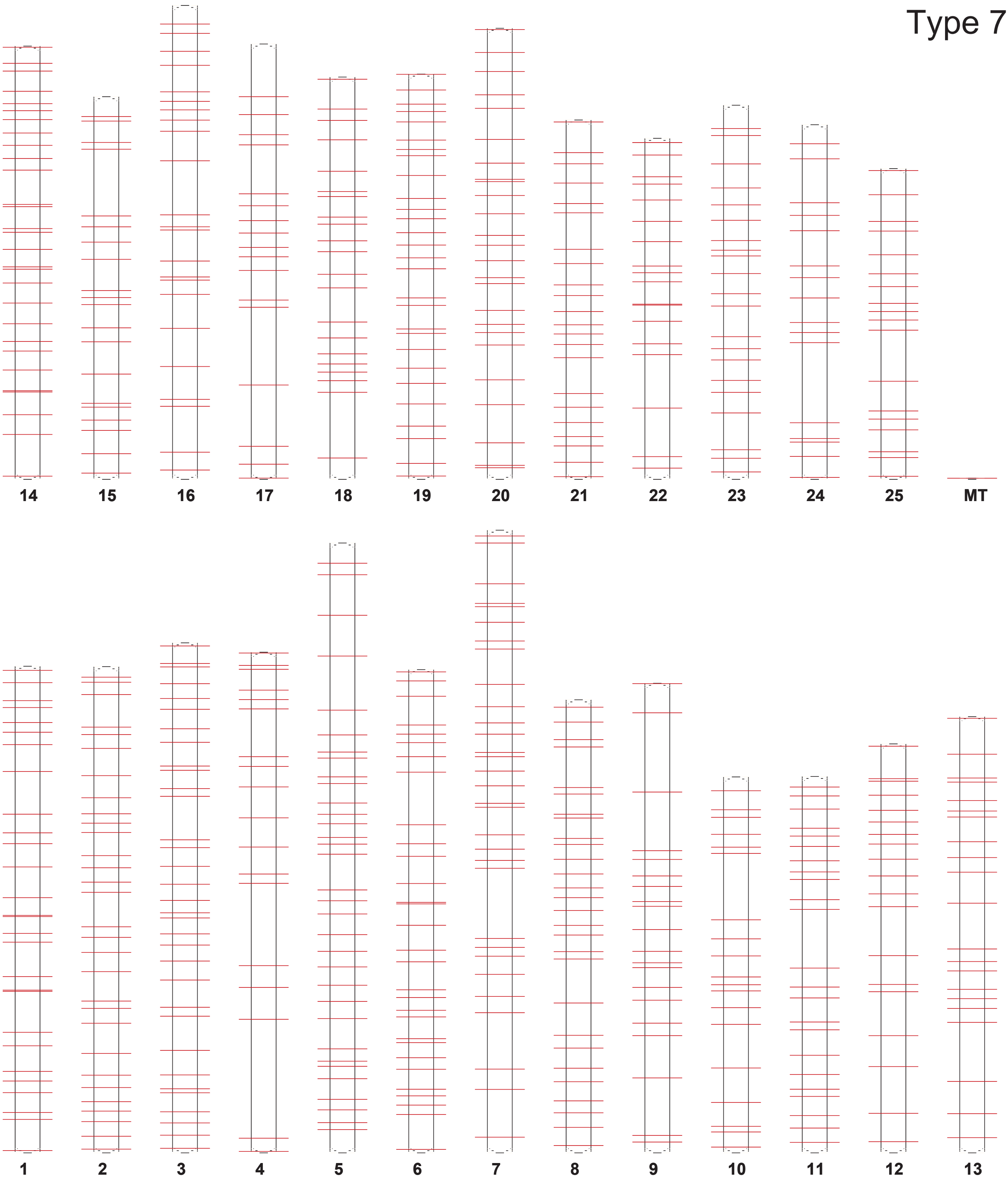

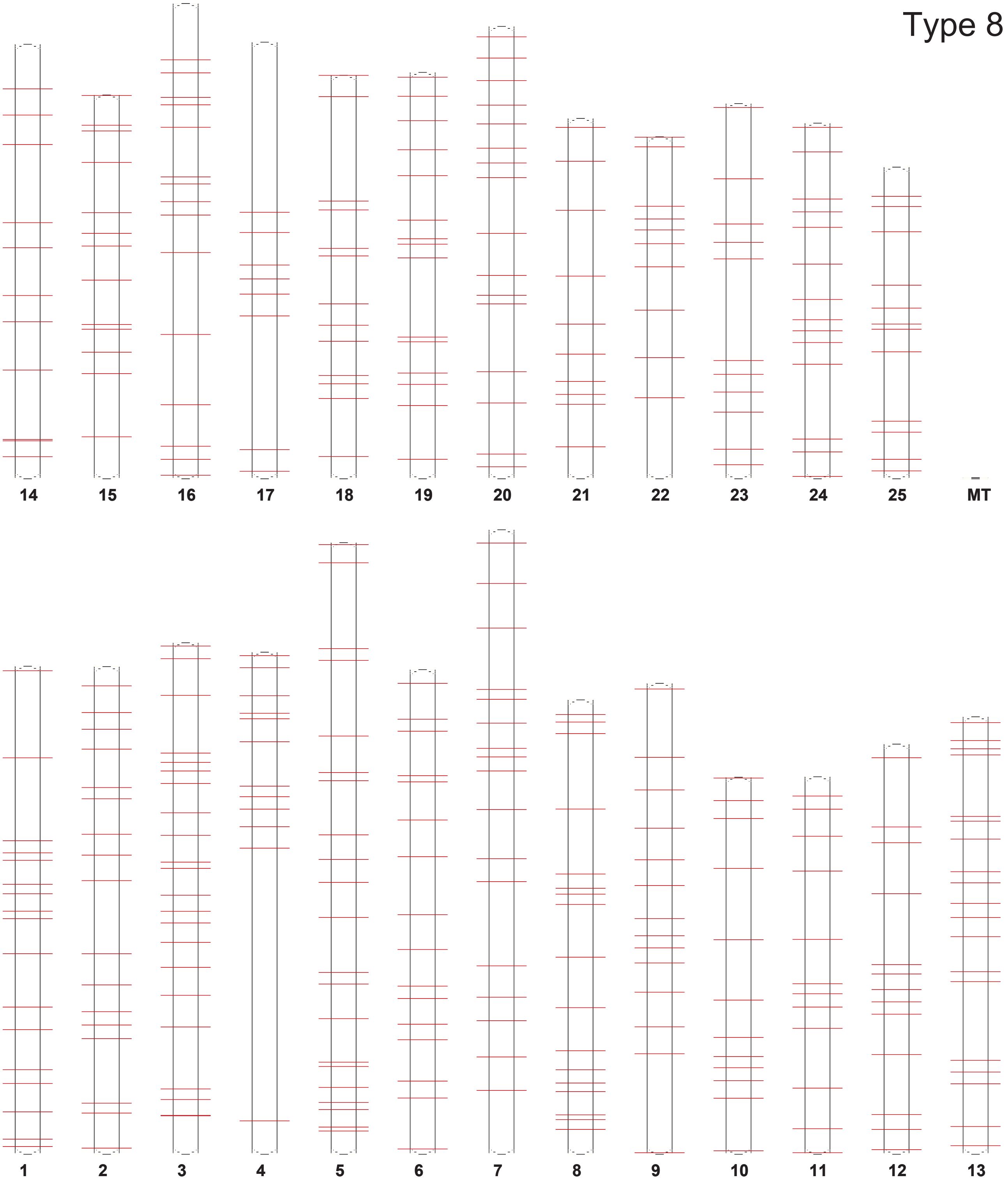
Type 8
MT
1
2
3
4
5
6
7
8
9
10
11
12
13

#### Type 9

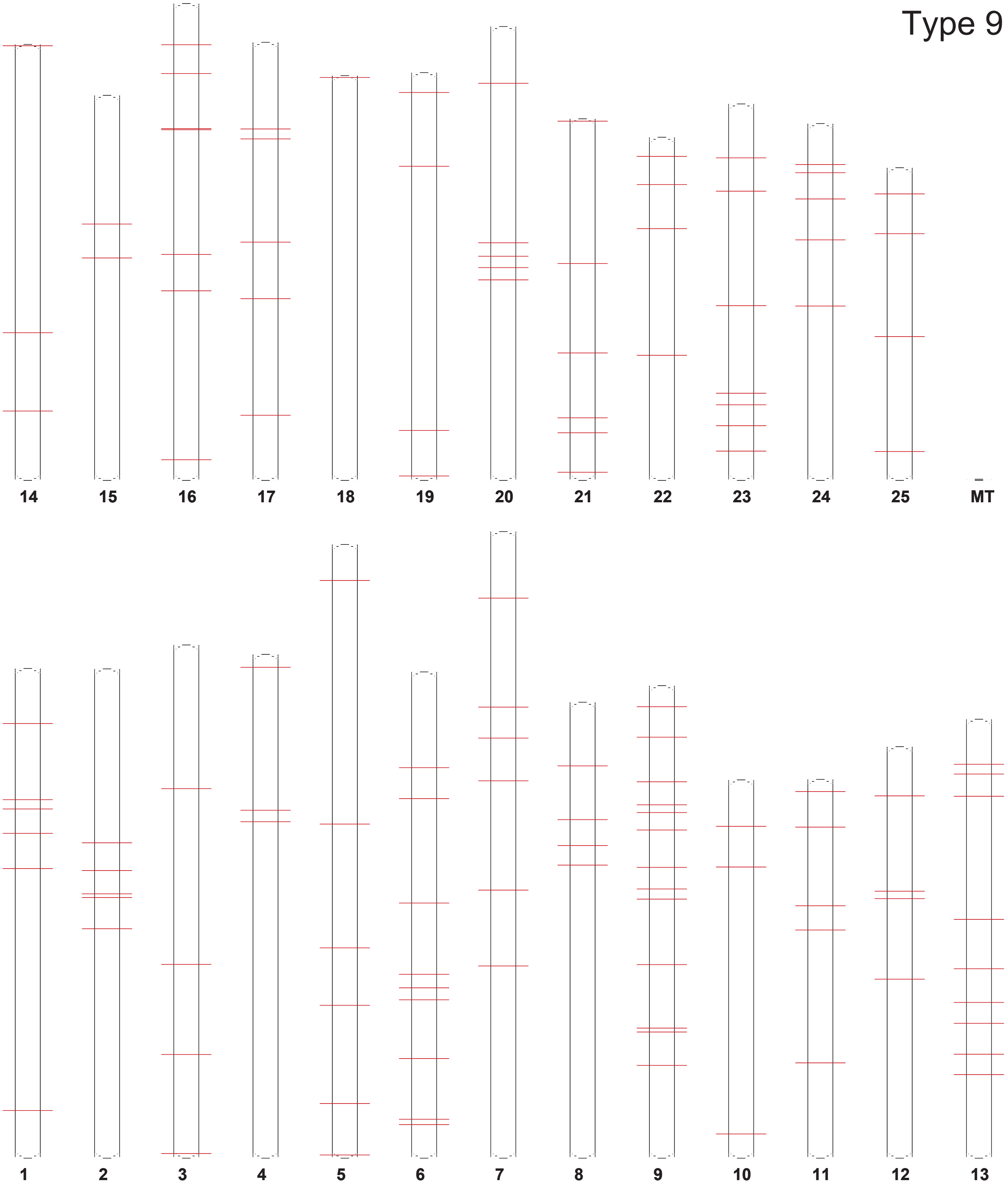

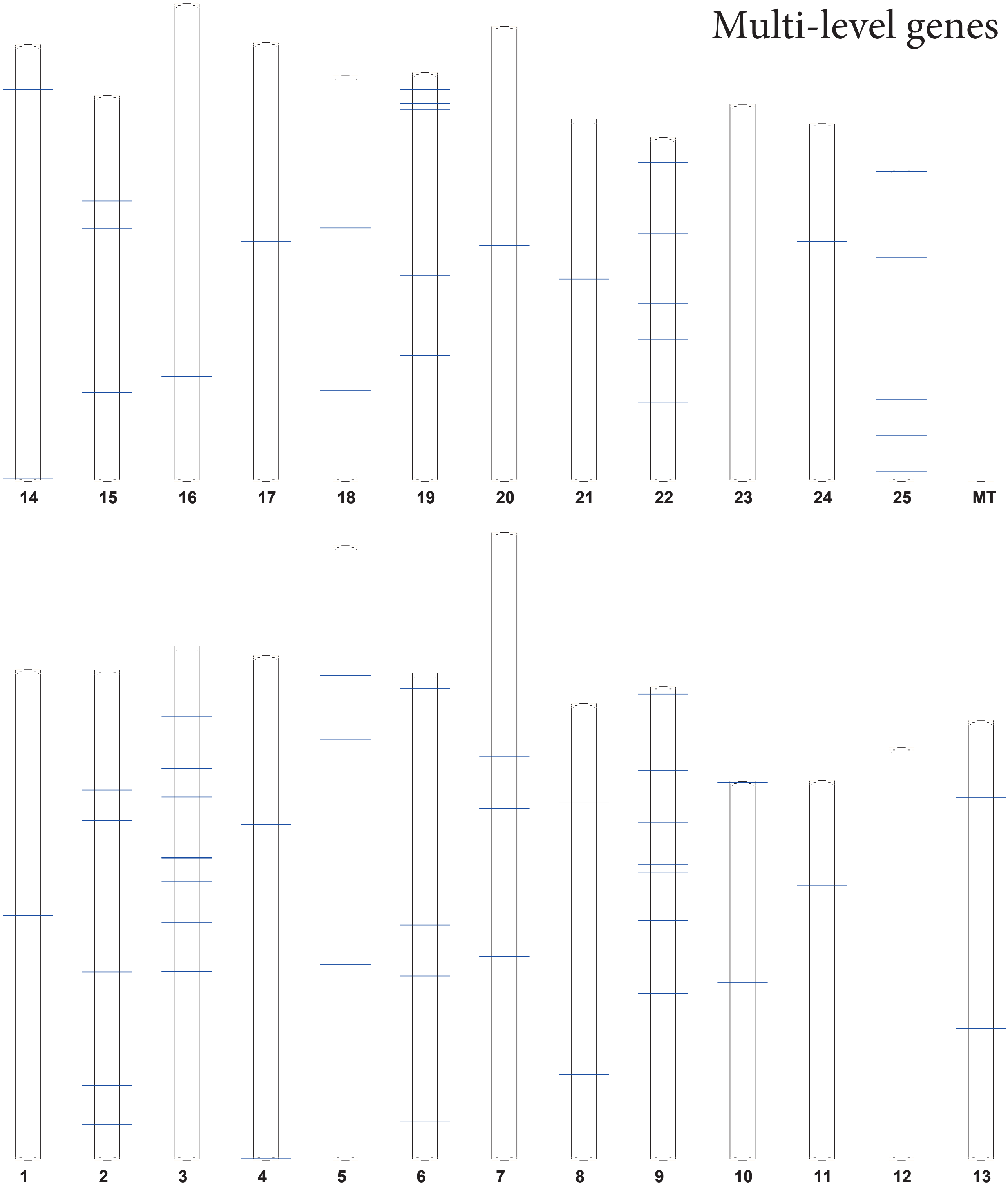

Multi-level genes
MT
1
2
3
4
5
6
7
8
9
10
11
12
13

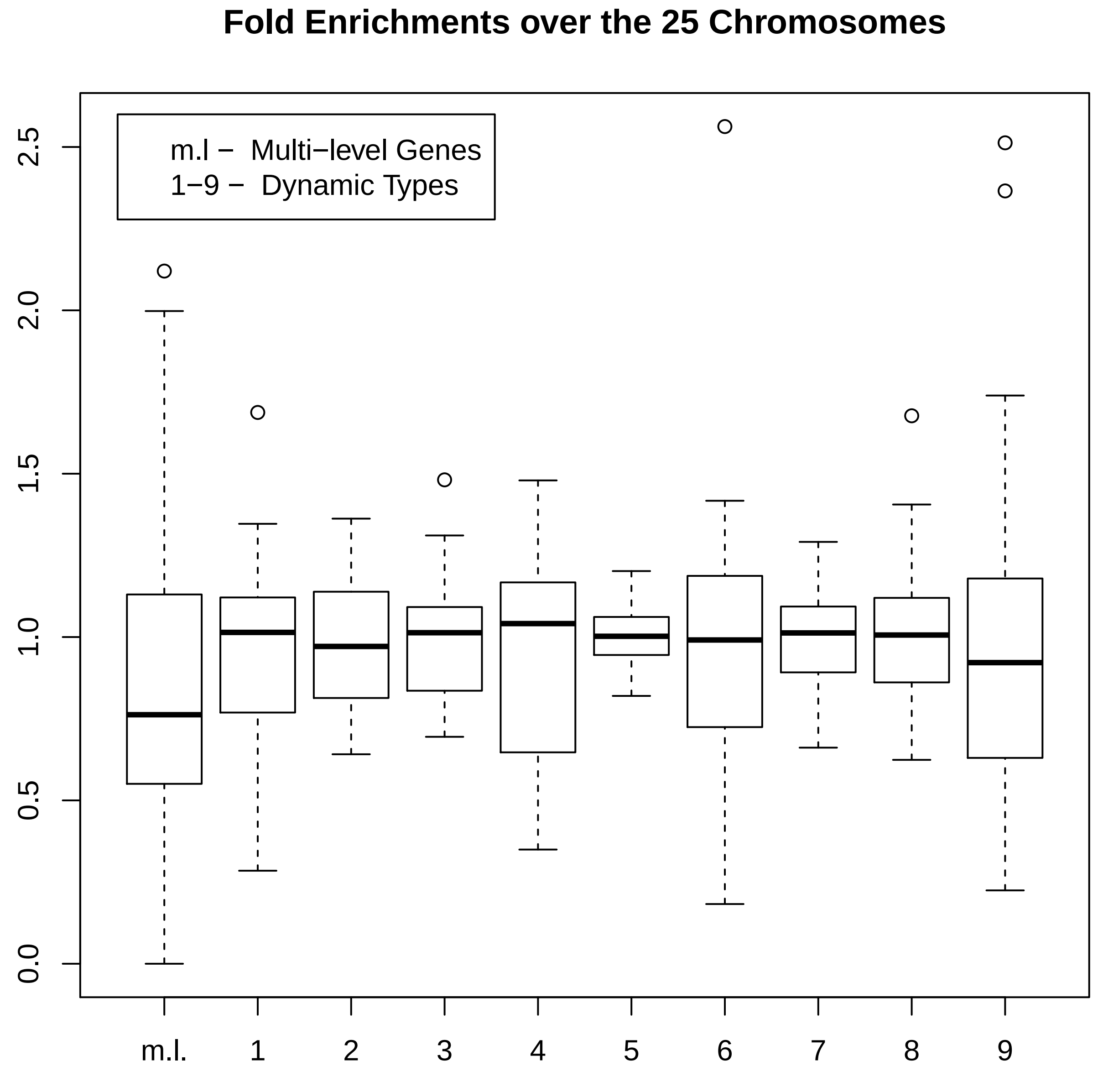

Fold Enrichments over the 25 Chromosomes
m.l − Multi−level Genes
1−9 − Dynamic Types
0.0
0.5
1.0
1.5
2.0
2.5
m.l.
1
2
3
4
5
6
7
8
9
