## Supplemental File S5 for "Cellular Factors Involved in Transcriptome Dynamics in Early Zebrafish Embryogenesis"

### Motif Enrichment in Promoter Sequences of Transcription Dynamic Types

300 nucleotides upstream and 300 nucleotides downstream of transcription start site.

Transcription Dynamic Types without enrichment at a corrected p-value below 0.5 are not shown.


| Type | Number of motifs | page |
|---|---|---|
| Non-expressed upstream | 137 | 2 |
| Non-expressed downstream | 71 | 16 |
| Expressed upstream | 217 | 24 |
| Expressed downstream | 157 | 45 |
| Down upstream | 12 | 61 |
| Down downstream | 11 | 64 |
| Up upstream | 27 | 67 |
| Type 1 downstream | 7 | 72 |
| Type 2 downstream | 1 | 75 |
| Type 4 downstream | 1 | 77 |
| Type 6 upstream | 8 | 79 |
| Type 7 downstream | 1 | 82 |
| Type 8 upstream | 2 | 84 |
| Type 9 upstream | 2 | 86 |

Non-expressed upstream

### AME
Analysis of Motif Enrichment

For further information on how to interpret these results or to get a copy of the MEME software please access http://meme-suite.org.

If you use AME in your research, please cite the following paper:
Robert McLeay and Timothy L. Bailey, "Motif Enrichment Analysis: A unified framework and method evaluation", *BMC Bioinformatics*, **11**:165, 2010, doi:10.1186/1471-2105-11-165. [full text]

Enriched Motifs | Input Files | Program information

#### Enriched Motifs

Fixed partition size: number of primary sequences (15480)

Sequence motif score: avg_odds
Background model source: file background.model
Background model frequencies: 0.291,0.209,0.209,0.291
Total pseudocount added to a motif column: 0.25

Statistical test: Wilcoxon rank-sum test
Ranksum method: quick
Threshold *p*-value for reporting results: 0.05
Number of multiple tests for Bonferroni correction: #Motifs × #PartitionsTested = 591 × 1 = 591Fixed partition size: number of primary sequences (15480)

Sequence motif score: avg_odds
Background model source: file background.model
Background model frequencies: 0.291,0.209,0.209,0.291
Total pseudocount added to a motif column: 0.25

Statistical test: Wilcoxon rank-sum test
Ranksum method: quick
Threshold *p*-value for reporting results: 0.05
Number of multiple tests for Bonferroni correction: #Motifs × #PartitionsTested = 591 × 1 = 591

| Logo | Database | ID | Name | *p*-value | Adjusted *p*-value |
|---|---|---|---|---|---|
| 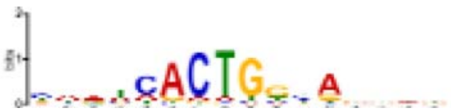 | uniprobe mouse | UP00031_2 | Zbtb3_secondary | 2.22e-37 | 1.31e-34 |

| | | | | | |
|---|---|---|---|---|---|
| 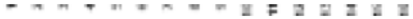 | | | | | |
| 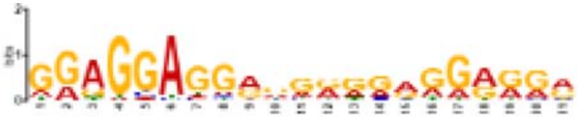 | JASPAR CORE 2014 vertebrates | MA0528.1 | ZNF263 | 3.48e-33 | 2.06e-30 |
| 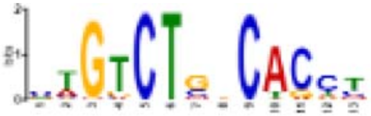 | JASPAR CORE 2014 vertebrates | MA0513.1 | SMAD2::SMAD3::SMAD4 | 1.32e-31 | 7.80e-29 |
| 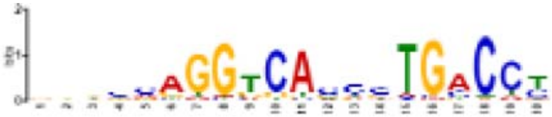 | JASPAR CORE 2014 vertebrates | MA0112.2 | ESR1 | 7.42e-30 | 4.38e-27 |
| 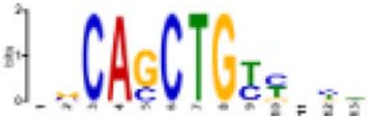 | JASPAR CORE 2014 vertebrates | MA0499.1 | Myod1 | 2.60e-28 | 1.54e-25 |
| 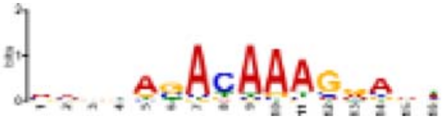 | uniprobe mouse | UP00101_2 | Sox12_secondary | 2.32e-27 | 1.37e-24 |
| 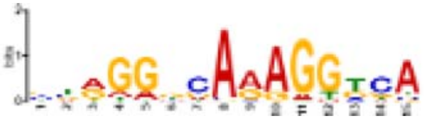 | JASPAR CORE 2014 vertebrates | MA0065.2 | PPARG::RXRA | 1.65e-26 | 9.78e-24 |
| 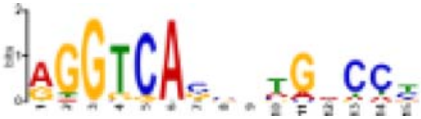 | JASPAR CORE 2014 vertebrates | MA0258.2 | ESR2 | 1.24e-25 | 7.35e-23 |
| 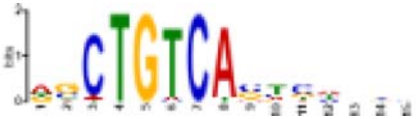 | JASPAR CORE 2014 vertebrates | MA0498.1 | Meis1 | 3.90e-25 | 2.30e-22 |
| 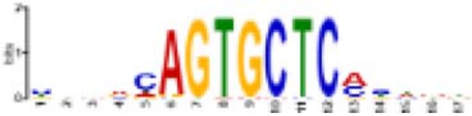 | uniprobe mouse | UP00095_1 | Zfp691_primary | 9.91e-25 | 5.85e-22 |
| 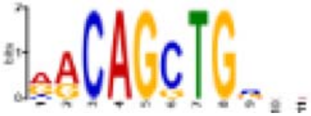 | JASPAR CORE 2014 vertebrates | MA0521.1 | Tcf12 | 1.09e-24 | 6.47e-22 |
| 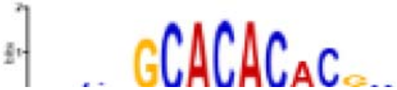 | uniprobe mouse | UP00042_2 | Gm397_secondary | 4.68e-24 | 2.76e-21 |

| | | | | | |
|---|---|---|---|---|---|
| 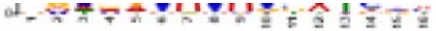 | | | | | |
| 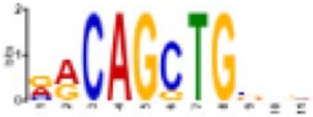 | JASPAR CORE 2014 vertebrates | MA0500.1 | Myog | 2.65e-23 | 1.57e-20 |
| 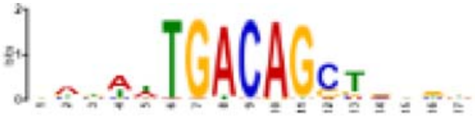 | uniprobe mouse | UP00122_1 | Tgif1_2342.2 | 7.80e-23 | 4.61e-20 |
| 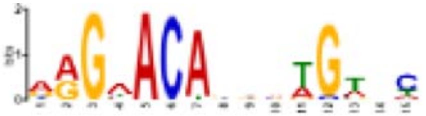 | JASPAR CORE 2014 vertebrates | MA0007.2 | AR | 9.31e-23 | 5.50e-20 |
| 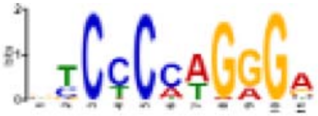 | JASPAR CORE 2014 vertebrates | MA0154.2 | EBF1 | 1.52e-22 | 8.97e-20 |
| 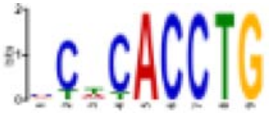 | JASPAR CORE 2014 vertebrates | MA0103.2 | ZEB1 | 2.89e-22 | 1.71e-19 |
| 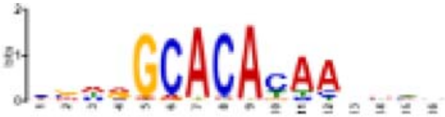 | uniprobe mouse | UP00026_2 | Zscan4_secondary | 3.38e-21 | 2.00e-18 |
| 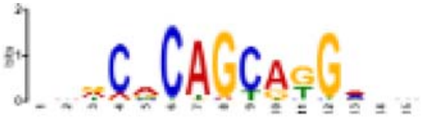 | uniprobe mouse | UP00006_2 | Zic3_secondary | 5.15e-21 | 3.05e-18 |
| 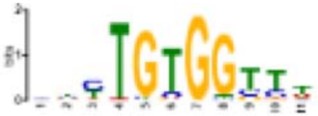 | JASPAR CORE 2014 vertebrates | MA0002.2 | RUNX1 | 6.43e-21 | 3.80e-18 |
| 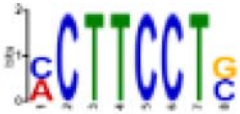 | JASPAR CORE 2014 vertebrates | MA0598.1 | EHF | 7.98e-20 | 4.72e-17 |
| 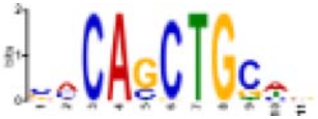 | JASPAR CORE 2014 vertebrates | MA0522.1 | Tcf3 | 8.02e-20 | 4.74e-17 |
| 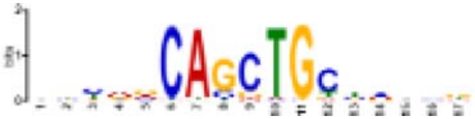 | uniprobe mouse | UP00099_1 | Ascl2_primary | 1.79e-18 | 1.06e-15 |
| 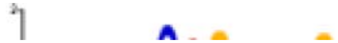 | | | | [illegible] | |

| | | | | | |
|---|---|---|---|---|---|
| 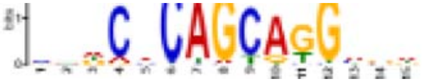 | uniprobe mouse | UP00102_2 | Zic1_secondary | 2.28e-18 | 1.35e-15 |
| 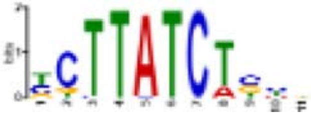 | JASPAR CORE 2014 vertebrates | MA0482.1 | Gata4 | 3.18e-18 | 1.88e-15 |
| 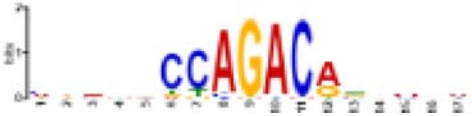 | uniprobe mouse | UP00000_1 | Smad3_primary | 4.37e-18 | 2.58e-15 |
| 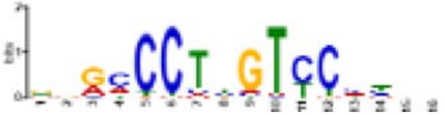 | uniprobe mouse | UP00082_2 | Zfp187_secondary | 4.99e-18 | 2.95e-15 |
| 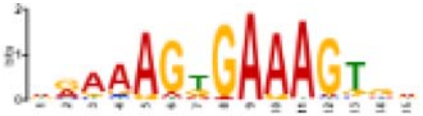 | JASPAR CORE 2014 vertebrates | MA0508.1 | PRDM1 | 5.12e-18 | 3.02e-15 |
| 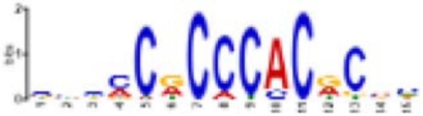 | JASPAR CORE 2014 vertebrates | MA0472.1 | EGR2 | 1.29e-17 | 7.64e-15 |
| 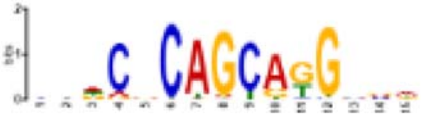 | uniprobe mouse | UP00057_2 | Zic2_secondary | 1.74e-17 | 1.03e-14 |
| 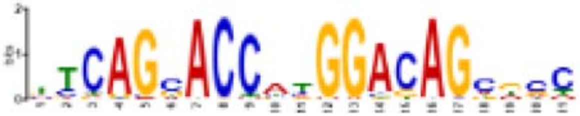 | JASPAR CORE 2014 vertebrates | MA0138.2 | REST | 2.50e-17 | 1.48e-14 |
| 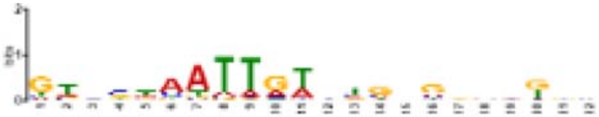 | uniprobe mouse | UP00034_2 | Sox7_secondary | 6.05e-17 | 3.57e-14 |
| 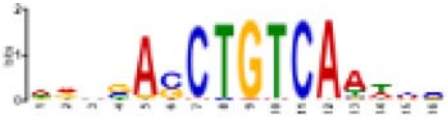 | uniprobe mouse | UP00203_1 | Pknox1_2364.2 | 9.90e-17 | 5.85e-14 |
| 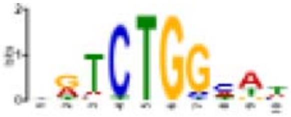 | JASPAR CORE 2014 vertebrates | MA0092.1 | Hand1::Tcfe2a | 1.64e-16 | 9.68e-14 |

| | | | | | |
|---|---|---|---|---|---|
| 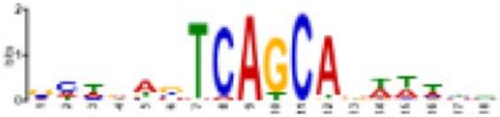 | JASPAR CORE 2014 vertebrates | MA0495.1 | MAFF | 2.34e-16 | 1.38e-13 |
| 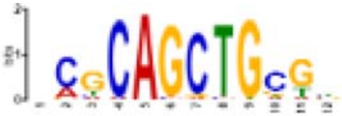 | JASPAR CORE 2014 vertebrates | MA0048.1 | NHLH1 | 2.40e-16 | 1.42e-13 |
| 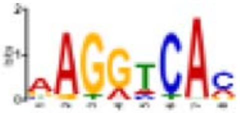 | JASPAR CORE 2014 vertebrates | MA0160.1 | NR4A2 | 3.71e-16 | 2.19e-13 |
| 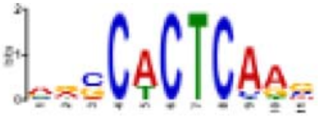 | JASPAR CORE 2014 vertebrates | MA0503.1 | Nkx2-5 | 4.81e-16 | 2.84e-13 |
| 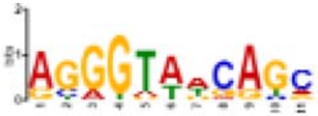 | JASPAR CORE 2014 vertebrates | MA0111.1 | Spz1 | 5.04e-16 | 2.98e-13 |
| 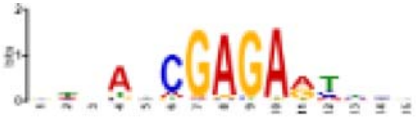 | uniprobe mouse | UP00040_2 | Irf5_secondary | 8.12e-16 | 4.80e-13 |
| 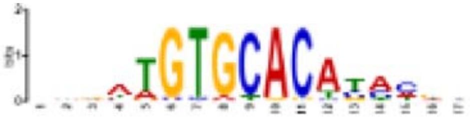 | uniprobe mouse | UP00042_1 | Gm397_primary | 8.61e-16 | 5.09e-13 |
| 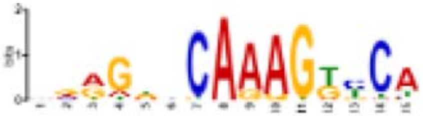 | JASPAR CORE 2014 vertebrates | MA0484.1 | HNF4G | 1.04e-15 | 6.14e-13 |
| 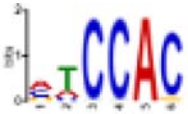 | JASPAR CORE 2014 vertebrates | MA0130.1 | ZNF354C | 3.45e-15 | 2.04e-12 |
| 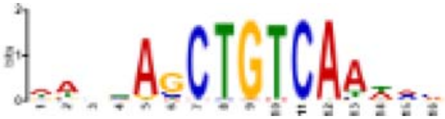 | uniprobe mouse | UP00258_1 | Tgif2_3451.1 | 3.70e-15 | 2.19e-12 |
| 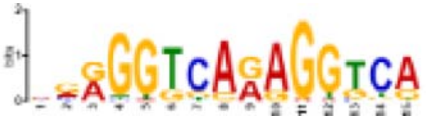 | JASPAR CORE 2014 vertebrates | MA0504.1 | NR2C2 | 9.86e-15 | 5.82e-12 |
| 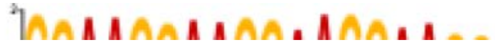 | JASPAR CORE 2014 [illegible] | [illegible] | [illegible] | [illegible] | [illegible] |

| | | | | | |
|---|---|---|---|---|---|
| 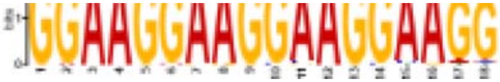 | JASPAR CORE 2014 vertebrates | MA0149.1 | EWSR1-FLI1 | 1.30e-14 | 7.69e-12 |
| 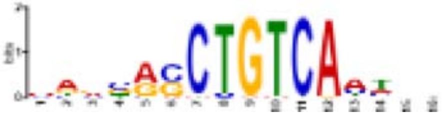 | uniprobe mouse | UP00205_1 | Pknox2_3077.2 | 1.46e-14 | 8.65e-12 |
| 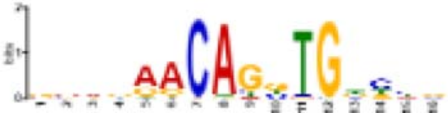 | uniprobe mouse | UP00036_1 | Myf6_primary | 1.78e-14 | 1.05e-11 |
| 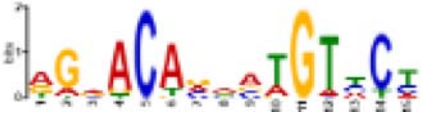 | JASPAR CORE 2014 vertebrates | MA0113.2 | NR3C1 | 1.80e-14 | 1.06e-11 |
| 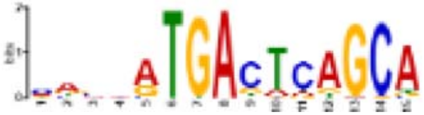 | JASPAR CORE 2014 vertebrates | MA0150.2 | Nfe2l2 | 2.52e-14 | 1.49e-11 |
| 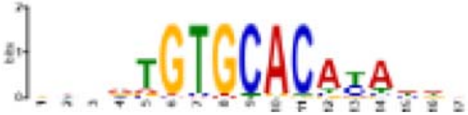 | uniprobe mouse | UP00026_1 | Zscan4_primary | 2.97e-14 | 1.75e-11 |
| 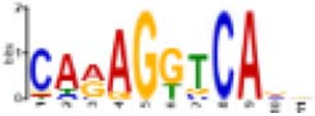 | JASPAR CORE 2014 vertebrates | MA0512.1 | Rxra | 3.18e-14 | 1.88e-11 |
| 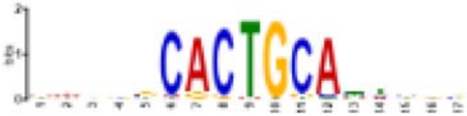 | uniprobe mouse | UP00031_1 | Zbtb3_primary | 4.88e-14 | 2.88e-11 |
| 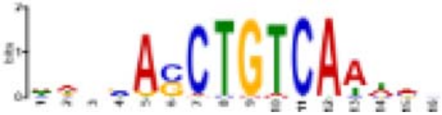 | uniprobe mouse | UP00226_1 | Mrg1_2246.2 | 5.83e-14 | 3.44e-11 |
| 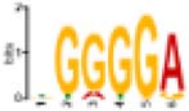 | JASPAR CORE 2014 vertebrates | MA0056.1 | MZF1_1-4 | 6.02e-14 | 3.56e-11 |
| 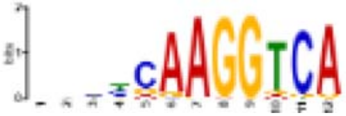 | JASPAR CORE 2014 vertebrates | MA0141.2 | Esrrb | 6.27e-14 | 3.71e-11 |

| | | | | | |
|---|---|---|---|---|---|
| 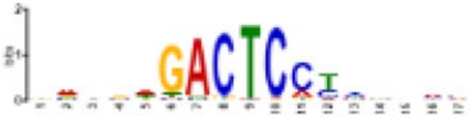 | uniprobe mouse | UP00095_2 | Zfp691_secondary | 7.51e-14 | 4.44e-11 |
| 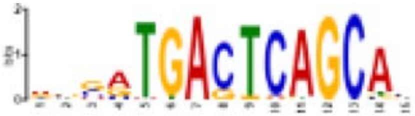 | JASPAR CORE 2014 vertebrates | MA0591.1 | Bach1::Mafk | 8.25e-14 | 4.88e-11 |
| 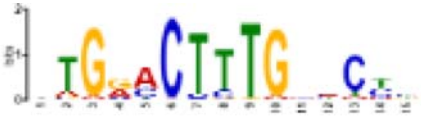 | JASPAR CORE 2014 vertebrates | MA0114.2 | HNF4A | 3.33e-13 | 1.97e-10 |
| 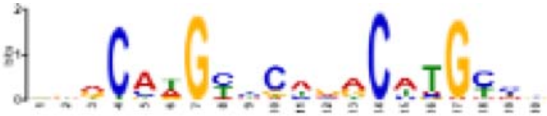 | JASPAR CORE 2014 vertebrates | MA0525.1 | TP63 | 3.68e-13 | 2.17e-10 |
| 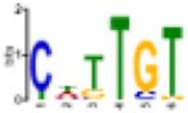 | JASPAR CORE 2014 vertebrates | MA0442.1 | SOX10 | 4.40e-13 | 2.60e-10 |
| 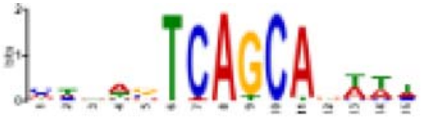 | JASPAR CORE 2014 vertebrates | MA0496.1 | MAFK | 5.44e-13 | 3.21e-10 |
| 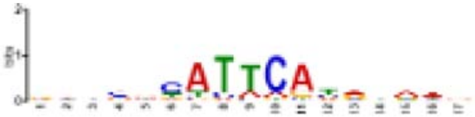 | uniprobe mouse | UP00014_2 | Sox17_secondary | 6.01e-13 | 3.55e-10 |
| 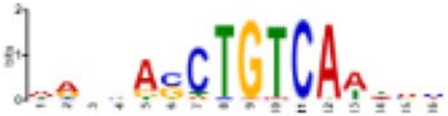 | uniprobe mouse | UP00210_1 | Mrg2_2302.1 | 8.07e-13 | 4.77e-10 |
| 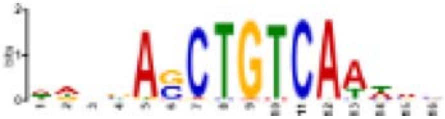 | uniprobe mouse | UP00186_1 | Meis1_2335.1 | 1.37e-12 | 8.09e-10 |
| 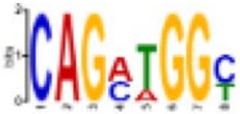 | JASPAR CORE 2014 vertebrates | MA0461.1 | Atoh1 | 3.61e-12 | 2.13e-9 |
| 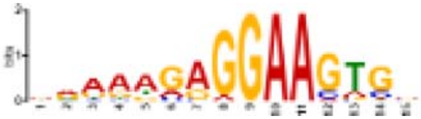 | JASPAR CORE 2014 vertebrates | MA0080.3 | Spi1 | 5.36e-12 | 3.17e-9 |
| 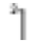 | | | | [illegible] | |

| | | | | | |
|---|---|---|---|---|---|
| 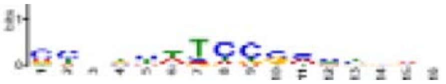 | uniprobe mouse | UP00013_2 | Gabpa_secondary | 1.31e-11 | 7.77e-9 |
| 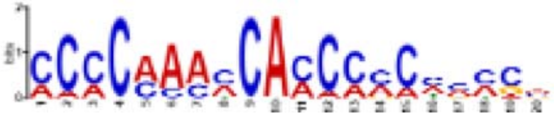 | JASPAR CORE 2014 vertebrates | MA0073.1 | RREB1 | 1.52e-11 | 8.96e-9 |
| 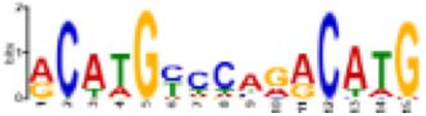 | JASPAR CORE 2014 vertebrates | MA0106.2 | TP53 | 2.08e-11 | 1.23e-8 |
| 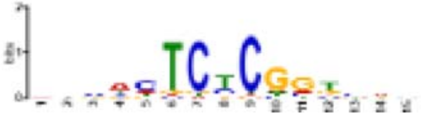 | uniprobe mouse | UP00011_2 | Irf6_secondary | 2.24e-11 | 1.33e-8 |
| 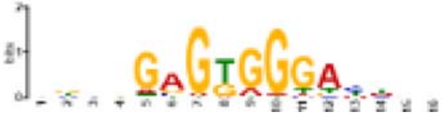 | uniprobe mouse | UP00007_2 | Egr1_secondary | 2.41e-11 | 1.42e-8 |
| 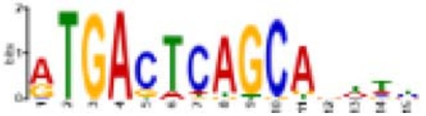 | JASPAR CORE 2014 vertebrates | MA0501.1 | NFE2::MAF | 5.34e-11 | 3.15e-8 |
| 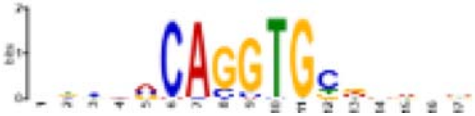 | uniprobe mouse | UP00046_1 | Tcfe2a_primary | 6.51e-11 | 3.85e-8 |
| 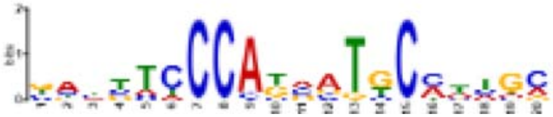 | JASPAR CORE 2014 vertebrates | MA0088.1 | znf143 | 9.75e-11 | 5.76e-8 |
| 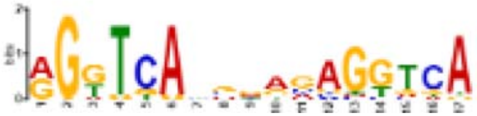 | JASPAR CORE 2014 vertebrates | MA0159.1 | RXR::RAR_DR5 | 1.07e-10 | 6.31e-8 |
| 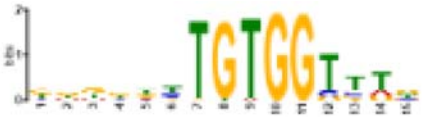 | JASPAR CORE 2014 vertebrates | MA0511.1 | RUNX2 | 1.34e-10 | 7.92e-8 |
| 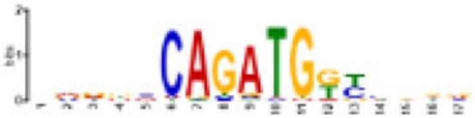 | uniprobe mouse | UP00046_2 | Tcfe2a_secondary | 2.45e-10 | 1.45e-7 |
| 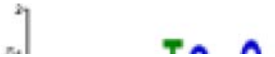 | [illegible] | [illegible] | [illegible] | 2.74e-[illegible] | [illegible] |

| | | | | | |
|---|---|---|---|---|---|
| 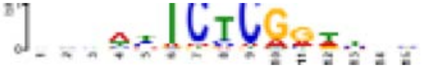 | uniprobe mouse | UP00018_2 | Irf4_secondary | [illegible] 10 | 1.62e-7 |
| 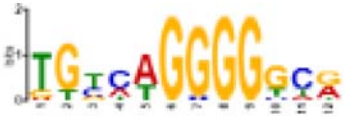 | JASPAR CORE 2014 vertebrates | MA0155.1 | INSM1 | 8.65e-10 | 5.11e-7 |
| 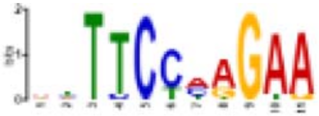 | JASPAR CORE 2014 vertebrates | MA0519.1 | Stat5a::Stat5b | 1.85e-9 | 1.09e-6 |
| 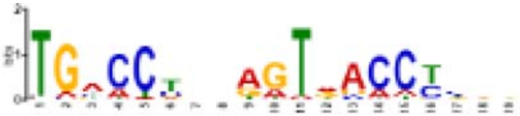 | JASPAR CORE 2014 vertebrates | MA0494.1 | Nr1h3::Rxra | 1.90e-9 | 1.12e-6 |
| 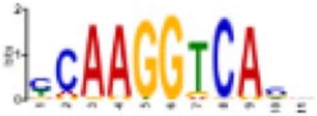 | JASPAR CORE 2014 vertebrates | MA0592.1 | ESRRA | 2.75e-9 | 1.63e-6 |
| 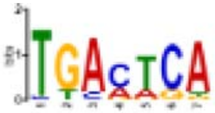 | JASPAR CORE 2014 vertebrates | MA0099.2 | JUN::FOS | 4.23e-9 | 2.50e-6 |
| 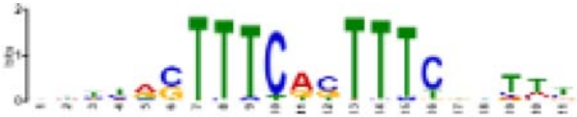 | JASPAR CORE 2014 vertebrates | MA0050.2 | IRF1 | 4.48e-9 | 2.65e-6 |
| 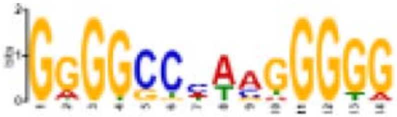 | JASPAR CORE 2014 vertebrates | MA0163.1 | PLAG1 | 5.21e-9 | 3.08e-6 |
| 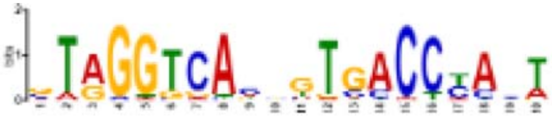 | JASPAR CORE 2014 vertebrates | MA0066.1 | PPARG | 5.36e-9 | 3.17e-6 |
| 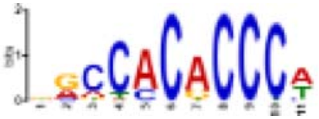 | JASPAR CORE 2014 vertebrates | MA0493.1 | Klf1 | 8.84e-9 | 5.22e-6 |
| 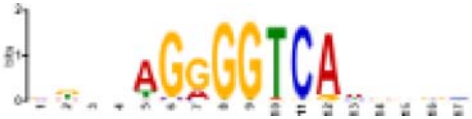 | uniprobe mouse | UP00079_2 | Esrra_secondary | 1.33e-8 | 7.89e-6 |
| 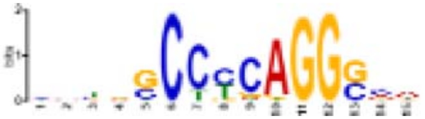 | JASPAR CORE 2014 vertebrates | MA0524.1 | TFAP2C | 1.53e-8 | 9.04e-6 |
| 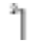 | | | | [illegible] | |

| | | | | | |
|---|---|---|---|---|---|
| 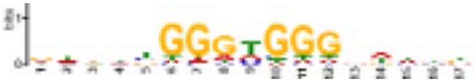 | uniprobe mouse | UP00096_2 | Sox13_secondary | 1.55e-8 | 9.14e-6 |
| 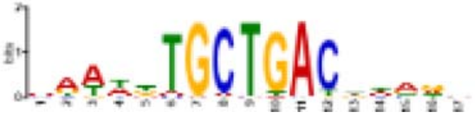 | uniprobe mouse | UP00045_1 | Mafb_primary | 1.78e-8 | 1.05e-5 |
| 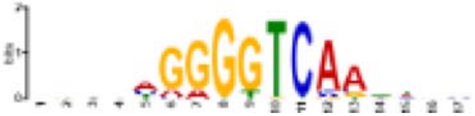 | uniprobe mouse | UP00066_1 | Hnf4a_primary | 2.74e-8 | 1.62e-5 |
| 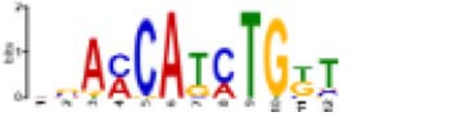 | JASPAR CORE 2014 vertebrates | MA0091.1 | TAL1::TCF3 | 6.58e-8 | 3.89e-5 |
| 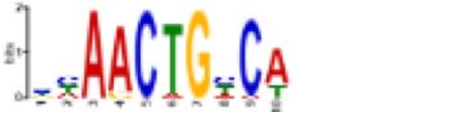 | JASPAR CORE 2014 vertebrates | MA0100.2 | Myb | 6.93e-8 | 4.09e-5 |
| 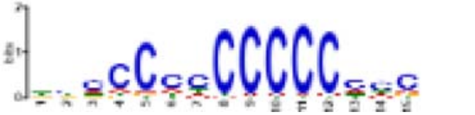 | uniprobe mouse | UP00021_1 | Zfp281_primary | 7.56e-8 | 4.47e-5 |
| 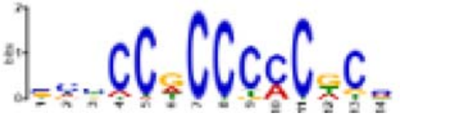 | JASPAR CORE 2014 vertebrates | MA0162.2 | EGR1 | 1.52e-7 | 9.00e-5 |
| 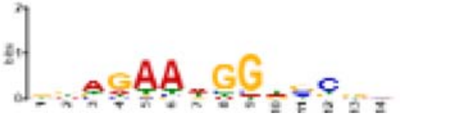 | uniprobe mouse | UP00086_2 | Irf3_secondary | 2.02e-7 | 1.19e-4 |
| 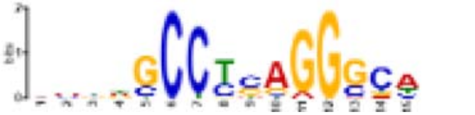 | JASPAR CORE 2014 vertebrates | MA0003.2 | TFAP2A | 3.05e-7 | 1.80e-4 |
| 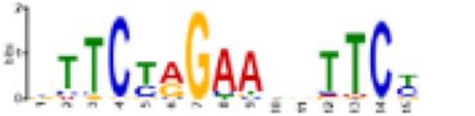 | JASPAR CORE 2014 vertebrates | MA0486.1 | HSF1 | 3.06e-7 | 1.81e-4 |
| 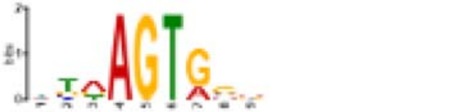 | JASPAR CORE 2014 vertebrates | MA0122.1 | Nkx3-2 | 3.15e-7 | 1.86e-4 |
| 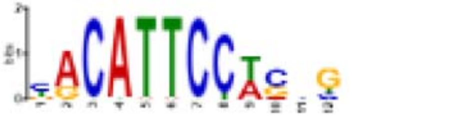 | JASPAR CORE 2014 vertebrates | MA0090.1 | TEAD1 | 3.87e-7 | 2.29e-4 |

| | | | | | |
|---|---|---|---|---|---|
| 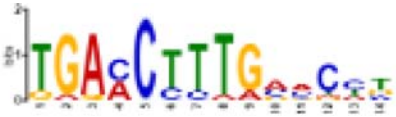 | JASPAR CORE 2014 vertebrates | MA0017.1 | NR2F1 | 3.98e-7 | 2.35e-4 |
| 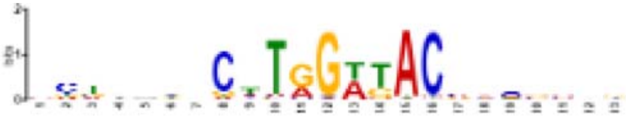 | uniprobe mouse | UP00098_2 | Rfx3_secondary | 4.21e-7 | 2.49e-4 |
| 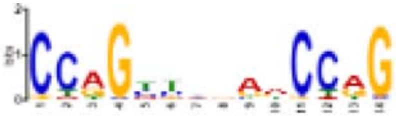 | JASPAR CORE 2014 vertebrates | MA0145.2 | Tcfcp2l1 | 4.44e-7 | 2.63e-4 |
| 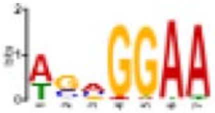 | JASPAR CORE 2014 vertebrates | MA0081.1 | SPIB | 9.82e-7 | 5.80e-4 |
| 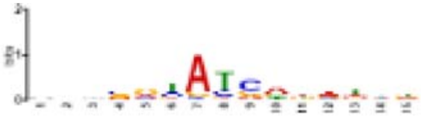 | uniprobe mouse | UP00078_2 | Arid3a_secondary | 1.04e-6 | 6.15e-4 |
| 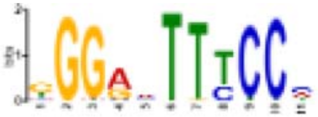 | JASPAR CORE 2014 vertebrates | MA0105.3 | NFKB1 | 1.12e-6 | 6.64e-4 |
| 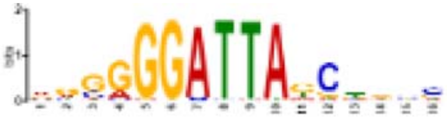 | uniprobe mouse | UP00265_1 | Pitx3_3497.2 | 1.12e-6 | 6.64e-4 |
| 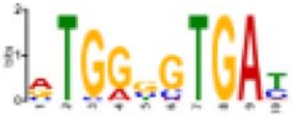 | JASPAR CORE 2014 vertebrates | MA0596.1 | SREBF2 | 1.50e-6 | 8.86e-4 |
| 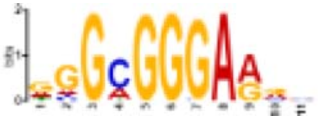 | JASPAR CORE 2014 vertebrates | MA0471.1 | E2F6 | 1.59e-6 | 9.40e-4 |
| 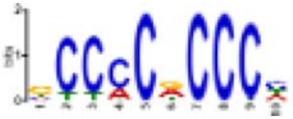 | JASPAR CORE 2014 vertebrates | MA0599.1 | KLF5 | 2.11e-6 | 1.25e-3 |
| 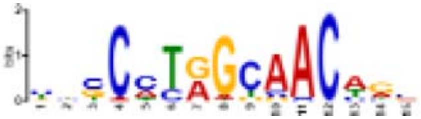 | JASPAR CORE 2014 vertebrates | MA0510.1 | RFX5 | 2.55e-6 | 1.50e-3 |
| 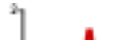 | [illegible] | [illegible] | [illegible] | [illegible] | [illegible] |

| | | | | | |
|---|---|---|---|---|---|
| 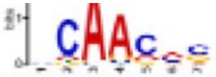 | JASPAR CORE 2014 vertebrates | MA0133.1 | BRCA1 | 2.84e-6 | 1.68e-3 |
| 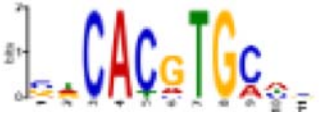 | JASPAR CORE 2014 vertebrates | MA0464.1 | Bhlhe40 | 3.16e-6 | 1.87e-3 |
| 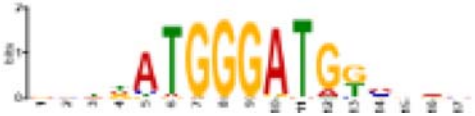 | uniprobe mouse | UP00033_1 | Zfp410_primary | 3.16e-6 | 1.87e-3 |
| 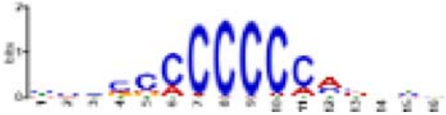 | uniprobe mouse | UP00022_1 | Zfp740_primary | 3.25e-6 | 1.92e-3 |
| 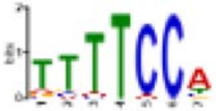 | JASPAR CORE 2014 vertebrates | MA0152.1 | NFATC2 | 4.70e-6 | 2.77e-3 |
| 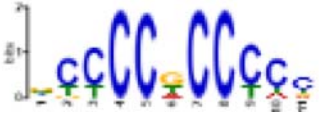 | JASPAR CORE 2014 vertebrates | MA0079.3 | SP1 | 5.91e-6 | 3.48e-3 |
| 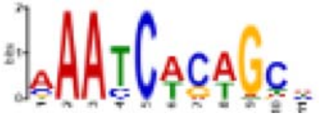 | JASPAR CORE 2014 vertebrates | MA0483.1 | Gfi1b | 6.30e-6 | 3.72e-3 |
| 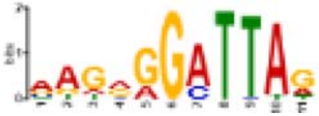 | JASPAR CORE 2014 vertebrates | MA0467.1 | Crx | 6.34e-6 | 3.74e-3 |
| 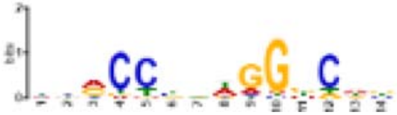 | uniprobe mouse | UP00005_2 | Tcfap2a_secondary | 7.82e-6 | 4.61e-3 |
| 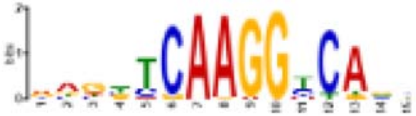 | JASPAR CORE 2014 vertebrates | MA0505.1 | Nr5a2 | 8.75e-6 | 5.16e-3 |
| 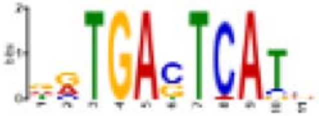 | JASPAR CORE 2014 vertebrates | MA0491.1 | JUND | 1.15e-5 | 6.79e-3 |
| 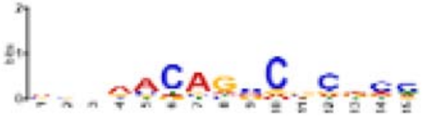 | uniprobe mouse | UP00036_2 | Myf6_secondary | 1.92e-5 | 1.13e-2 |

| | | | | | |
|---|---|---|---|---|---|
| 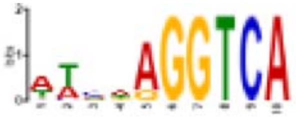 | JASPAR CORE 2014 vertebrates | MA0071.1 | RORA_1 | 1.97e-5 | 1.16e-2 |
| 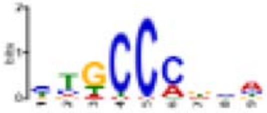 | JASPAR CORE 2014 vertebrates | MA0597.1 | THAP1 | 2.30e-5 | 1.35e-2 |
| 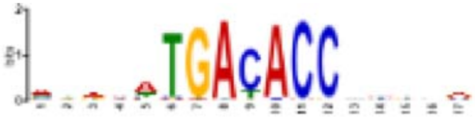 | uniprobe mouse | UP00199_1 | Six4_2860.1 | 2.30e-5 | 1.35e-2 |
| 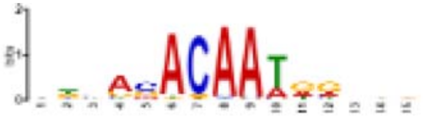 | uniprobe mouse | UP00004_2 | Sox14_secondary | 3.00e-5 | 1.76e-2 |
| 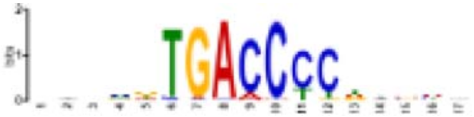 | uniprobe mouse | UP00053_1 | Rxra_primary | 3.16e-5 | 1.85e-2 |
| 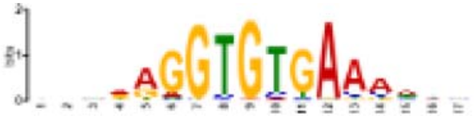 | uniprobe mouse | UP00068_1 | Eomes_primary | 3.83e-5 | 2.24e-2 |
| 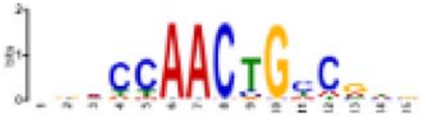 | uniprobe mouse | UP00081_2 | Mybl1_secondary | 4.28e-5 | 2.50e-2 |
| 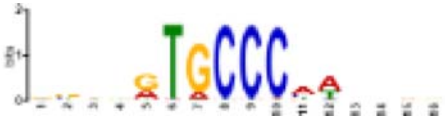 | uniprobe mouse | UP00035_2 | Hic1_secondary | 4.69e-5 | 2.74e-2 |
| 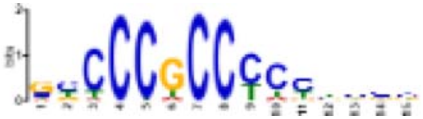 | JASPAR CORE 2014 vertebrates | MA0516.1 | SP2 | 5.08e-5 | 2.96e-2 |
| 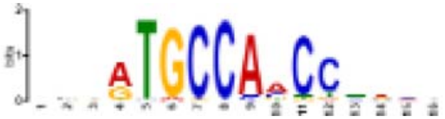 | uniprobe mouse | UP00035_1 | Hic1_primary | 6.17e-5 | 3.58e-2 |
| 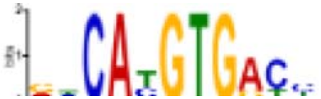 | JASPAR CORE 2014 vertebrates | MA0526.1 | USF2 | 8.05e-5 | 4.65e-2 |

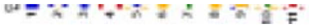

| | | | | | |
|---|---|---|---|---|---|
| 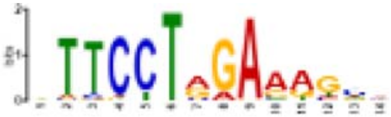 | JASPAR CORE 2014 vertebrates | MA0463.1 | Bcl6 | 8.15e-5 | 4.70e-2 |

### Input Files

#### Sequences

| Primary Sequences | Number | Control Sequences | Number | | | | |
|---|---|---|---|---|---|---|---|
| Control Sequences | Number | | | | | | |
| ud300nonexpr.fa | 15480 | ud300expr.fa | 6525 | ud300nonexpr.fa | 15480 | ud300expr.fa | 6525 |

#### Motifs

| Database | Source | Motif Count |
|---|---|---|
| JASPAR CORE 2014 vertebrates | db/JASPAR/JASPAR_CORE_2014_vertebrates.meme | 205 |
| uniprobe mouse | db/MOUSE/uniprobe_mouse.meme | 386 |
| JASPAR CORE 2014 vertebrates | db/JASPAR/JASPAR_CORE_2014_vertebrates.meme | 205 |
| uniprobe mouse | db/MOUSE/uniprobe_mouse.meme | 386 |

**AME version**
4.11.24.11.2 (Release date: Thu May 05 14:58:55 2016 -0700Thu May 05 14:58:55 2016 -0700)
Copyright © Robert McLeay & Timothy Bailey, 2009.

**Command line summary**

```
/home/meme/meme_4.11.2/bin/ame --verbose 1 --oc . --control ud300expr.fa --bgformat 2 --bgfile background.model --scoring avg --method ranksum --pvalue-report-threshold 0.05 ud300nonexpr.fa db/JASPAR/JASPAR_CORE_2014_vertebrates.meme db/MOUSE/uniprobe_mouse.meme
```

Non-expressed downstream

### AME
Analysis of Motif Enrichment

For further information on how to interpret these results or to get a copy of the MEME software please access http://meme-suite.org.

If you use AME in your research, please cite the following paper:
Robert McLeay and Timothy L. Bailey, "Motif Enrichment Analysis: A unified framework and method evaluation", *BMC Bioinformatics*, **11**:165, 2010, doi:10.1186/1471-2105-11-165. [full text]

Enriched Motifs | Input Files | Program information

#### Enriched Motifs

Fixed partition size: number of primary sequences (15480)

Sequence motif score: avg_odds
Background model source: file background.model
Background model frequencies: 0.291,0.209,0.209,0.291
Total pseudocount added to a motif column: 0.25

Statistical test: Wilcoxon rank-sum test
Ranksum method: quick
Threshold *p*-value for reporting results: 0.05
Number of multiple tests for Bonferroni correction: #Motifs × #PartitionsTested = 591 × 1 = 591Fixed partition size: number of primary sequences (15480)

Sequence motif score: avg_odds
Background model source: file background.model
Background model frequencies: 0.291,0.209,0.209,0.291
Total pseudocount added to a motif column: 0.25

Statistical test: Wilcoxon rank-sum test
Ranksum method: quick
Threshold *p*-value for reporting results: 0.05
Number of multiple tests for Bonferroni correction: #Motifs × #PartitionsTested = 591 × 1 = 591

| Logo | Database | ID | Name | *p*-value | Adjusted *p*-value |
|---|---|---|---|---|---|
| 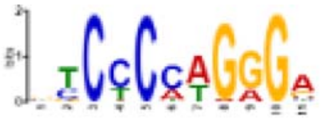 | JASPAR CORE 2014 vertebrates | MA0154.2 | EBF1 | 1.65e-33 | 9.78e-31 |

| | | | | | |
|---|---|---|---|---|---|
| 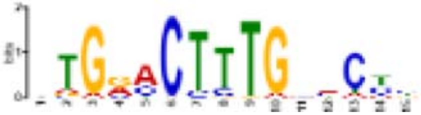 | JASPAR CORE 2014 vertebrates | MA0114.2 | HNF4A | 8.48e-21 | 5.01e-18 |
| 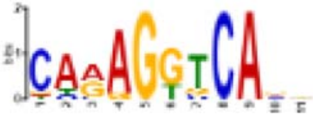 | JASPAR CORE 2014 vertebrates | MA0512.1 | Rxra | 2.57e-20 | 1.52e-17 |
| 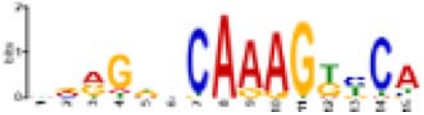 | JASPAR CORE 2014 vertebrates | MA0484.1 | HNF4G | 6.17e-20 | 3.65e-17 |
| 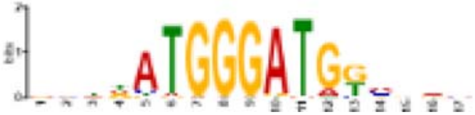 | uniprobe mouse | UP00033_1 | Zfp410_primary | 3.57e-19 | 2.11e-16 |
| 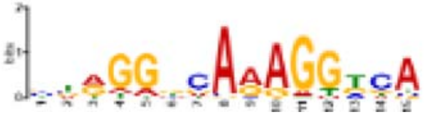 | JASPAR CORE 2014 vertebrates | MA0065.2 | PPARG::RXRA | 4.61e-16 | 2.72e-13 |
| 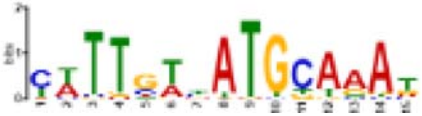 | JASPAR CORE 2014 vertebrates | MA0142.1 | Pou5f1::Sox2 | 7.38e-16 | 4.36e-13 |
| 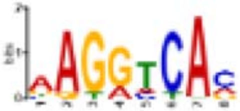 | JASPAR CORE 2014 vertebrates | MA0160.1 | NR4A2 | 1.55e-15 | 9.15e-13 |
| 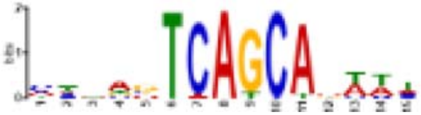 | JASPAR CORE 2014 vertebrates | MA0496.1 | MAFK | 6.76e-15 | 3.99e-12 |
| 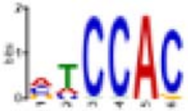 | JASPAR CORE 2014 vertebrates | MA0130.1 | ZNF354C | 9.21e-14 | 5.45e-11 |
| 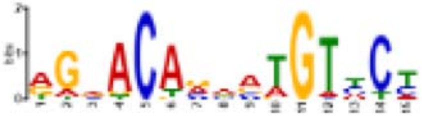 | JASPAR CORE 2014 vertebrates | MA0113.2 | NR3C1 | 1.27e-13 | 7.52e-11 |
| 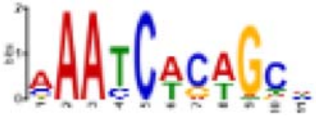 | JASPAR CORE 2014 vertebrates | MA0483.1 | Gfi1b | 1.73e-13 | 1.02e-10 |

| | | | | | |
|---|---|---|---|---|---|
| 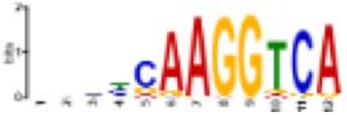 | JASPAR CORE 2014 vertebrates | MA0141.2 | Esrrb | 2.24e-13 | 1.32e-10 |
| 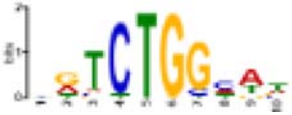 | JASPAR CORE 2014 vertebrates | MA0092.1 | Hand1::Tcfe2a | 6.44e-13 | 3.81e-10 |
| 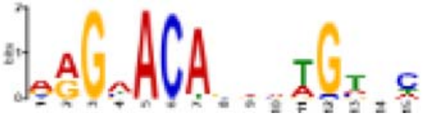 | JASPAR CORE 2014 vertebrates | MA0007.2 | AR | 6.64e-13 | 3.93e-10 |
| 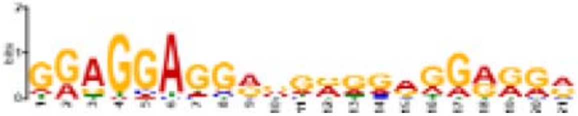 | JASPAR CORE 2014 vertebrates | MA0528.1 | ZNF263 | 1.22e-12 | 7.20e-10 |
| 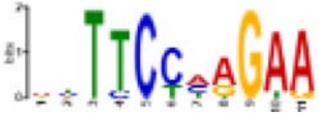 | JASPAR CORE 2014 vertebrates | MA0519.1 | Stat5a::Stat5b | 1.94e-12 | 1.14e-9 |
| 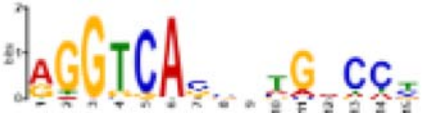 | JASPAR CORE 2014 vertebrates | MA0258.2 | ESR2 | 2.15e-12 | 1.27e-9 |
| 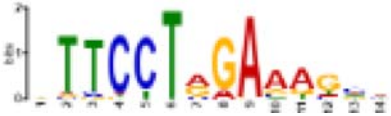 | JASPAR CORE 2014 vertebrates | MA0463.1 | Bcl6 | 2.31e-11 | 1.37e-8 |
| 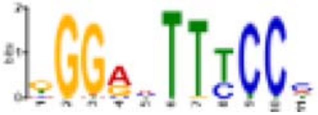 | JASPAR CORE 2014 vertebrates | MA0105.3 | NFKB1 | 5.26e-11 | 3.11e-8 |
| 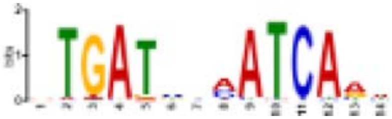 | uniprobe mouse | UP00198_1 | Cphx_3484.1 | 1.02e-10 | 6.04e-8 |
| 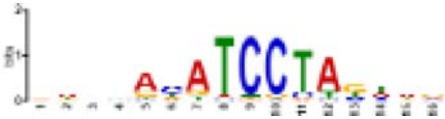 | uniprobe mouse | UP00038_2 | Spdef_secondary | 1.15e-10 | 6.82e-8 |
| 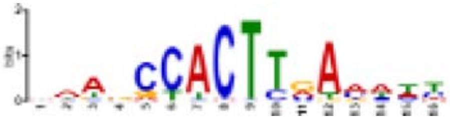 | uniprobe mouse | UP00107_1 | Nkx2-4_3074.1 | 1.44e-10 | 8.52e-8 |

| | | | | | |
|---|---|---|---|---|---|
| 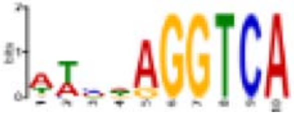 | JASPAR CORE 2014 vertebrates | MA0071.1 | RORA_1 | 1.49e-10 | 8.78e-8 |
| 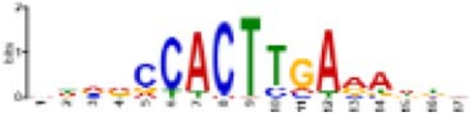 | uniprobe mouse | UP00231_1 | Nkx2-2_2823.1 | 2.18e-10 | 1.29e-7 |
| 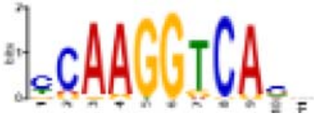 | JASPAR CORE 2014 vertebrates | MA0592.1 | ESRRA | 2.66e-10 | 1.57e-7 |
| 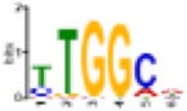 | JASPAR CORE 2014 vertebrates | MA0161.1 | NFIC | 3.29e-10 | 1.94e-7 |
| 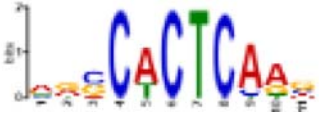 | JASPAR CORE 2014 vertebrates | MA0503.1 | Nkx2-5 | 3.47e-10 | 2.05e-7 |
| 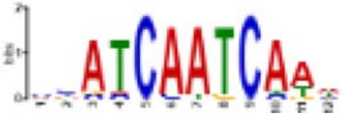 | JASPAR CORE 2014 vertebrates | MA0070.1 | PBX1 | 6.82e-10 | 4.03e-7 |
| 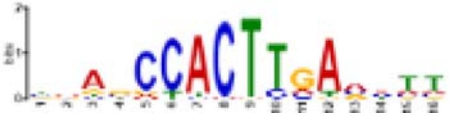 | uniprobe mouse | UP00165_1 | Titf1_1722.2 | 8.86e-10 | 5.24e-7 |
| 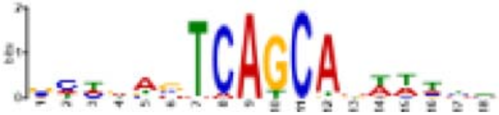 | JASPAR CORE 2014 vertebrates | MA0495.1 | MAFF | 1.30e-9 | 7.66e-7 |
| 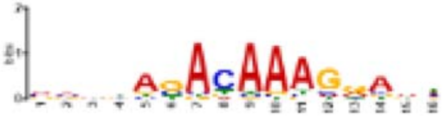 | uniprobe mouse | UP00101_2 | Sox12_secondary | 1.53e-9 | 9.06e-7 |
| 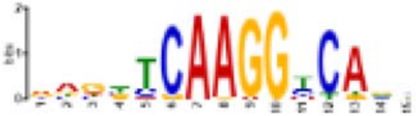 | JASPAR CORE 2014 vertebrates | MA0505.1 | Nr5a2 | 2.14e-9 | 1.27e-6 |
| 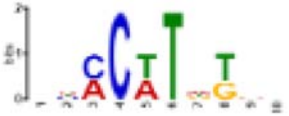 | JASPAR CORE 2014 vertebrates | MA0109.1 | Hltf | 6.54e-9 | 3.87e-6 |
| 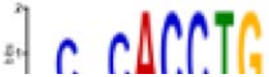 | JASPAR CORE 2014 vertebrates | MA0103.2 | ZEB1 | 7.02e-9 | 4.15e-6 |

| | | | | | |
|---|---|---|---|---|---|
| 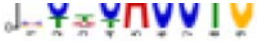 | vertebrates | | | | |
| 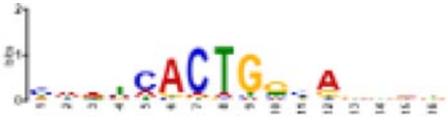 | uniprobe mouse | UP00031_2 | Zbtb3_secondary | 1.21e-8 | 7.17e-6 |
| 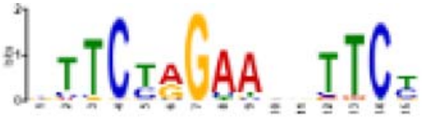 | JASPAR CORE 2014 vertebrates | MA0486.1 | HSF1 | 1.87e-8 | 1.10e-5 |
| 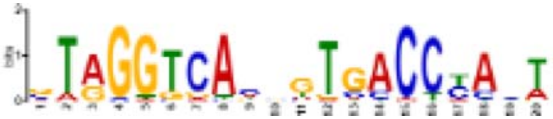 | JASPAR CORE 2014 vertebrates | MA0066.1 | PPARG | 2.44e-8 | 1.44e-5 |
| 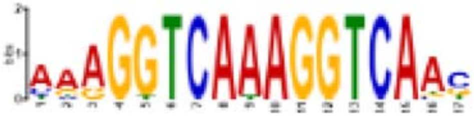 | JASPAR CORE 2014 vertebrates | MA0115.1 | NR1H2::RXRA | 2.89e-8 | 1.71e-5 |
| 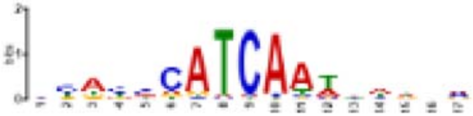 | uniprobe mouse | UP00185_1 | Pbx1_3203.1 | 3.69e-8 | 2.18e-5 |
| 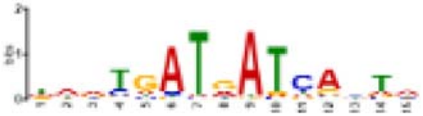 | uniprobe mouse | UP00219_2 | Cutl1_3494.2 | 3.91e-8 | 2.31e-5 |
| 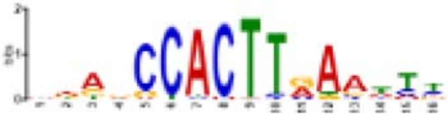 | uniprobe mouse | UP00249_1 | Nkx2-5_3436.1 | 5.24e-8 | 3.10e-5 |
| 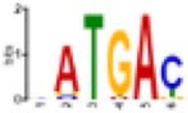 | JASPAR CORE 2014 vertebrates | MA0089.1 | NFE2L1::MafG | 9.75e-8 | 5.76e-5 |
| 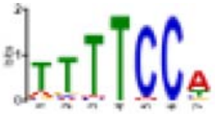 | JASPAR CORE 2014 vertebrates | MA0152.1 | NFATC2 | 1.02e-7 | 6.03e-5 |
| 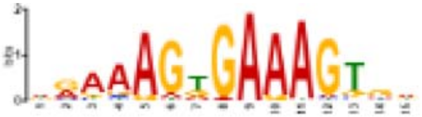 | JASPAR CORE 2014 vertebrates | MA0508.1 | PRDM1 | 1.09e-7 | 6.42e-5 |
| 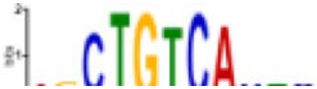 | JASPAR CORE 2014 vertebrates | MA0498.1 | Meis1 | 1.49e-7 | 8.81e-5 |

| | | | | | |
|---|---|---|---|---|---|
| 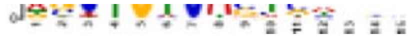 | [illegible] | | | | |
| 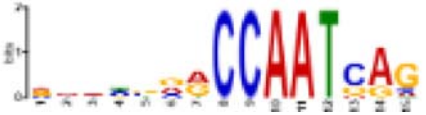 | JASPAR CORE 2014 vertebrates | MA0502.1 | NFYB | 2.97e-7 | 1.76e-4 |
| 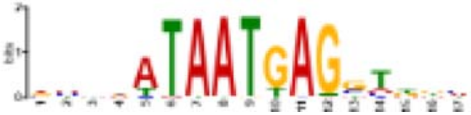 | uniprobe mouse | UP00146_2 | Pou6f1_3733.1 | 4.36e-7 | 2.57e-4 |
| 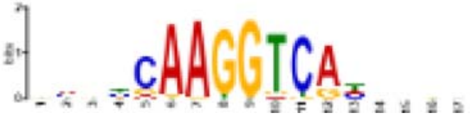 | uniprobe mouse | UP00079_1 | Esrra_primary | 6.52e-7 | 3.85e-4 |
| 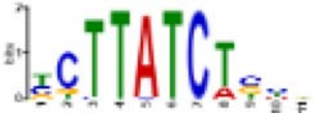 | JASPAR CORE 2014 vertebrates | MA0482.1 | Gata4 | 1.48e-6 | 8.71e-4 |
| 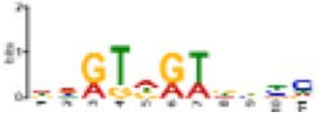 | JASPAR CORE 2014 vertebrates | MA0027.1 | En1 | 2.03e-6 | 1.20e-3 |
| 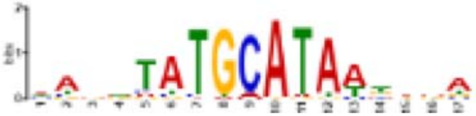 | uniprobe mouse | UP00211_1 | Pou3f3_3235.2 | 2.26e-6 | 1.34e-3 |
| 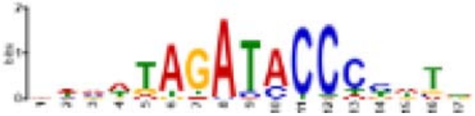 | uniprobe mouse | UP00232_1 | Dobox4_3956.2 | 2.29e-6 | 1.35e-3 |
| 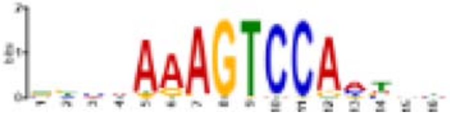 | uniprobe mouse | UP00066_2 | Hnf4a_secondary | 2.70e-6 | 1.59e-3 |
| 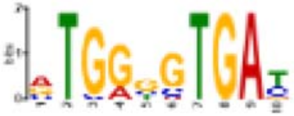 | JASPAR CORE 2014 vertebrates | MA0596.1 | SREBF2 | 3.28e-6 | 1.94e-3 |
| 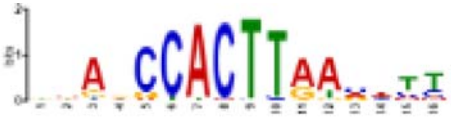 | uniprobe mouse | UP00147_1 | Nkx2-6_3437.1 | 3.92e-6 | 2.32e-3 |
| 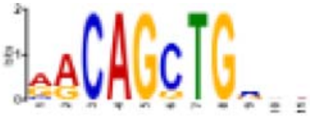 | JASPAR CORE 2014 vertebrates | MA0521.1 | Tcf12 | 4.76e-6 | 2.81e-3 |

| | | | | | |
|---|---|---|---|---|---|
| 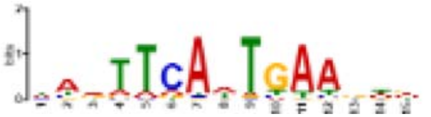 | uniprobe mouse | UP00012_1 | Bbx_primary | 6.71e-6 | 3.96e-3 |
| 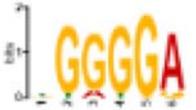 | JASPAR CORE 2014 vertebrates | MA0056.1 | MZF1_1-4 | 8.07e-6 | 4.76e-3 |
| 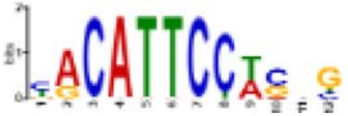 | JASPAR CORE 2014 vertebrates | MA0090.1 | TEAD1 | 1.61e-5 | 9.49e-3 |
| 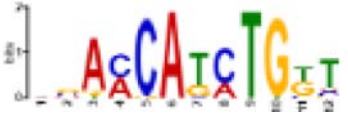 | JASPAR CORE 2014 vertebrates | MA0091.1 | TAL1::TCF3 | 1.83e-5 | 1.08e-2 |
| 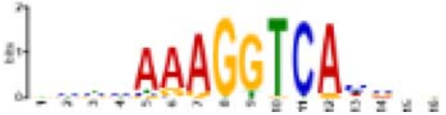 | uniprobe mouse | UP00048_1 | Rara_primary | 1.97e-5 | 1.16e-2 |
| 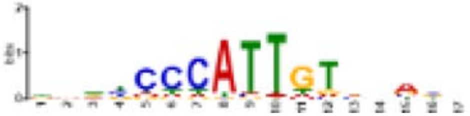 | uniprobe mouse | UP00055_2 | Hbp1_secondary | 2.17e-5 | 1.28e-2 |
| 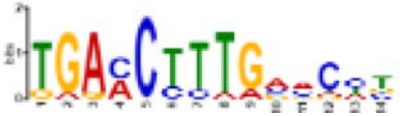 | JASPAR CORE 2014 vertebrates | MA0017.1 | NR2F1 | 3.15e-5 | 1.84e-2 |
| 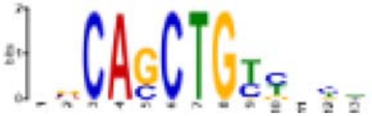 | JASPAR CORE 2014 vertebrates | MA0499.1 | Myod1 | 3.24e-5 | 1.90e-2 |
| 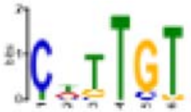 | JASPAR CORE 2014 vertebrates | MA0442.1 | SOX10 | 3.71e-5 | 2.17e-2 |
| 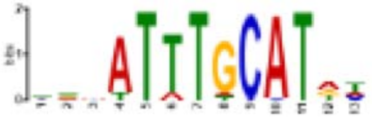 | JASPAR CORE 2014 vertebrates | MA0507.1 | POU2F2 | 4.20e-5 | 2.45e-2 |
| 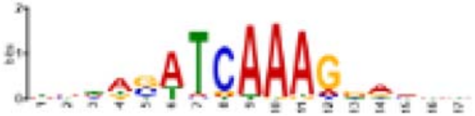 | uniprobe mouse | UP00058_1 | Tcf3_primary | 4.60e-5 | 2.68e-2 |
| 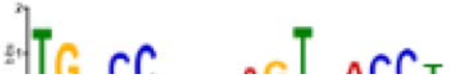 | JASPAR CORE 2014 vertebrates | MA0494.1 | Nr1h3::Rxra | 6.04e-5 | 3.51e-2 |

| | | | | | |
|---|---|---|---|---|---|
| 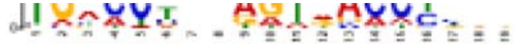 | vertebrates | | | | |
| 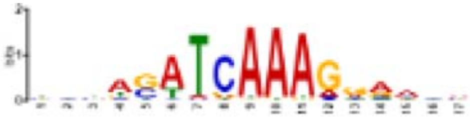 | uniprobe mouse | UP00054_1 | Tcf7_primary | 6.83e-5 | 3.95e-2 |
| 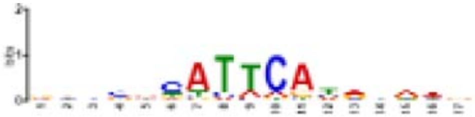 | uniprobe mouse | UP00014_2 | Sox17_secondary | 7.06e-5 | 4.09e-2 |

### Input Files

#### Sequences

| Primary Sequences | Number | Control Sequences | Number | | | | |
|---|---|---|---|---|---|---|---|
| Control Sequences | Number | | | | | | |
| ud-300nonexpr.fa | 15480 | ud-300expr.fa | 6525 | ud-300nonexpr.fa | 15480 | ud-300expr.fa | 6525 |

#### Motifs

| Database | Source | Motif Count |
|---|---|---|
| JASPAR CORE 2014 vertebrates | db/JASPAR/JASPAR_CORE_2014_vertebrates.meme | 205 |
| uniprobe mouse | db/MOUSE/uniprobe_mouse.meme | 386 |
| JASPAR CORE 2014 vertebrates | db/JASPAR/JASPAR_CORE_2014_vertebrates.meme | 205 |
| uniprobe mouse | db/MOUSE/uniprobe_mouse.meme | 386 |

**AME version**
4.11.24.11.2 (Release date: Thu May 05 14:58:55 2016 -0700Thu May 05 14:58:55 2016 -0700)
Copyright © Robert McLeay & Timothy Bailey, 2009.

**Command line summary**

```
/home/meme/meme_4.11.2/bin/ame --verbose 1 --oc . --control ud-300expr.fa --bgformat 2 --bgfile background.model --scoring avg --method ranksum --pvalue-report-threshold 0.05 ud-300nonexpr.fa db/JASPAR/JASPAR_CORE_2014_vertebrates.meme db/MOUSE/uniprobe_mouse.meme
```

Expressed upstream

### AME
#### Analysis of Motif Enrichment

For further information on how to interpret these results or to get a copy of the MEME software please access http://meme-suite.org.

If you use AME in your research, please cite the following paper:
Robert McLeay and Timothy L. Bailey, "Motif Enrichment Analysis: A unified framework and method evaluation", *BMC Bioinformatics*, **11**:165, 2010, doi:10.1186/1471-2105-11-165. [full text]

Enriched Motifs | Input Files | Program information

#### Enriched Motifs

Fixed partition size: number of primary sequences (6525)

Sequence motif score: avg_odds
Background model source: file background.model
Background model frequencies: 0.291,0.209,0.209,0.291
Total pseudocount added to a motif column: 0.25

Statistical test: Wilcoxon rank-sum test
Ranksum method: quick
Threshold *p*-value for reporting results: 0.05
Number of multiple tests for Bonferroni correction: #Motifs × #PartitionsTested = 591 × 1 = 591Fixed partition size: number of primary sequences (6525)

Sequence motif score: avg_odds
Background model source: file background.model
Background model frequencies: 0.291,0.209,0.209,0.291
Total pseudocount added to a motif column: 0.25

Statistical test: Wilcoxon rank-sum test
Ranksum method: quick
Threshold *p*-value for reporting results: 0.05
Number of multiple tests for Bonferroni correction: #Motifs × #PartitionsTested = 591 × 1 = 591

| Logo | Database | ID | Name | *p*-value | Adjusted *p*-value |
|---|---|---|---|---|---|
| 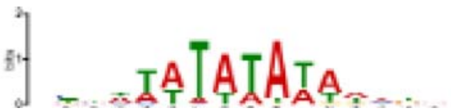 | uniprobe mouse | UP00029_1 | Tbp_primary | 2.04e-61 | 1.21e-58 |

| | | | | | |
|---|---|---|---|---|---|
| 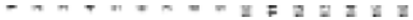 | | | | | |
| 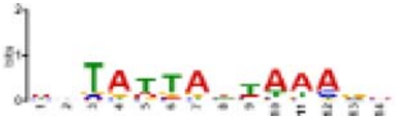 | uniprobe mouse | UP00024_2 | Glis2_secondary | 5.41e-54 | 3.20e-51 |
| 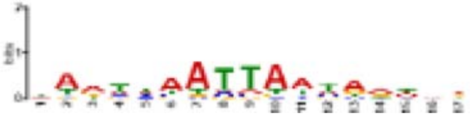 | uniprobe mouse | UP00255_1 | Dbx1_3486.1 | 2.63e-52 | 1.55e-49 |
| 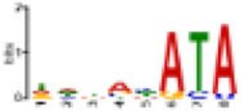 | JASPAR CORE 2014 vertebrates | MA0033.1 | FOXL1 | 3.94e-52 | 2.33e-49 |
| 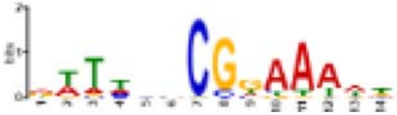 | uniprobe mouse | UP00049_1 | Sp100_primary | 6.37e-52 | 3.77e-49 |
| 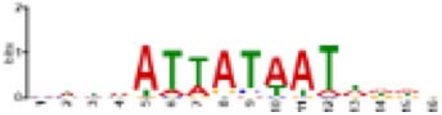 | uniprobe mouse | UP00071_1 | Sox21_primary | 8.77e-46 | 5.18e-43 |
| 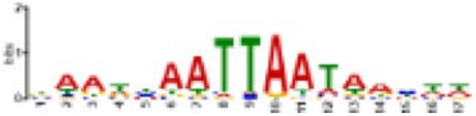 | uniprobe mouse | UP00244_1 | Tlx2_3498.2 | 4.39e-43 | 2.60e-40 |
| 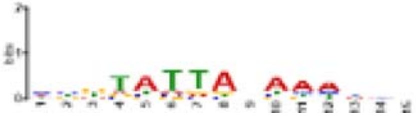 | uniprobe mouse | UP00054_2 | Tcf7_secondary | 1.47e-41 | 8.66e-39 |
| 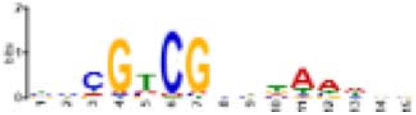 | uniprobe mouse | UP00049_2 | Sp100_secondary | 6.22e-39 | 3.68e-36 |
| 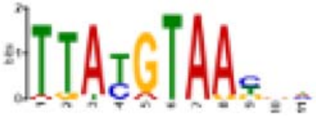 | JASPAR CORE 2014 vertebrates | MA0025.1 | NFIL3 | 2.86e-38 | 1.69e-35 |
| 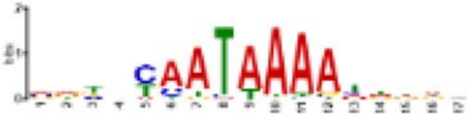 | uniprobe mouse | UP00121_1 | Hoxd10_2368.2 | 4.86e-37 | 2.87e-34 |
| 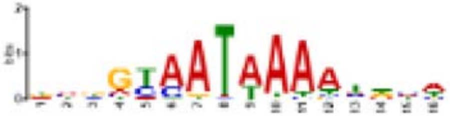 | uniprobe mouse | UP00217_1 | Hoxa10_2318.1 | 6.65e-36 | 3.93e-33 |

| | | | | | |
|---|---|---|---|---|---|
| 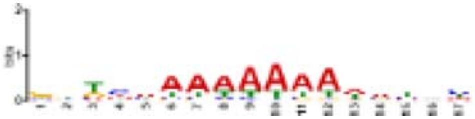 | uniprobe mouse | UP00090_2 | Elf3_secondary | 7.45e-36 | 4.40e-33 |
| 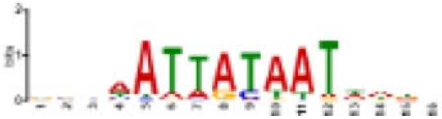 | uniprobe mouse | UP00004_1 | Sox14_primary | 7.87e-36 | 4.65e-33 |
| 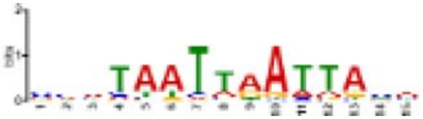 | uniprobe mouse | UP00225_1 | Hlx1_2350.1 | 3.57e-35 | 2.11e-32 |
| 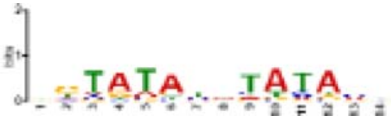 | uniprobe mouse | UP00094_2 | Zfp128_secondary | 5.32e-35 | 3.14e-32 |
| 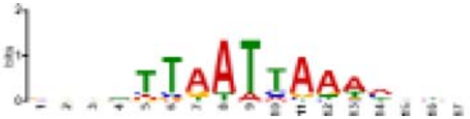 | uniprobe mouse | UP00078_1 | Arid3a_primary | 1.52e-33 | 8.98e-31 |
| 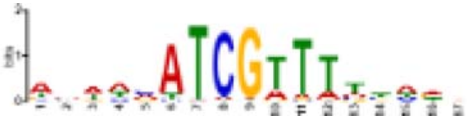 | uniprobe mouse | UP00114_1 | Homez_1063.2 | 9.40e-33 | 5.56e-30 |
| 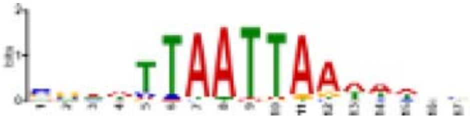 | uniprobe mouse | UP00188_1 | Lmx1a_2238.2 | 3.53e-32 | 2.09e-29 |
| 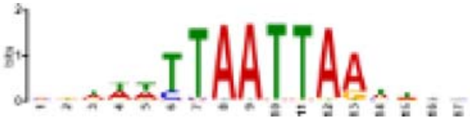 | uniprobe mouse | UP00169_1 | Lmx1b_3433.2 | 2.53e-31 | 1.50e-28 |
| 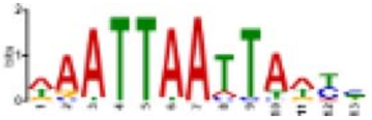 | JASPAR CORE 2014 vertebrates | MA0135.1 | Lhx3 | 2.84e-31 | 1.68e-28 |
| 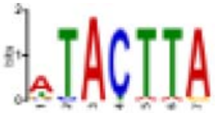 | JASPAR CORE 2014 vertebrates | MA0124.1 | NKX3-1 | 2.98e-31 | 1.76e-28 |
| | JASPAR CORE 2014 [illegible] | [illegible] | [illegible] | 1.08[illegible] | [illegible] |

| | | | | | |
|---|---|---|---|---|---|
| 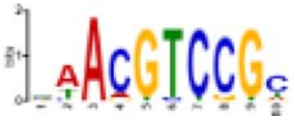 | JASPAR CORE 2014 vertebrates | MA0131.1 | HINFP | 1.08e-30 | 6.36e-28 |
| 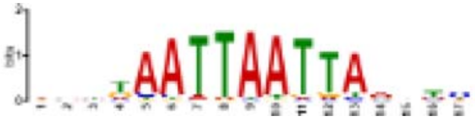 | uniprobe mouse | UP00105_1 | Pou3f4_3773.1 | 1.19e-30 | 7.03e-28 |
| 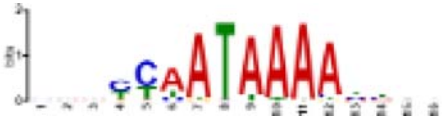 | uniprobe mouse | UP00180_1 | Hoxd13_2356.1 | 2.33e-30 | 1.38e-27 |
| 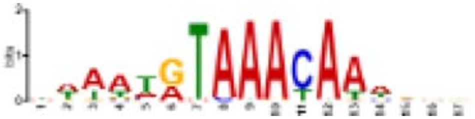 | uniprobe mouse | UP00061_1 | Foxl1_primary | 3.12e-30 | 1.84e-27 |
| 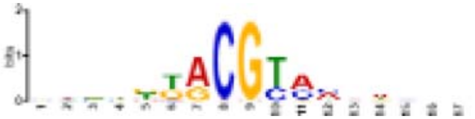 | uniprobe mouse | UP00084_1 | Gmeb1_primary | 3.47e-29 | 2.05e-26 |
| 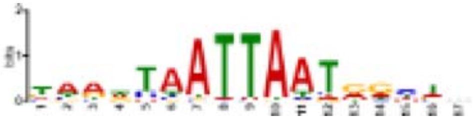 | uniprobe mouse | UP00168_1 | Hoxd8_2644.1 | 1.85e-28 | 1.09e-25 |
| 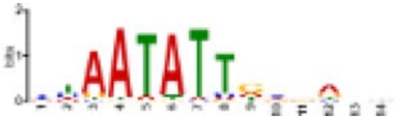 | uniprobe mouse | UP00059_1 | Arid5a_primary | 7.47e-28 | 4.41e-25 |
| 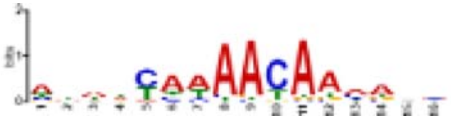 | uniprobe mouse | UP00061_2 | Foxl1_secondary | 2.02e-27 | 1.19e-24 |
| 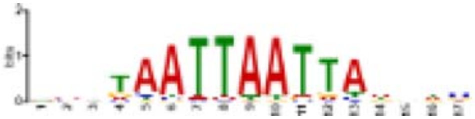 | uniprobe mouse | UP00158_1 | Pou1f1_3818.1 | 2.44e-27 | 1.44e-24 |
| 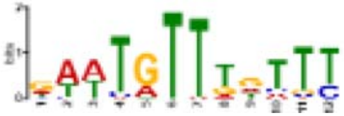 | JASPAR CORE 2014 vertebrates | MA0041.1 | Foxd3 | 2.45e-27 | 1.45e-24 |
| 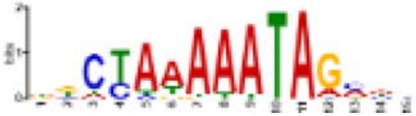 | JASPAR CORE 2014 vertebrates | MA0052.2 | MEF2A | 1.66e-26 | 9.83e-24 |
| 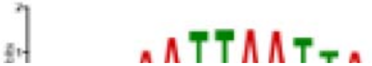 | uniprobe mouse | UP00129_1 | Pou3f1_3819.1 | 3.10e- | 1.83e-23 |

| | | | | | |
|---|---|---|---|---|---|
| 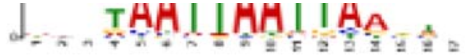 | [illegible] | [illegible] | [illegible] | [illegible] 26 | [illegible] |
| 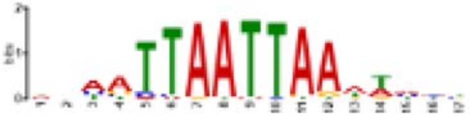 | uniprobe mouse | UP00130_1 | Lhx3_3431.1 | 6.06e-26 | 3.58e-23 |
| 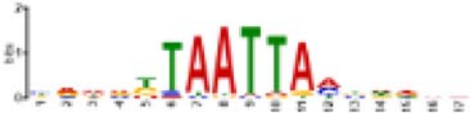 | uniprobe mouse | UP00260_1 | Hoxc6_3954.2 | 1.18e-25 | 6.95e-23 |
| 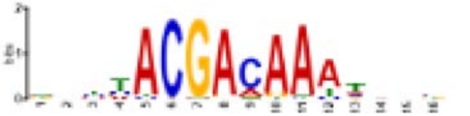 | uniprobe mouse | UP00072_1 | IRC900814_primary | 1.22e-25 | 7.21e-23 |
| 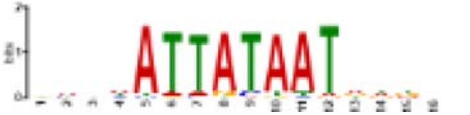 | uniprobe mouse | UP00016_1 | Sry_primary | 2.27e-25 | 1.34e-22 |
| 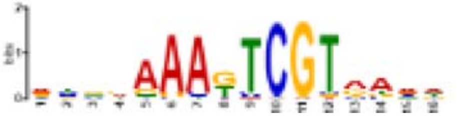 | uniprobe mouse | UP00072_2 | IRC900814_secondary | 2.30e-25 | 1.36e-22 |
| 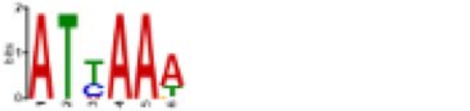 | JASPAR CORE 2014 vertebrates | MA0151.1 | ARID3A | 3.39e-25 | 2.01e-22 |
| 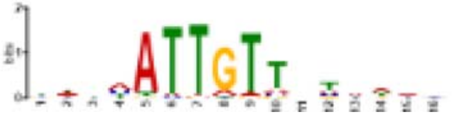 | uniprobe mouse | UP00064_1 | Sox18_primary | 1.44e-24 | 8.49e-22 |
| 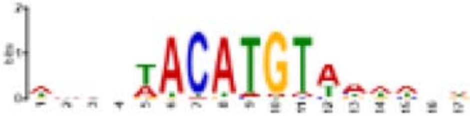 | uniprobe mouse | UP00194_1 | Irx4_2242.3 | 2.41e-24 | 1.42e-21 |
| 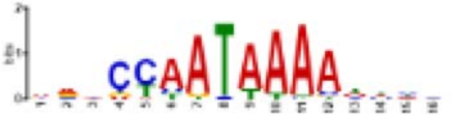 | uniprobe mouse | UP00134_1 | Hoxb13_3479.1 | 3.88e-24 | 2.29e-21 |
| 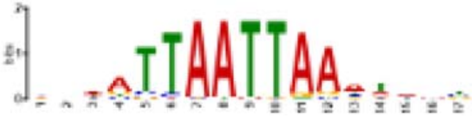 | uniprobe mouse | UP00212_1 | Lhx5_2279.1 | 4.20e-24 | 2.48e-21 |
| 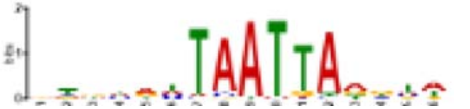 | uniprobe mouse | UP00254_1 | Pou2f1_3081.2 | 8.45e-24 | 4.99e-21 |

| | | | | | |
|---|---|---|---|---|---|
| 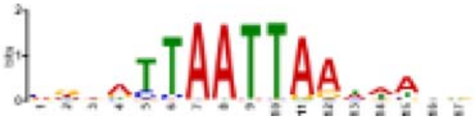 | uniprobe mouse | UP00262_1 | Lhx1_2240.2 | 1.85e-23 | 1.09e-20 |
| 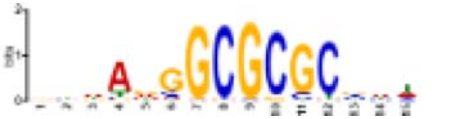 | uniprobe mouse | UP00003_1 | E2F3_primary | 3.75e-23 | 2.22e-20 |
| 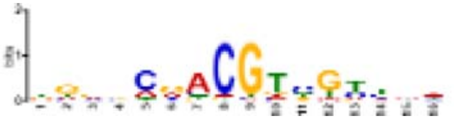 | uniprobe mouse | UP00084_2 | Gmeb1_secondary | 6.84e-23 | 4.04e-20 |
| 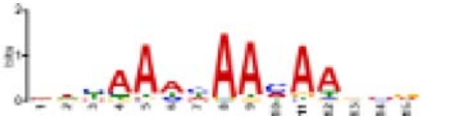 | uniprobe mouse | UP00037_1 | Zfp105_primary | 1.15e-22 | 6.77e-20 |
| 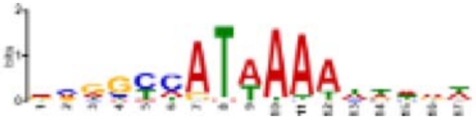 | uniprobe mouse | UP00213_1 | Hoxa9_2622.2 | 1.18e-22 | 6.95e-20 |
| 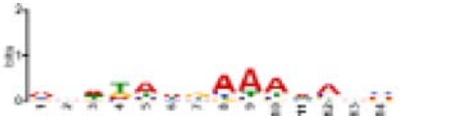 | uniprobe mouse | UP00097_2 | Mtf1_secondary | 1.18e-22 | 6.95e-20 |
| 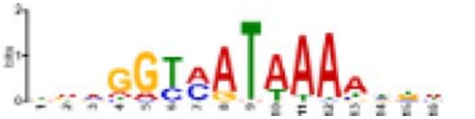 | uniprobe mouse | UP00240_1 | Cdx1_2245.1 | 1.39e-22 | 8.21e-20 |
| 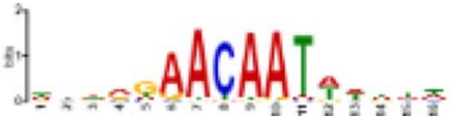 | uniprobe mouse | UP00096_1 | Sox13_primary | 1.40e-22 | 8.30e-20 |
| 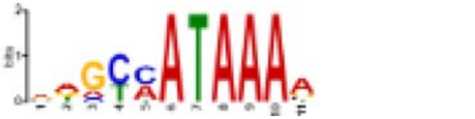 | JASPAR CORE 2014 vertebrates | MA0465.1 | CDX2 | 1.99e-22 | 1.18e-19 |
| 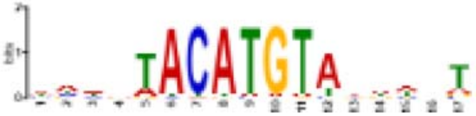 | uniprobe mouse | UP00150_1 | Irx6_2623.2 | 2.09e-22 | 1.23e-19 |
| 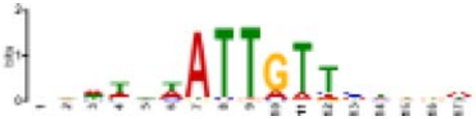 | uniprobe mouse | UP00051_1 | Sox8_primary | 2.87e-22 | 1.69e-19 |

| | | | | | |
|---|---|---|---|---|---|
| 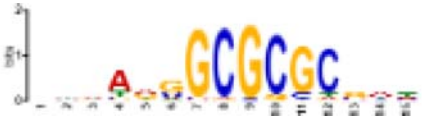 | uniprobe mouse | UP00001_1 | E2F2_primary | 3.65e-22 | 2.15e-19 |
| 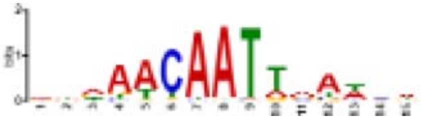 | uniprobe mouse | UP00014_1 | Sox17_primary | 1.59e-21 | 9.38e-19 |
| 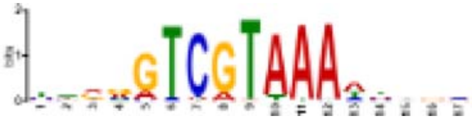 | uniprobe mouse | UP00135_1 | Hoxc12_3480.1 | 2.14e-21 | 1.26e-18 |
| 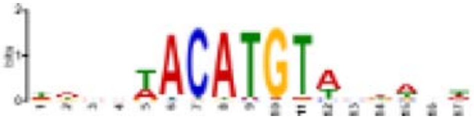 | uniprobe mouse | UP00250_1 | Irx5_2385.1 | 3.64e-21 | 2.15e-18 |
| 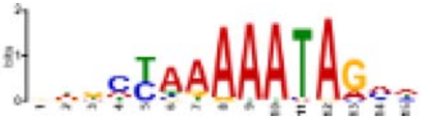 | JASPAR CORE 2014 vertebrates | MA0497.1 | MEF2C | 4.68e-21 | 2.77e-18 |
| 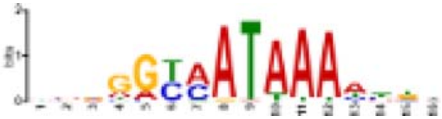 | uniprobe mouse | UP00133_1 | Cdx2_4272.1 | 5.77e-21 | 3.41e-18 |
| 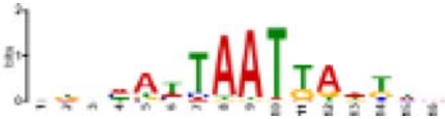 | uniprobe mouse | UP00218_1 | Dbx2_3487.1 | 6.81e-21 | 4.03e-18 |
| 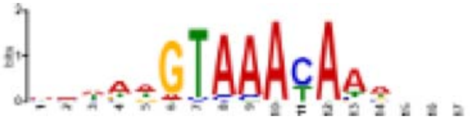 | uniprobe mouse | UP00073_1 | Foxa2_primary | 6.98e-21 | 4.12e-18 |
| 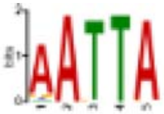 | JASPAR CORE 2014 vertebrates | MA0075.1 | Prrx2 | 8.00e-21 | 4.73e-18 |
| 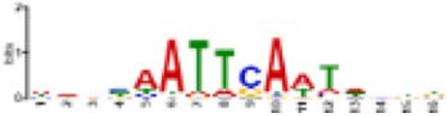 | uniprobe mouse | UP00069_1 | Sox1_primary | 1.12e-20 | 6.62e-18 |
| 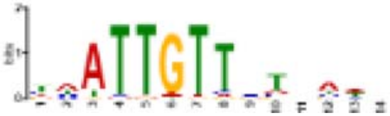 | uniprobe mouse | UP00101_1 | Sox12_primary | 1.67e-20 | 9.90e-18 |
| 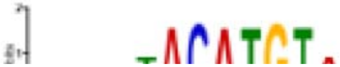 | uniprobe mouse | UP00223_2 | Irx3_2226.1 | 3.82e- | 2.26e-17 |

| | | | | | |
|---|---|---|---|---|---|
| 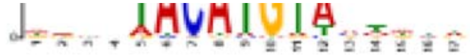 | uniprobe mouse | [illegible] | [illegible] | [illegible] 20 | [illegible] |
| 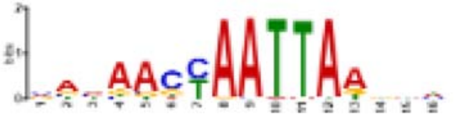 | uniprobe mouse | UP00171_1 | Msx3_3206.1 | 8.61e-20 | 5.09e-17 |
| 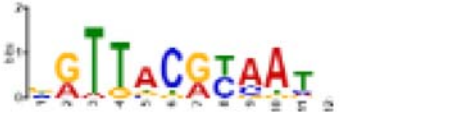 | JASPAR CORE 2014 vertebrates | MA0043.1 | HLF | 1.41e-19 | 8.32e-17 |
| 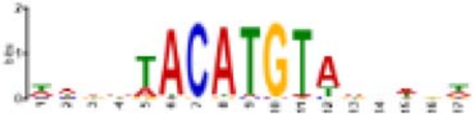 | uniprobe mouse | UP00236_1 | Irx2_0900.3 | 3.04e-19 | 1.80e-16 |
| 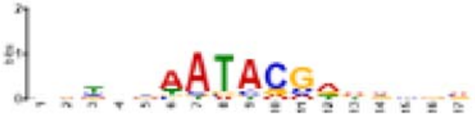 | uniprobe mouse | UP00059_2 | Arid5a_secondary | 4.57e-19 | 2.70e-16 |
| 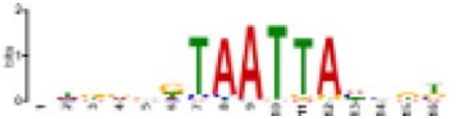 | uniprobe mouse | UP00242_1 | Hoxc8_3429.2 | 5.23e-19 | 3.09e-16 |
| 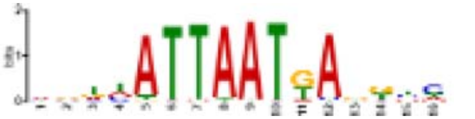 | uniprobe mouse | UP00118_1 | Pou4f3_2791.1 | 8.54e-19 | 5.04e-16 |
| 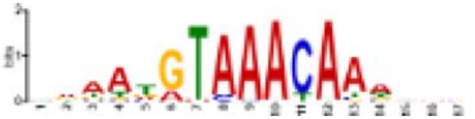 | uniprobe mouse | UP00025_1 | Foxk1_primary | 1.57e-18 | 9.26e-16 |
| 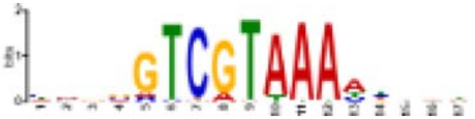 | uniprobe mouse | UP00177_1 | Hoxd12_3481.1 | 2.84e-18 | 1.68e-15 |
| 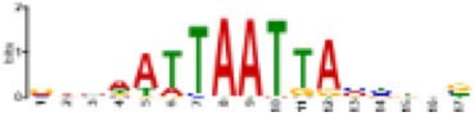 | uniprobe mouse | UP00200_1 | Nkx6-1_2825.1 | 4.69e-18 | 2.77e-15 |
| 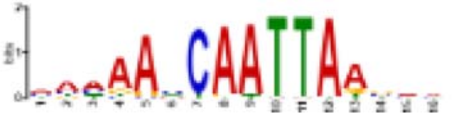 | uniprobe mouse | UP00145_1 | Barhl2_3868.1 | 5.06e-18 | 2.99e-15 |
| 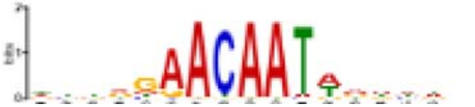 | uniprobe mouse | UP00091_1 | Sox5_primary | 5.71e-18 | 3.37e-15 |

| | | | | | |
|---|---|---|---|---|---|
| 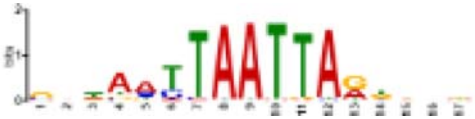 | uniprobe mouse | UP00128_1 | Pou3f2_2824.1 | 1.43e-17 | 8.47e-15 |
| 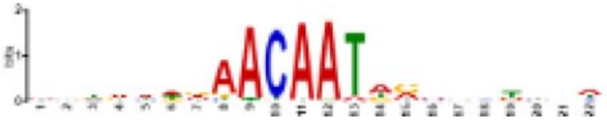 | uniprobe mouse | UP00034_1 | Sox7_primary | 1.67e-17 | 9.89e-15 |
| 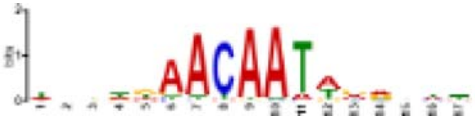 | uniprobe mouse | UP00075_1 | Sox15_primary | 4.49e-17 | 2.66e-14 |
| 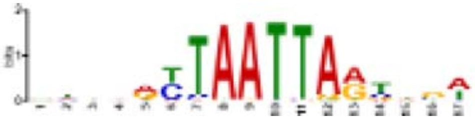 | uniprobe mouse | UP00251_1 | Esx1_3124.2 | 5.27e-17 | 3.12e-14 |
| 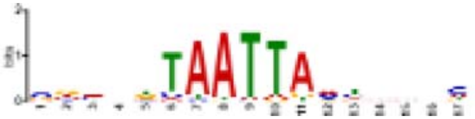 | uniprobe mouse | UP00164_1 | Hoxa7_2668.2 | 6.95e-17 | 4.11e-14 |
| 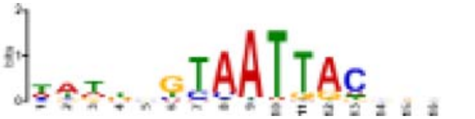 | uniprobe mouse | UP00259_1 | Hoxb6_3428.2 | 8.56e-17 | 5.06e-14 |
| 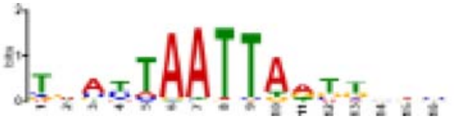 | uniprobe mouse | UP00224_1 | Pax6_3838.3 | 8.78e-17 | 5.19e-14 |
| 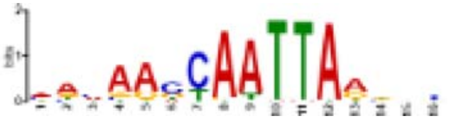 | uniprobe mouse | UP00166_1 | Barhl1_2590.2 | 1.76e-16 | 1.04e-13 |
| 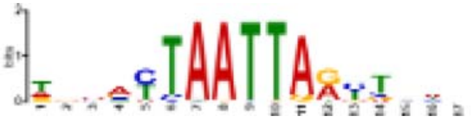 | uniprobe mouse | UP00108_1 | Alx3_3418.2 | 1.86e-16 | 1.10e-13 |
| 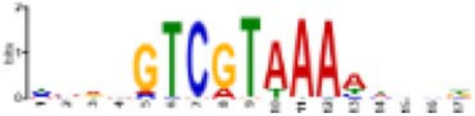 | uniprobe mouse | UP00117_1 | Hoxd11_3873.1 | 3.42e-16 | 2.02e-13 |
| 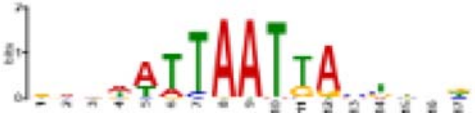 | uniprobe mouse | UP00238_1 | Nkx6-3_3446.1 | 7.32e-16 | 4.33e-13 |

| | | | | | |
|---|---|---|---|---|---|
| 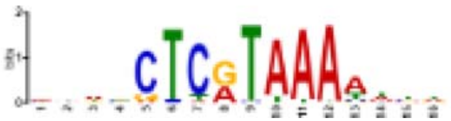 | uniprobe mouse | UP00173_1 | Hoxc13_3127.1 | 7.55e-16 | 4.46e-13 |
| 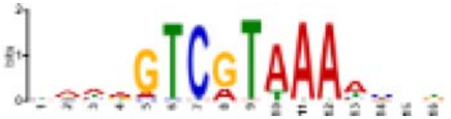 | uniprobe mouse | UP00246_1 | Hoxa11_2218.1 | 1.00e-15 | 5.91e-13 |
| 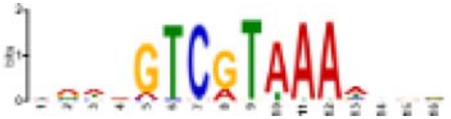 | uniprobe mouse | UP00235_1 | Hoxc11_3718.2 | 1.27e-15 | 7.48e-13 |
| 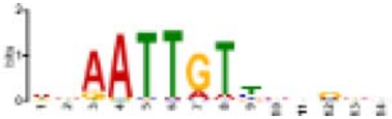 | uniprobe mouse | UP00030_2 | Sox11_secondary | 2.23e-15 | 1.32e-12 |
| 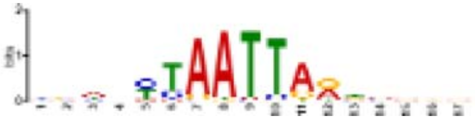 | uniprobe mouse | UP00248_1 | Pax7_3783.1 | 3.71e-15 | 2.19e-12 |
| 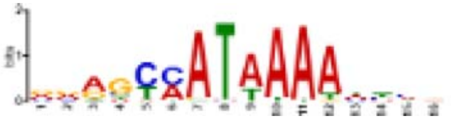 | uniprobe mouse | UP00207_1 | Hoxb9_3413.1 | 3.72e-15 | 2.20e-12 |
| 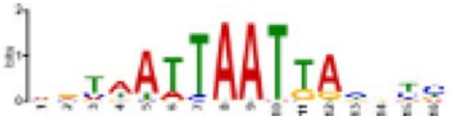 | uniprobe mouse | UP00200_2 | Nkx6-1_2825.2 | 4.87e-15 | 2.88e-12 |
| 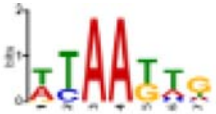 | JASPAR CORE 2014 vertebrates | MA0063.1 | Nkx2-5 | 6.74e-15 | 3.99e-12 |
| 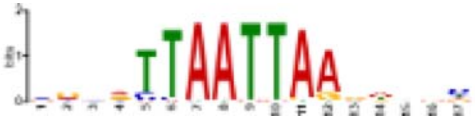 | uniprobe mouse | UP00172_1 | Prop1_3949.1 | 9.94e-15 | 5.88e-12 |
| 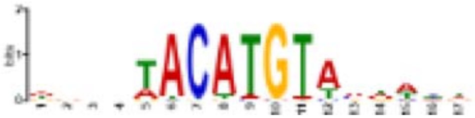 | uniprobe mouse | UP00223_1 | Irx3_0920.1 | 1.13e-14 | 6.69e-12 |
| 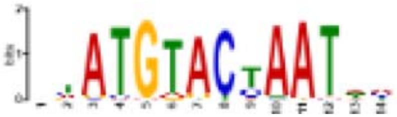 | uniprobe mouse | UP00082_1 | Zfp187_primary | 1.42e-14 | 8.40e-12 |
| 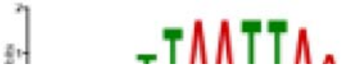 | uniprobe mouse | UP00209_2 | Cart1_1275.1 | 1.67e-[illegible] | 9.85e-12 |

| | | | | | |
|---|---|---|---|---|---|
| 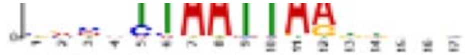 | [illegible] | [illegible] | [illegible] | [illegible] 14 | [illegible] |
| 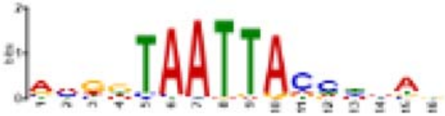 | uniprobe mouse | UP00182_1 | Hoxa6_1040.1 | 2.40e-14 | 1.42e-11 |
| 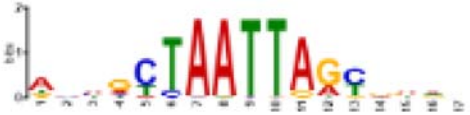 | uniprobe mouse | UP00136_1 | Prrx2_3072.1 | 2.89e-14 | 1.71e-11 |
| 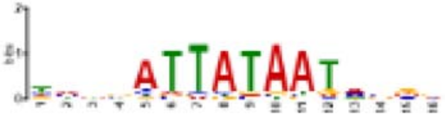 | uniprobe mouse | UP00023_2 | Sox30_secondary | 3.84e-14 | 2.27e-11 |
| 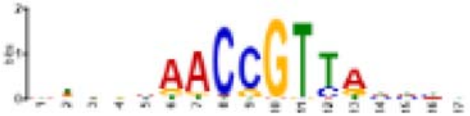 | uniprobe mouse | UP00092_1 | Myb_primary | 9.14e-14 | 5.40e-11 |
| 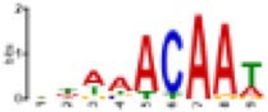 | JASPAR CORE 2014 vertebrates | MA0084.1 | SRY | 9.25e-14 | 5.47e-11 |
| 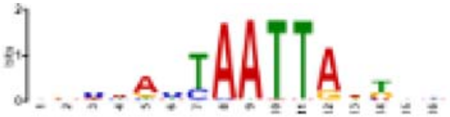 | uniprobe mouse | UP00234_1 | Msx1_3031.2 | 9.72e-14 | 5.74e-11 |
| 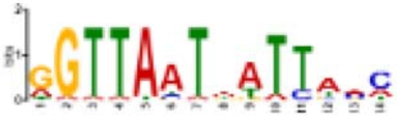 | JASPAR CORE 2014 vertebrates | MA0046.1 | HNF1A | 1.10e-13 | 6.53e-11 |
| 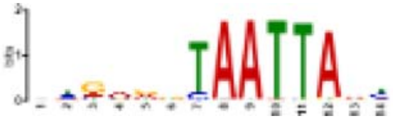 | uniprobe mouse | UP00202_1 | Dlx1_1741.2 | 2.73e-13 | 1.61e-10 |
| 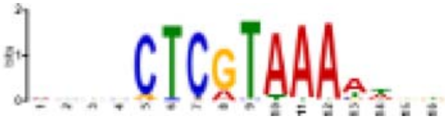 | uniprobe mouse | UP00183_1 | Hoxa13_3126.1 | 7.31e-13 | 4.32e-10 |
| 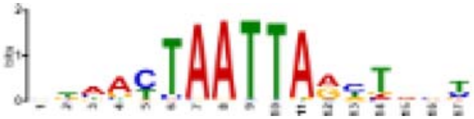 | uniprobe mouse | UP00266_1 | Prrx1_3442.1 | 8.58e-13 | 5.07e-10 |
| 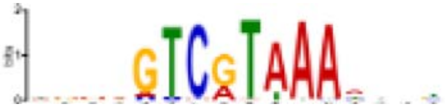 | uniprobe mouse | UP00245_1 | Hoxc10_2779.2 | 9.29e-13 | 5.49e-10 |

| | | | | | |
|---|---|---|---|---|---|
| 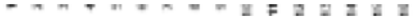 | | | | | |
| 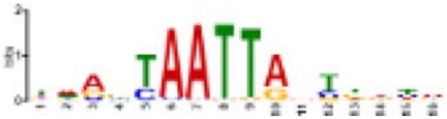 | uniprobe mouse | UP00151_1 | Barx2_3447.2 | 1.25e-12 | 7.38e-10 |
| 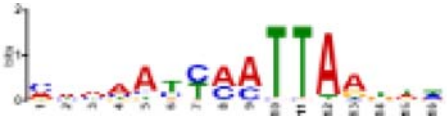 | uniprobe mouse | UP00170_1 | Isl2_3430.1 | 1.34e-12 | 7.93e-10 |
| 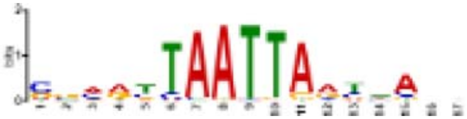 | uniprobe mouse | UP00252_1 | Hoxc5_2630.2 | 1.71e-12 | 1.01e-9 |
| 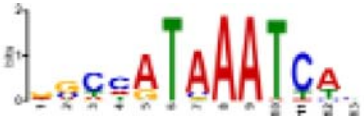 | JASPAR CORE 2014 vertebrates | MA0485.1 | Hoxc9 | 1.88e-12 | 1.11e-9 |
| 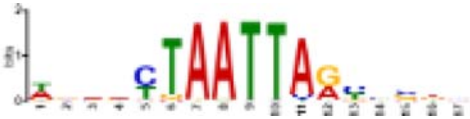 | uniprobe mouse | UP00247_1 | Pax4_3989.2 | 2.10e-12 | 1.24e-9 |
| 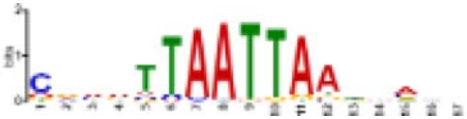 | uniprobe mouse | UP00113_1 | Hoxc4_3491.1 | 2.31e-12 | 1.36e-9 |
| 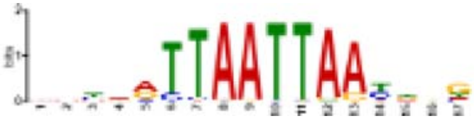 | uniprobe mouse | UP00237_1 | Otp_3496.1 | 2.86e-12 | 1.69e-9 |
| 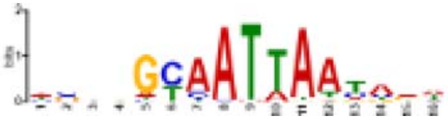 | uniprobe mouse | UP00263_1 | Hoxb8_3780.2 | 3.45e-12 | 2.04e-9 |
| 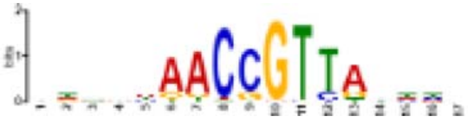 | uniprobe mouse | UP00081_1 | Mybl1_primary | 4.93e-12 | 2.92e-9 |
| 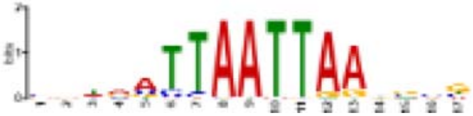 | uniprobe mouse | UP00142_1 | Uncx4.1_2281.2 | 6.91e-12 | 4.09e-9 |
| 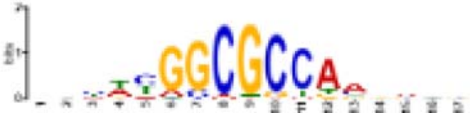 | uniprobe mouse | UP00003_2 | E2F3_secondary | 8.56e-12 | 5.06e-9 |
| 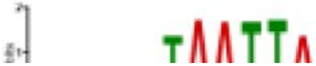 | | | | 1.09e- | |

| | | | | | |
|---|---|---|---|---|---|
| 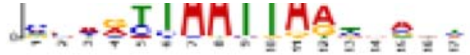 | uniprobe mouse | UP00141_1 | Vsx1_1728.1 | [illegible] 11 | 6.42e-9 |
| 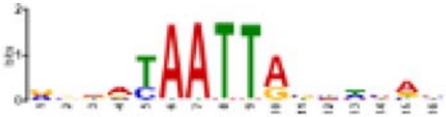 | uniprobe mouse | UP00126_1 | Dlx2_2273.2 | 1.12e-11 | 6.59e-9 |
| 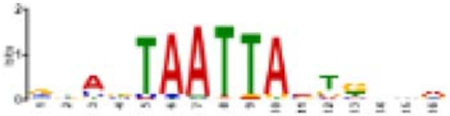 | uniprobe mouse | UP00206_1 | Hoxb7_3953.1 | 1.27e-11 | 7.52e-9 |
| 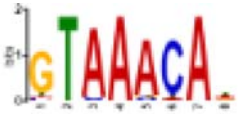 | JASPAR CORE 2014 vertebrates | MA0031.1 | FOXD1 | 1.61e-11 | 9.54e-9 |
| 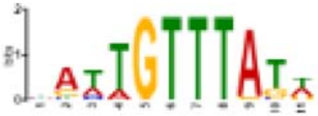 | JASPAR CORE 2014 vertebrates | MA0040.1 | Foxq1 | 2.22e-11 | 1.31e-8 |
| 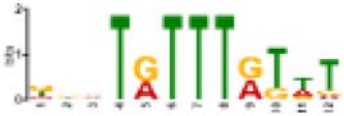 | JASPAR CORE 2014 vertebrates | MA0042.1 | FOXI1 | 2.57e-11 | 1.52e-8 |
| 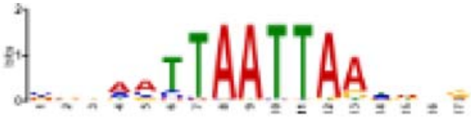 | uniprobe mouse | UP00149_1 | Phox2b_3948.1 | 4.31e-11 | 2.55e-8 |
| 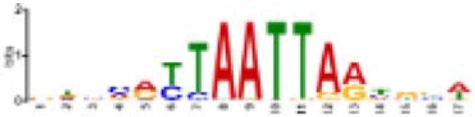 | uniprobe mouse | UP00152_1 | Arx_1738.2 | 5.48e-11 | 3.24e-8 |
| 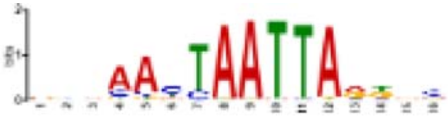 | uniprobe mouse | UP00167_1 | En1_3123.2 | 6.60e-11 | 3.90e-8 |
| 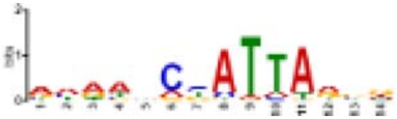 | uniprobe mouse | UP00063_2 | Hoxa3_secondary | 6.77e-11 | 4.00e-8 |
| 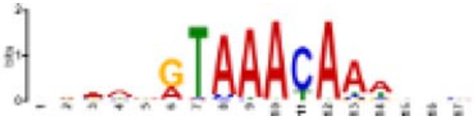 | uniprobe mouse | UP00039_1 | Foxj3_primary | 7.64e-11 | 4.51e-8 |
| 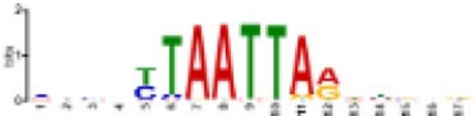 | uniprobe mouse | UP00257_1 | Shox2_2641.2 | 7.79e-11 | 4.61e-8 |

| | | | | | |
|---|---|---|---|---|---|
| 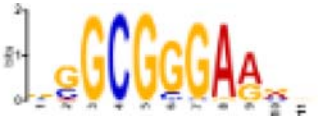 | JASPAR CORE 2014 vertebrates | MA0470.1 | E2F4 | 8.68e-11 | 5.13e-8 |
| 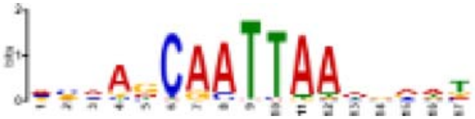 | uniprobe mouse | UP00157_1 | Hmx3_3490.2 | 1.20e-10 | 7.07e-8 |
| 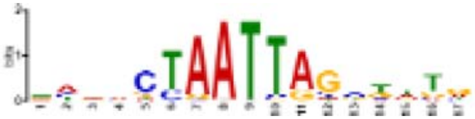 | uniprobe mouse | UP00261_1 | Lhx4_1719.2 | 1.83e-10 | 1.08e-7 |
| 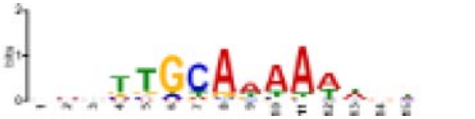 | uniprobe mouse | UP00045_2 | Mafb_secondary | 1.94e-10 | 1.15e-7 |
| 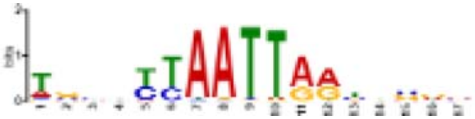 | uniprobe mouse | UP00120_1 | Lbx2_3869.2 | 2.46e-10 | 1.45e-7 |
| 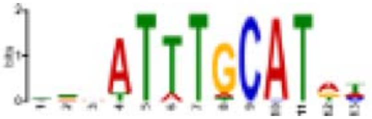 | JASPAR CORE 2014 vertebrates | MA0507.1 | POU2F2 | 2.48e-10 | 1.47e-7 |
| 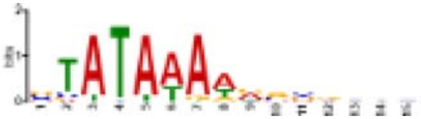 | JASPAR CORE 2014 vertebrates | MA0108.2 | TBP | 2.82e-10 | 1.67e-7 |
| 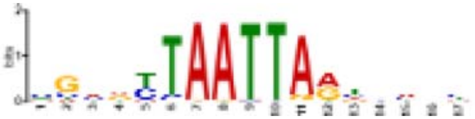 | uniprobe mouse | UP00209_1 | Cart1_0997.1 | 4.04e-10 | 2.39e-7 |
| 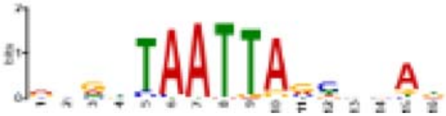 | uniprobe mouse | UP00127_1 | Gsh2_3990.2 | 4.59e-10 | 2.71e-7 |
| 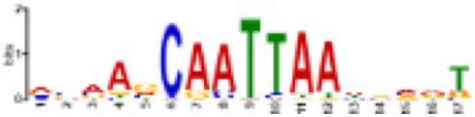 | uniprobe mouse | UP00155_1 | Hmx2_3424.3 | 5.02e-10 | 2.97e-7 |
| 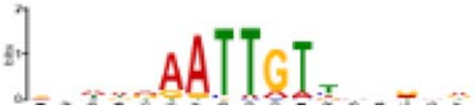 | uniprobe mouse | UP00062_2 | Sox4_secondary | 6.24e-10 | 3.69e-7 |

| | | | | | |
|---|---|---|---|---|---|
| 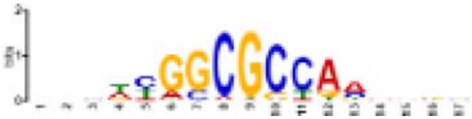 | uniprobe mouse | UP00001_2 | E2F2_secondary | 9.22e-10 | 5.45e-7 |
| 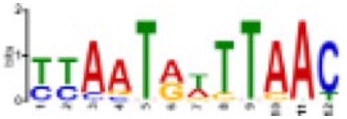 | JASPAR CORE 2014 vertebrates | MA0153.1 | HNF1B | 1.26e-9 | 7.42e-7 |
| 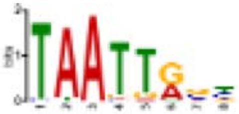 | JASPAR CORE 2014 vertebrates | MA0125.1 | Nobox | 1.28e-9 | 7.58e-7 |
| 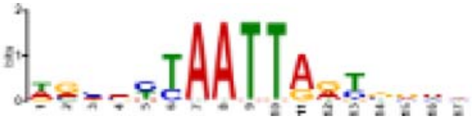 | uniprobe mouse | UP00163_1 | En2_0952.1 | 2.27e-9 | 1.34e-6 |
| 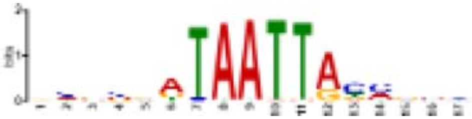 | uniprobe mouse | UP00154_1 | Dlx3_1030.1 | 2.42e-9 | 1.43e-6 |
| 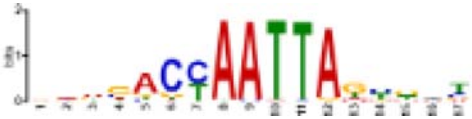 | uniprobe mouse | UP00156_1 | Msx2_3449.1 | 2.55e-9 | 1.51e-6 |
| 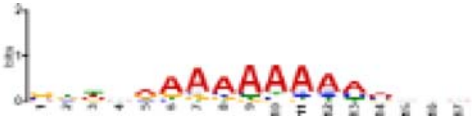 | uniprobe mouse | UP00077_2 | Srf_secondary | 2.70e-9 | 1.60e-6 |
| 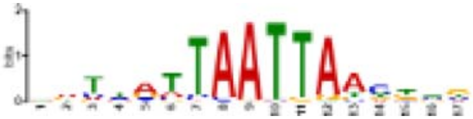 | uniprobe mouse | UP00196_1 | Hoxa4_3426.1 | 3.11e-9 | 1.84e-6 |
| 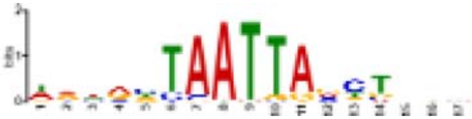 | uniprobe mouse | UP00140_1 | Hoxd1_3448.1 | 4.08e-9 | 2.41e-6 |
| 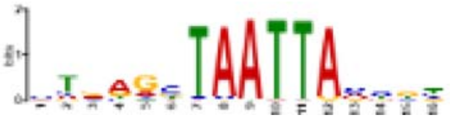 | uniprobe mouse | UP00264_1 | Hoxa1_3425.1 | 6.06e-9 | 3.58e-6 |
| 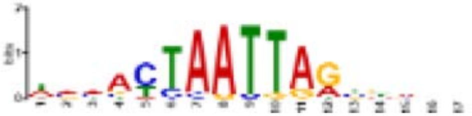 | uniprobe mouse | UP00115_1 | Lhx2_0953.2 | 7.03e-9 | 4.16e-6 |
| 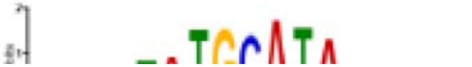 | uniprobe mouse | UP00211_1 | Pou3f3_3235.2 | 8.40e- | 4.96e-6 |

| | | | | | |
|---|---|---|---|---|---|
| 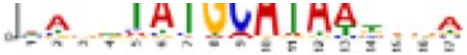 | [illegible] | [illegible] | [illegible] | [illegible] 9 | [illegible] |
| 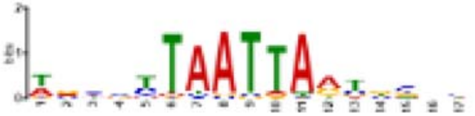 | uniprobe mouse | UP00116_1 | Rhox6_4251.1 | 1.07e-8 | 6.29e-6 |
| 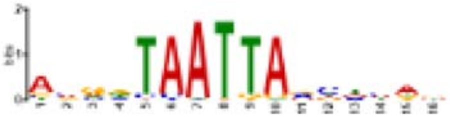 | uniprobe mouse | UP00189_1 | Hoxa5_3415.1 | 1.28e-8 | 7.54e-6 |
| 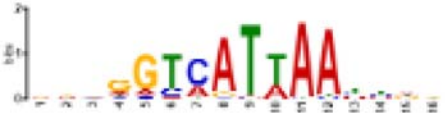 | uniprobe mouse | UP00197_1 | Hoxc9_2367.2 | 1.30e-8 | 7.71e-6 |
| 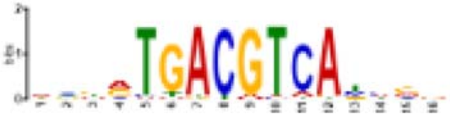 | uniprobe mouse | UP00020_1 | Atf1_primary | 1.34e-8 | 7.95e-6 |
| 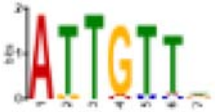 | JASPAR CORE 2014 vertebrates | MA0087.1 | Sox5 | 1.38e-8 | 8.17e-6 |
| 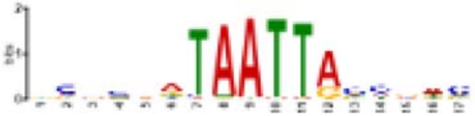 | uniprobe mouse | UP00110_1 | Dlx4_3488.2 | 1.90e-8 | 1.12e-5 |
| 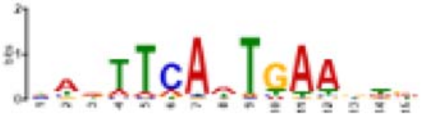 | uniprobe mouse | UP00012_1 | Bbx_primary | 2.16e-8 | 1.28e-5 |
| 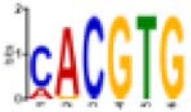 | JASPAR CORE 2014 vertebrates | MA0004.1 | Arnt | 2.45e-8 | 1.45e-5 |
| 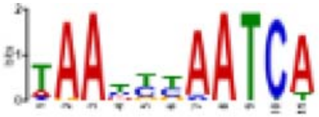 | JASPAR CORE 2014 vertebrates | MA0468.1 | DUX4 | 2.56e-8 | 1.51e-5 |
| 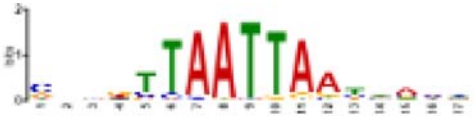 | uniprobe mouse | UP00144_1 | Hoxb4_2627.1 | 3.29e-8 | 1.94e-5 |
| 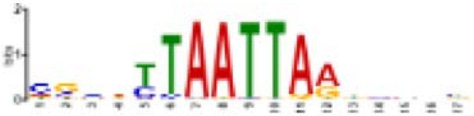 | uniprobe mouse | UP00187_1 | Alx4_1744.1 | 3.57e-8 | 2.11e-5 |

| | | | | | |
|---|---|---|---|---|---|
| 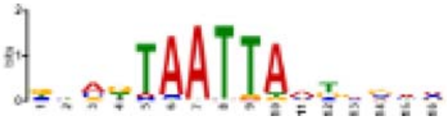 | uniprobe mouse | UP00164_2 | Hoxa7_3750.1 | 4.28e-8 | 2.53e-5 |
| 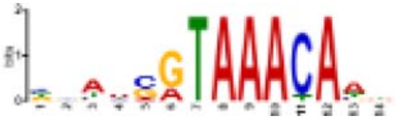 | JASPAR CORE 2014 vertebrates | MA0030.1 | FOXF2 | 4.74e-8 | 2.80e-5 |
| 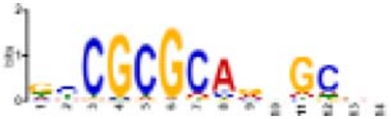 | uniprobe mouse | UP00065_2 | Zfp161_secondary | 5.00e-8 | 2.95e-5 |
| 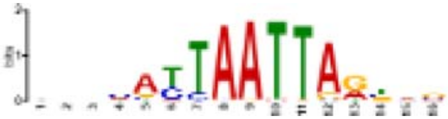 | uniprobe mouse | UP00221_1 | Phox2a_3947.1 | 6.86e-8 | 4.06e-5 |
| 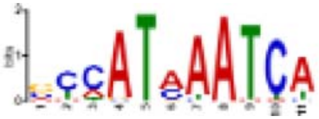 | JASPAR CORE 2014 vertebrates | MA0594.1 | Hoxa9 | 6.96e-8 | 4.11e-5 |
| 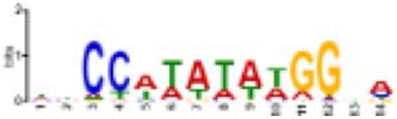 | uniprobe mouse | UP00077_1 | Srf_primary | 7.01e-8 | 4.14e-5 |
| 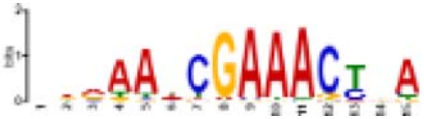 | uniprobe mouse | UP00074_1 | Isgf3g_primary | 7.71e-8 | 4.55e-5 |
| 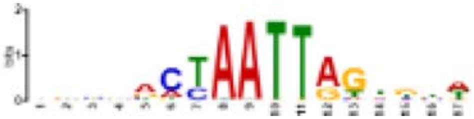 | uniprobe mouse | UP00204_1 | Gbx1_2883.2 | 8.50e-8 | 5.02e-5 |
| 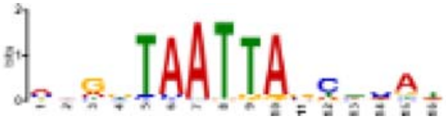 | uniprobe mouse | UP00214_1 | Hoxb5_3122.2 | 8.86e-8 | 5.23e-5 |
| 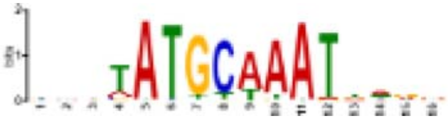 | uniprobe mouse | UP00191_1 | Pou2f2_3748.1 | 1.27e-7 | 7.51e-5 |
| 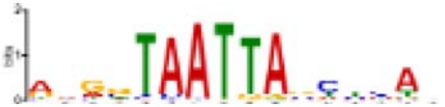 | uniprobe mouse | UP00174_1 | Hoxa2_3079.1 | 2.96e-7 | 1.75e-4 |

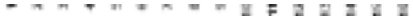

| | | | | | |
|---|---|---|---|---|---|
| 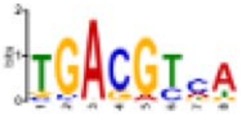 | JASPAR CORE 2014 vertebrates | MA0018.2 | CREB1 | 2.99e-7 | 1.77e-4 |
| 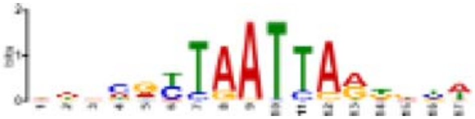 | uniprobe mouse | UP00256_1 | Lhx6_2272.1 | 3.39e-7 | 2.00e-4 |
| 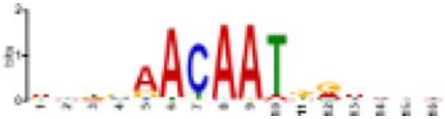 | uniprobe mouse | UP00023_1 | Sox30_primary | 5.71e-7 | 3.37e-4 |
| 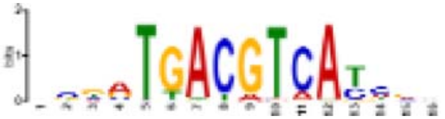 | uniprobe mouse | UP00103_1 | Jundm2_primary | 6.53e-7 | 3.86e-4 |
| 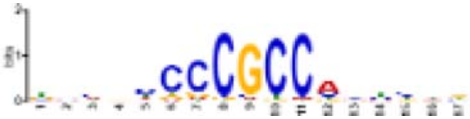 | uniprobe mouse | UP00000_2 | Smad3_secondary | 8.02e-7 | 4.74e-4 |
| 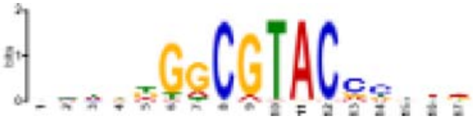 | uniprobe mouse | UP00094_1 | Zfp128_primary | 8.57e-7 | 5.06e-4 |
| 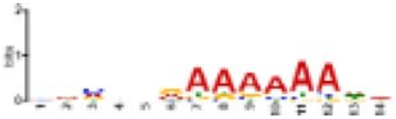 | uniprobe mouse | UP00028_2 | Tcfap2e_secondary | 8.97e-7 | 5.30e-4 |
| 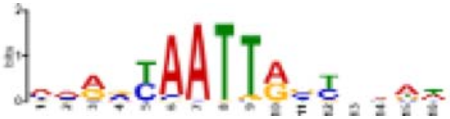 | uniprobe mouse | UP00181_1 | Barx1_2877.1 | 9.60e-7 | 5.67e-4 |
| 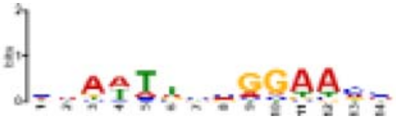 | uniprobe mouse | UP00085_2 | Sfpi1_secondary | 1.07e-6 | 6.34e-4 |
| 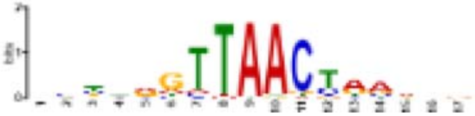 | uniprobe mouse | UP00089_1 | Tcf1_primary | 1.43e-6 | 8.44e-4 |
| 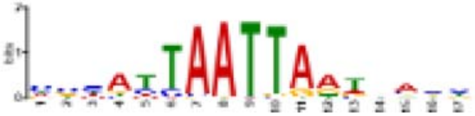 | uniprobe mouse | UP00175_1 | Lhx9_3492.1 | 1.98e-6 | 1.17e-3 |
| 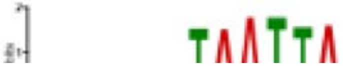 | uniprobe mouse | UP00063_1 | Hoxa3_primary | 2.01e-[illegible] | 1.19e-3 |

| | | | | | |
|---|---|---|---|---|---|
| 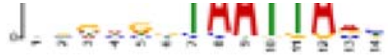 | uniprobe mouse | [illegible] | Hoxd3_primary | [illegible]6 | [illegible] |
| 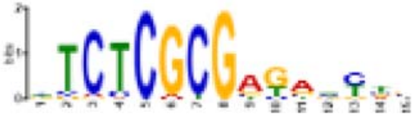 | JASPAR CORE 2014 vertebrates | MA0527.1 | ZBTB33 | 2.64e-6 | 1.56e-3 |
| 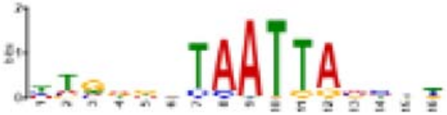 | uniprobe mouse | UP00241_1 | Hoxd3_1742.2 | 3.08e-6 | 1.82e-3 |
| 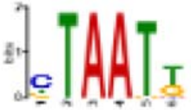 | JASPAR CORE 2014 vertebrates | MA0132.1 | Pdx1 | 3.41e-6 | 2.01e-3 |
| 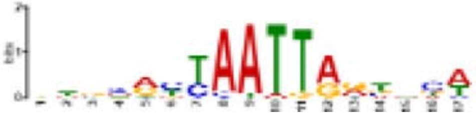 | uniprobe mouse | UP00139_1 | Nkx1-2_3214.1 | 3.90e-6 | 2.30e-3 |
| 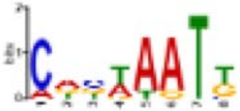 | JASPAR CORE 2014 vertebrates | MA0158.1 | HOXA5 | 4.54e-6 | 2.68e-3 |
| 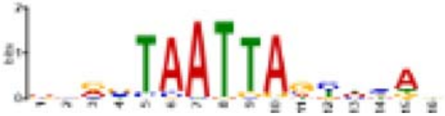 | uniprobe mouse | UP00124_1 | Ipf1_3815.1 | 4.67e-6 | 2.76e-3 |
| 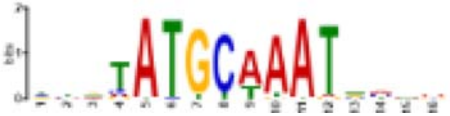 | uniprobe mouse | UP00179_1 | Pou2f3_3986.2 | 5.08e-6 | 3.00e-3 |
| 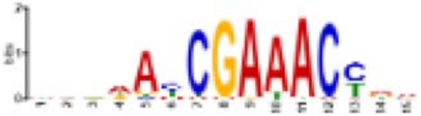 | uniprobe mouse | UP00040_1 | Irf5_primary | 6.31e-6 | 3.72e-3 |
| 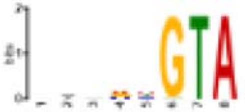 | JASPAR CORE 2014 vertebrates | MA0032.1 | FOXC1 | 8.08e-6 | 4.76e-3 |
| 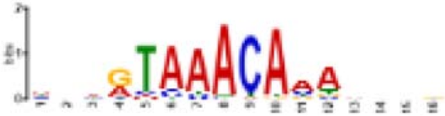 | uniprobe mouse | UP00041_1 | Foxj1_primary | 1.02e-5 | 6.03e-3 |
| 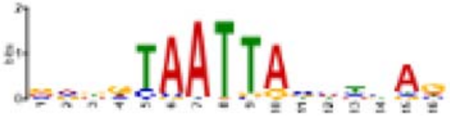 | uniprobe mouse | UP00233_1 | Meox1_2310.2 | 1.07e-5 | 6.29e-3 |

| | | | | | |
|---|---|---|---|---|---|
| 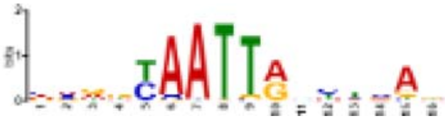 | uniprobe mouse | UP00138_1 | Bsx_3483.2 | 1.17e-5 | 6.87e-3 |
| 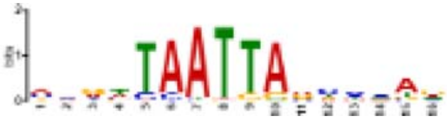 | uniprobe mouse | UP00215_1 | Vax1_3499.1 | 1.70e-5 | 9.99e-3 |
| 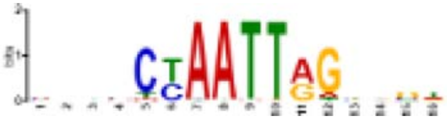 | uniprobe mouse | UP00243_1 | Isx_3445.1 | 1.75e-5 | 1.03e-2 |
| 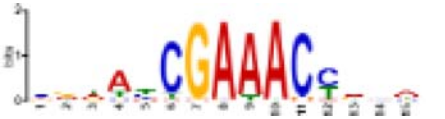 | uniprobe mouse | UP00018_1 | Irf4_primary | 2.05e-5 | 1.21e-2 |
| 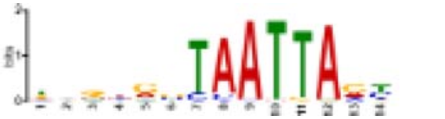 | uniprobe mouse | UP00391_1 | Hoxa3_2783.2 | 2.07e-5 | 1.22e-2 |
| 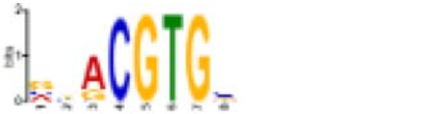 | JASPAR CORE 2014 vertebrates | MA0259.1 | HIF1A::ARNT | 2.32e-5 | 1.36e-2 |
| 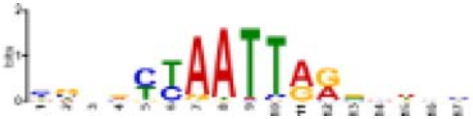 | uniprobe mouse | UP00178_1 | Og2x_3719.1 | 2.49e-5 | 1.46e-2 |
| 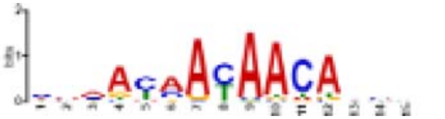 | uniprobe mouse | UP00025_2 | Foxk1_secondary | 3.16e-5 | 1.85e-2 |
| 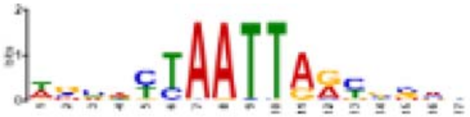 | uniprobe mouse | UP00253_1 | Rax_3443.1 | 4.11e-5 | 2.40e-2 |
| 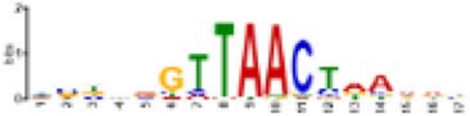 | uniprobe mouse | UP00390_1 | Tcf1_2666.2 | 5.14e-5 | 2.99e-2 |
| 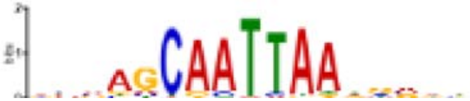 | uniprobe mouse | UP00104_1 | Hmx1_3423.1 | 5.57e-5 | 3.24e-2 |

| | | | | | |
|---|---|---|---|---|---|
| 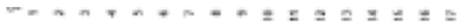 | | | | | |
| 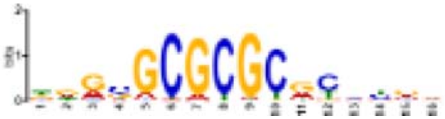 | uniprobe mouse | UP00065_1 | Zfp161_primary | 5.72e-5 | 3.32e-2 |
| 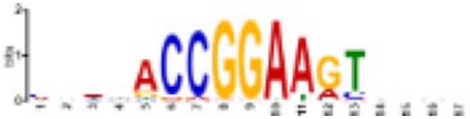 | uniprobe mouse | UP00013_1 | Gabpa_primary | 7.50e-5 | 4.33e-2 |

#### Input Files

##### Sequences

| Primary Sequences | Number | Control Sequences | Number | | | | |
|---|---|---|---|---|---|---|---|
| Control Sequences | Number | | | | | | |
| ud300expr.fa | 6525 | ud300nonexpr.fa | 15480 | ud300expr.fa | 6525 | ud300nonexpr.fa | 15480 |

##### Motifs

| Database | Source | Motif Count |
|---|---|---|
| JASPAR CORE 2014 vertebrates | db/JASPAR/JASPAR_CORE_2014_vertebrates.meme | 205 |
| uniprobe mouse | db/MOUSE/uniprobe_mouse.meme | 386 |
| JASPAR CORE 2014 vertebrates | db/JASPAR/JASPAR_CORE_2014_vertebrates.meme | 205 |
| uniprobe mouse | db/MOUSE/uniprobe_mouse.meme | 386 |

**AME version**
4.11.24.11.2 (Release date: Thu May 05 14:58:55 2016 -0700Thu May 05 14:58:55 2016 -0700)
Copyright © Robert McLeay & Timothy Bailey, 2009.

**Command line summary**

```
/home/meme/meme_4.11.2/bin/ame --verbose 1 --oc . --control ud300nonexpr.fa --bgformat 2 --bgfile background.model --scoring avg --method ranksum --pvalue-report-threshold 0.05 ud300expr.fa db/JASPAR/JASPAR_CORE_2014_vertebrates.meme db/MOUSE/uniprobe_mouse.meme
```

### AME
Analysis of Motif Enrichment

For further information on how to interpret these results or to get a copy of the MEME software please access http://meme-suite.org.

If you use AME in your research, please cite the following paper:
Robert McLeay and Timothy L. Bailey, "Motif Enrichment Analysis: A unified framework and method evaluation", *BMC Bioinformatics*, **11**:165, 2010, doi:10.1186/1471-2105-11-165. [full text]

Enriched Motifs | Input Files | Program information

#### Enriched Motifs

Fixed partition size: number of primary sequences (6525)

Sequence motif score: avg_odds
Background model source: file background.model
Background model frequencies: 0.291,0.209,0.209,0.291
Total pseudocount added to a motif column: 0.25

Statistical test: Wilcoxon rank-sum test
Ranksum method: quick
Threshold *p*-value for reporting results: 0.05
Number of multiple tests for Bonferroni correction: #Motifs × #PartitionsTested = 591 × 1 = 591Fixed partition size: number of primary sequences (6525)

Sequence motif score: avg_odds
Background model source: file background.model
Background model frequencies: 0.291,0.209,0.209,0.291
Total pseudocount added to a motif column: 0.25

Statistical test: Wilcoxon rank-sum test
Ranksum method: quick
Threshold *p*-value for reporting results: 0.05
Number of multiple tests for Bonferroni correction: #Motifs × #PartitionsTested = 591 × 1 = 591

| Logo | Database | ID | Name | *p*-value | Adjusted *p*-value |
|---|---|---|---|---|---|
| 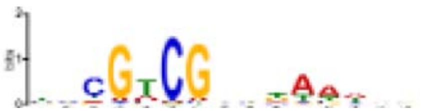 | uniprobe mouse | UP00049_2 | Sp100_secondary | 2.35e-131 | 1.39e-128 |

| | | | | | |
|---|---|---|---|---|---|
| 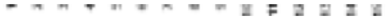 | | | | | |
| 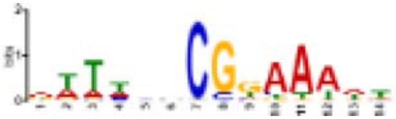 | uniprobe mouse | UP00049_1 | Sp100_primary | 2.74e-90 | 1.62e-87 |
| 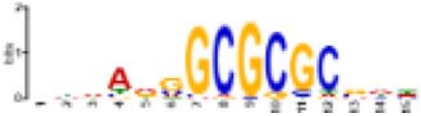 | uniprobe mouse | UP00001_1 | E2F2_primary | 1.32e-85 | 7.78e-83 |
| 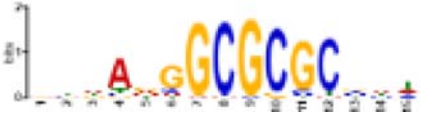 | uniprobe mouse | UP00003_1 | E2F3_primary | 8.07e-82 | 4.77e-79 |
| 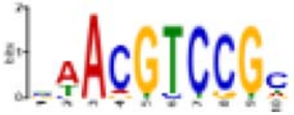 | JASPAR CORE 2014 vertebrates | MA0131.1 | HINFP | 1.16e-79 | 6.87e-77 |
| 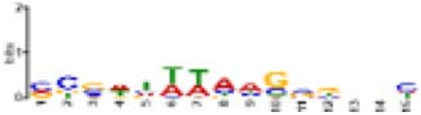 | uniprobe mouse | UP00029_2 | Tbp_secondary | 3.44e-71 | 2.03e-68 |
| 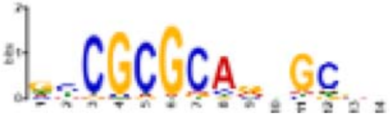 | uniprobe mouse | UP00065_2 | Zfp161_secondary | 1.07e-67 | 6.34e-65 |
| 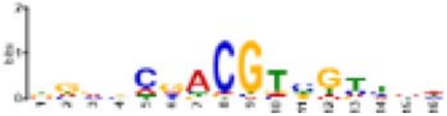 | uniprobe mouse | UP00084_2 | Gmeb1_secondary | 5.24e-63 | 3.09e-60 |
| 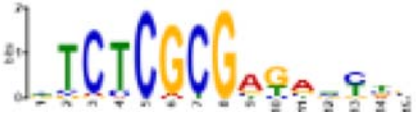 | JASPAR CORE 2014 vertebrates | MA0527.1 | ZBTB33 | 5.92e-62 | 3.50e-59 |
| 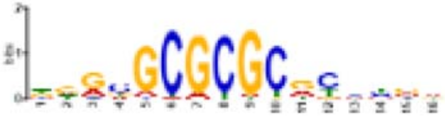 | uniprobe mouse | UP00065_1 | Zfp161_primary | 3.16e-61 | 1.87e-58 |
| 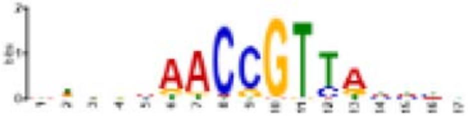 | uniprobe mouse | UP00092_1 | Myb_primary | 4.57e-60 | 2.70e-57 |
| 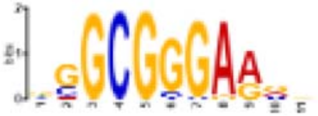 | JASPAR CORE 2014 vertebrates | MA0470.1 | E2F4 | 4.63e-60 | 2.73e-57 |

| | | | | | |
|---|---|---|---|---|---|
| 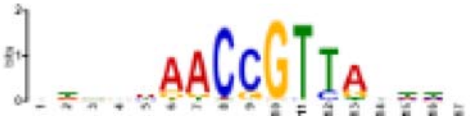 | uniprobe mouse | UP00081_1 | Mybl1_primary | 2.22e-58 | 1.31e-55 |
| 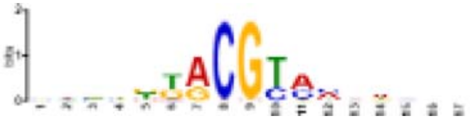 | uniprobe mouse | UP00084_1 | Gmeb1_primary | 4.47e-58 | 2.64e-55 |
| 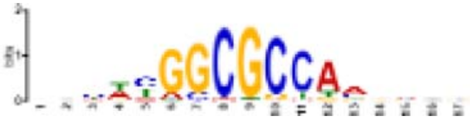 | uniprobe mouse | UP00003_2 | E2F3_secondary | 1.71e-52 | 1.01e-49 |
| 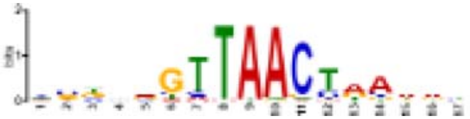 | uniprobe mouse | UP00390_1 | Tcf1_2666.2 | 1.13e-48 | 6.68e-46 |
| 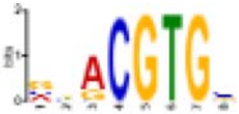 | JASPAR CORE 2014 vertebrates | MA0259.1 | HIF1A::ARNT | 1.60e-48 | 9.47e-46 |
| 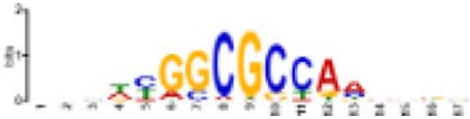 | uniprobe mouse | UP00001_2 | E2F2_secondary | 5.83e-48 | 3.45e-45 |
| 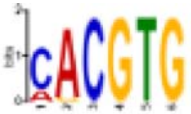 | JASPAR CORE 2014 vertebrates | MA0004.1 | Arnt | 1.86e-44 | 1.10e-41 |
| 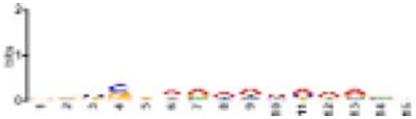 | uniprobe mouse | UP00058_2 | Tcf3_secondary | 1.03e-42 | 6.10e-40 |
| 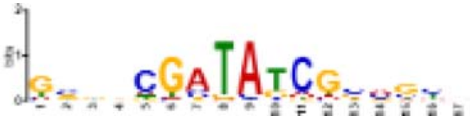 | uniprobe mouse | UP00100_2 | Gata6_secondary | 1.67e-41 | 9.84e-39 |
| 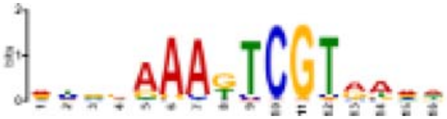 | uniprobe mouse | UP00072_2 | IRC900814_secondary | 5.34e-41 | 3.16e-38 |
| | | | | [illegible] | |

| | | | | | |
|---|---|---|---|---|---|
| 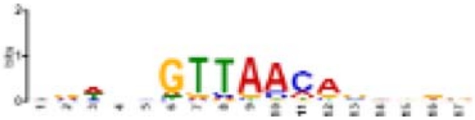 | uniprobe mouse | UP00012_2 | Bbx_secondary | 8.47e-41 | 5.00e-38 |
| 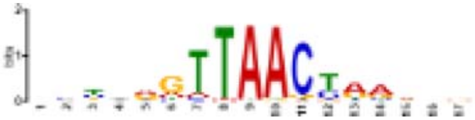 | uniprobe mouse | UP00089_1 | Tcf1_primary | 1.53e-40 | 9.07e-38 |
| 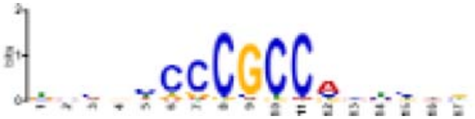 | uniprobe mouse | UP00000_2 | Smad3_secondary | 9.46e-40 | 5.59e-37 |
| 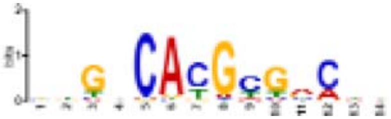 | uniprobe mouse | UP00060_2 | Max_secondary | 2.08e-35 | 1.23e-32 |
| 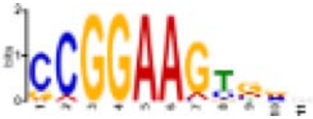 | JASPAR CORE 2014 vertebrates | MA0062.2 | GABPA | 4.60e-35 | 2.72e-32 |
| 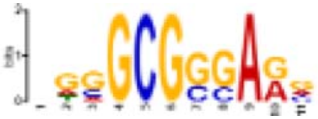 | JASPAR CORE 2014 vertebrates | MA0024.2 | E2F1 | 7.31e-35 | 4.32e-32 |
| 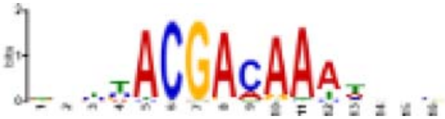 | uniprobe mouse | UP00072_1 | IRC900814_primary | 8.04e-34 | 4.75e-31 |
| 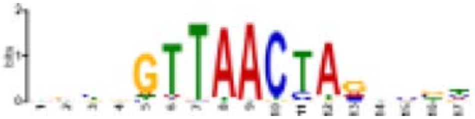 | uniprobe mouse | UP00222_1 | Tcf2_0913.2 | 1.25e-33 | 7.39e-31 |
| 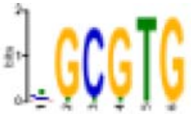 | JASPAR CORE 2014 vertebrates | MA0006.1 | Arnt::Ahr | 2.84e-31 | 1.68e-28 |
| 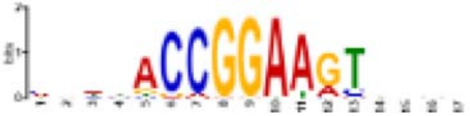 | uniprobe mouse | UP00013_1 | Gabpa_primary | 3.62e-31 | 2.14e-28 |
| 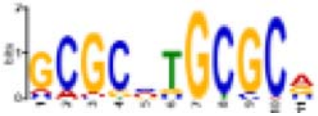 | JASPAR CORE 2014 vertebrates | MA0506.1 | NRF1 | 5.24e-31 | 3.10e-28 |
| 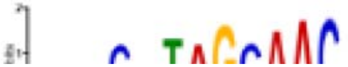 | uniprobe mouse | UP00076_1 | Rfxdc2_primary | 6.03e- | 3.56e-28 |

| | | | | | |
|---|---|---|---|---|---|
| 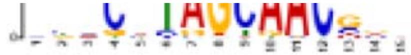 | [illegible] | [illegible] | [illegible] | [illegible] 31 | [illegible] |
| 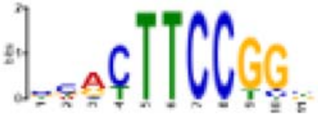 | JASPAR CORE 2014 vertebrates | MA0076.2 | ELK4 | 5.96e-28 | 3.52e-25 |
| 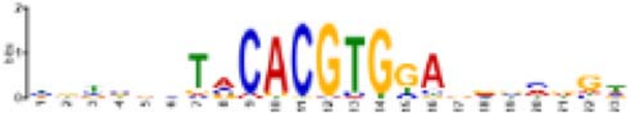 | uniprobe mouse | UP00050_2 | Bhlhb2_secondary | 3.61e-27 | 2.13e-24 |
| 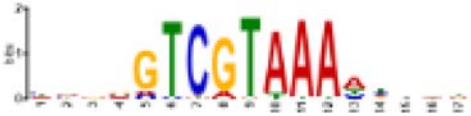 | uniprobe mouse | UP00177_1 | Hoxd12_3481.1 | 1.62e-26 | 9.60e-24 |
| 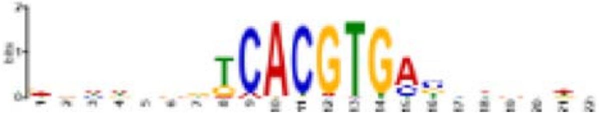 | uniprobe mouse | UP00050_1 | Bhlhb2_primary | 1.67e-26 | 9.86e-24 |
| 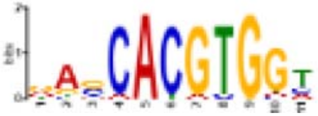 | JASPAR CORE 2014 vertebrates | MA0059.1 | MYC::MAX | 4.00e-26 | 2.36e-23 |
| 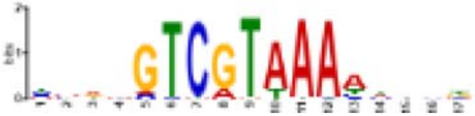 | uniprobe mouse | UP00117_1 | Hoxd11_3873.1 | 2.43e-25 | 1.43e-22 |
| 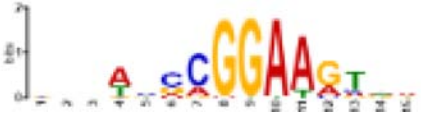 | uniprobe mouse | UP00015_1 | Ehf_primary | 6.13e-25 | 3.63e-22 |
| 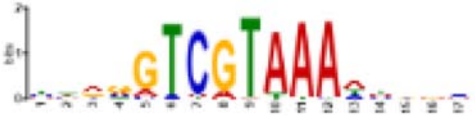 | uniprobe mouse | UP00135_1 | Hoxc12_3480.1 | 7.71e-24 | 4.56e-21 |
| 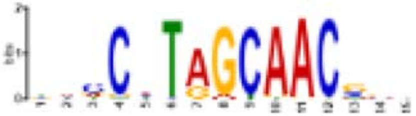 | uniprobe mouse | UP00056_1 | Rfx4_primary | 1.02e-23 | 6.03e-21 |
| 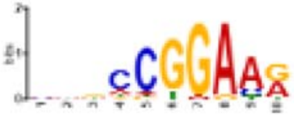 | JASPAR CORE 2014 vertebrates | MA0028.1 | ELK1 | 1.12e-20 | 6.65e-18 |
| 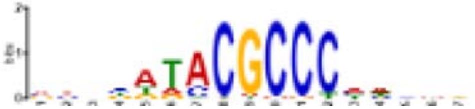 | uniprobe mouse | UP00093_2 | Klf7_secondary | 1.33e-20 | 7.88e-18 |

| | | | | | |
|---|---|---|---|---|---|
| 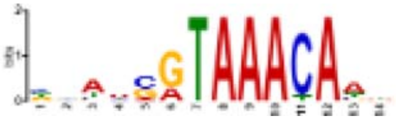 | JASPAR CORE 2014 vertebrates | MA0030.1 | FOXF2 | 9.67e-20 | 5.71e-17 |
| 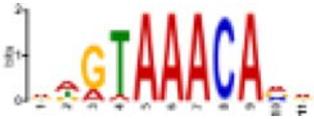 | JASPAR CORE 2014 vertebrates | MA0593.1 | FOXP2 | 8.79e-18 | 5.20e-15 |
| 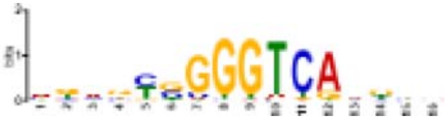 | uniprobe mouse | UP00048_2 | Rara_secondary | 2.47e-17 | 1.46e-14 |
| 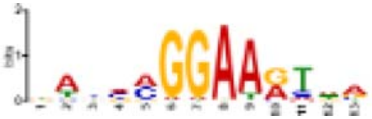 | uniprobe mouse | UP00090_1 | Elf3_primary | 2.60e-17 | 1.54e-14 |
| 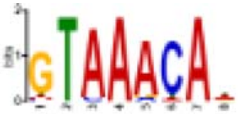 | JASPAR CORE 2014 vertebrates | MA0031.1 | FOXD1 | 2.60e-17 | 1.54e-14 |
| 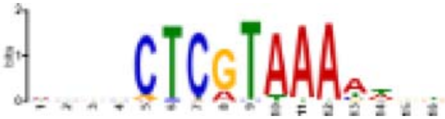 | uniprobe mouse | UP00183_1 | Hoxa13_3126.1 | 8.07e-17 | 4.77e-14 |
| 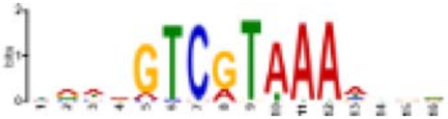 | uniprobe mouse | UP00235_1 | Hoxc11_3718.2 | 1.22e-16 | 7.20e-14 |
| 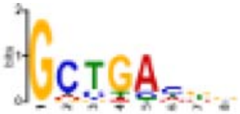 | JASPAR CORE 2014 vertebrates | MA0117.1 | Mafb | 1.22e-16 | 7.20e-14 |
| 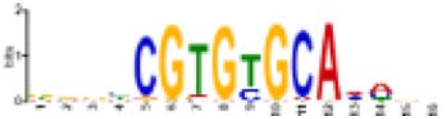 | uniprobe mouse | UP00097_1 | Mtf1_primary | 2.98e-16 | 1.76e-13 |
| 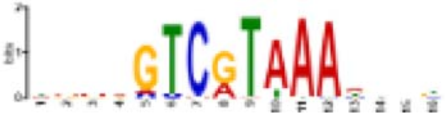 | uniprobe mouse | UP00245_1 | Hoxc10_2779.2 | 4.10e-16 | 2.42e-13 |
| 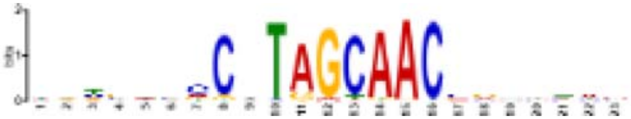 | uniprobe mouse | UP00098_1 | Rfx3_primary | 6.70e-16 | 3.96e-13 |

| | | | | | |
|---|---|---|---|---|---|
| 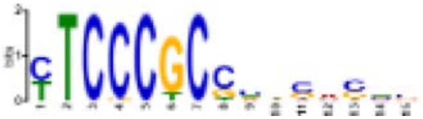 | JASPAR CORE 2014 vertebrates | MA0469.1 | E2F3 | 7.53e-16 | 4.45e-13 |
| 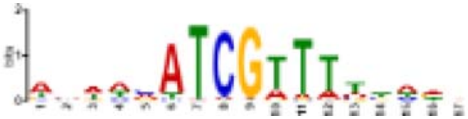 | uniprobe mouse | UP00114_1 | Homez_1063.2 | 9.10e-16 | 5.38e-13 |
| 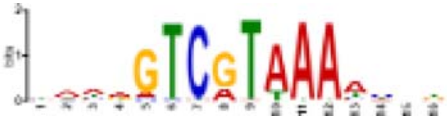 | uniprobe mouse | UP00246_1 | Hoxa11_2218.1 | 1.68e-15 | 9.91e-13 |
| 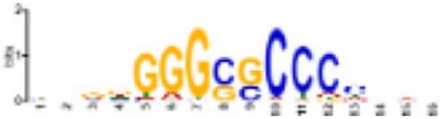 | uniprobe mouse | UP00088_1 | Plagl1_primary | 2.39e-15 | 1.41e-12 |
| 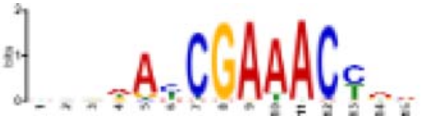 | uniprobe mouse | UP00040_1 | Irf5_primary | 3.04e-15 | 1.80e-12 |
| 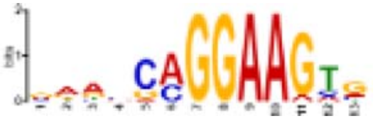 | JASPAR CORE 2014 vertebrates | MA0473.1 | ELF1 | 5.59e-15 | 3.30e-12 |
| 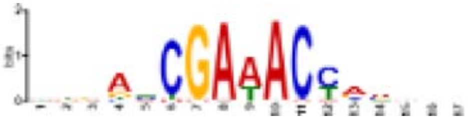 | uniprobe mouse | UP00011_1 | Irf6_primary | 7.73e-15 | 4.57e-12 |
| 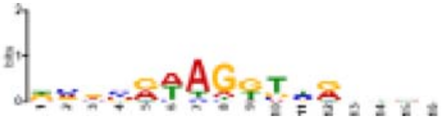 | uniprobe mouse | UP00053_2 | Rxra_secondary | 9.93e-15 | 5.87e-12 |
| 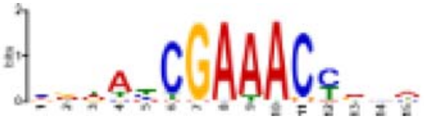 | uniprobe mouse | UP00018_1 | Irf4_primary | 1.42e-14 | 8.38e-12 |
| 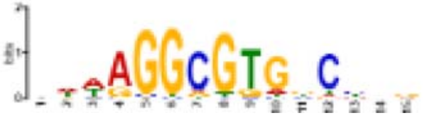 | uniprobe mouse | UP00002_2 | Sp4_secondary | 1.73e-14 | 1.02e-11 |
| 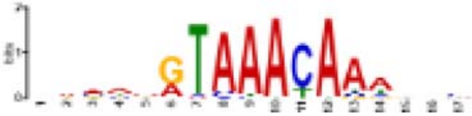 | uniprobe mouse | UP00039_1 | Foxj3_primary | 1.99e-14 | 1.18e-11 |
| 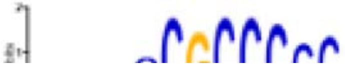 | uniprobe mouse | UP00002_1 | Sp4_primary | 2.38e- | 1.40e-11 |

| | | | | | |
|---|---|---|---|---|---|
| 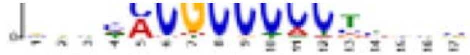 | uniprobe mouse | [illegible] | Sp1_primary | [illegible] 14 | [illegible] |
| 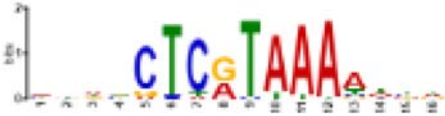 | uniprobe mouse | UP00173_1 | Hoxc13_3127.1 | 1.30e-13 | 7.67e-11 |
| 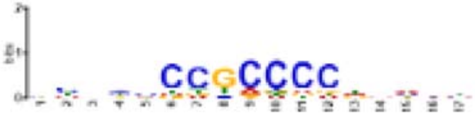 | uniprobe mouse | UP00033_2 | Zfp410_secondary | 1.75e-13 | 1.04e-10 |
| 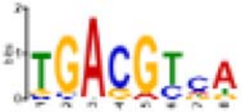 | JASPAR CORE 2014 vertebrates | MA0018.2 | CREB1 | 3.29e-13 | 1.94e-10 |
| 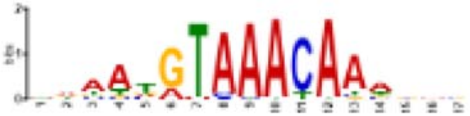 | uniprobe mouse | UP00025_1 | Foxk1_primary | 3.38e-13 | 2.00e-10 |
| 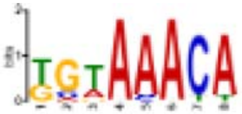 | JASPAR CORE 2014 vertebrates | MA0157.1 | FOXO3 | 8.57e-13 | 5.06e-10 |
| 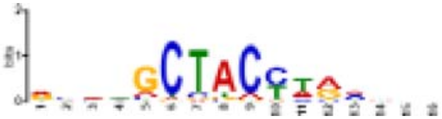 | uniprobe mouse | UP00027_2 | Osr1_secondary | 9.39e-13 | 5.55e-10 |
| 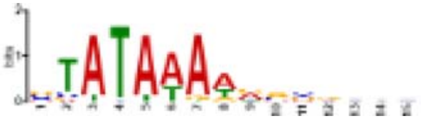 | JASPAR CORE 2014 vertebrates | MA0108.2 | TBP | 1.58e-12 | 9.32e-10 |
| 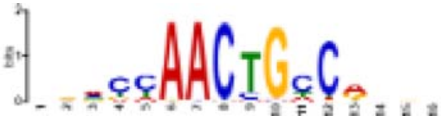 | uniprobe mouse | UP00092_2 | Myb_secondary | 1.76e-12 | 1.04e-9 |
| 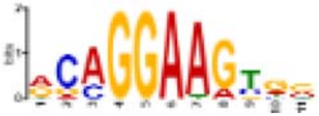 | JASPAR CORE 2014 vertebrates | MA0475.1 | FLI1 | 2.11e-12 | 1.24e-9 |
| 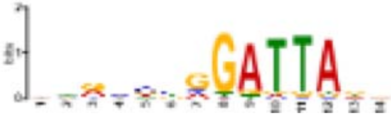 | uniprobe mouse | UP00089_2 | Tcf1_secondary | 2.77e-12 | 1.64e-9 |
| 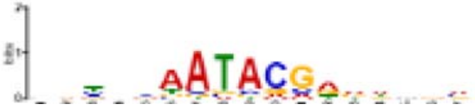 | uniprobe mouse | UP00059_2 | Arid5a_secondary | 3.82e-12 | 2.26e-9 |

| | | | | | |
|---|---|---|---|---|---|
| 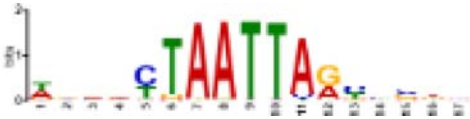 | uniprobe mouse | UP00247_1 | Pax4_3989.2 | 7.15e-12 | 4.23e-9 |
| 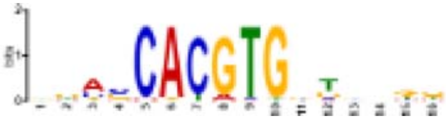 | uniprobe mouse | UP00060_1 | Max_primary | 9.08e-12 | 5.37e-9 |
| 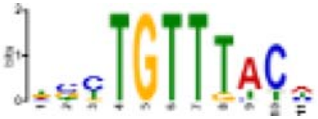 | JASPAR CORE 2014 vertebrates | MA0480.1 | Foxo1 | 1.01e-11 | 5.97e-9 |
| 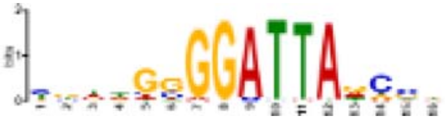 | uniprobe mouse | UP00176_1 | Crx_3485.1 | 1.93e-11 | 1.14e-8 |
| 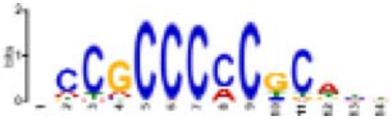 | uniprobe mouse | UP00007_1 | Egr1_primary | 2.00e-11 | 1.18e-8 |
| 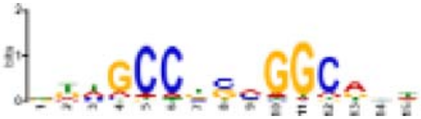 | uniprobe mouse | UP00010_2 | Tcfap2b_secondary | 2.99e-11 | 1.77e-8 |
| 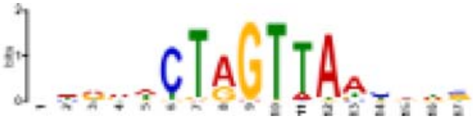 | uniprobe mouse | UP00161_1 | Hmbox1_2674.1 | 5.59e-11 | 3.30e-8 |
| 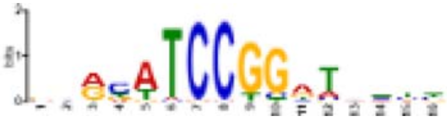 | uniprobe mouse | UP00038_1 | Spdef_primary | 6.61e-11 | 3.91e-8 |
| 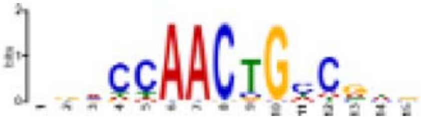 | uniprobe mouse | UP00081_2 | Mybl1_secondary | 1.08e-10 | 6.40e-8 |
| 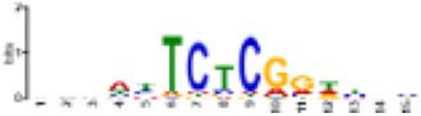 | uniprobe mouse | UP00018_2 | Irf4_secondary | 1.15e-10 | 6.81e-8 |
| 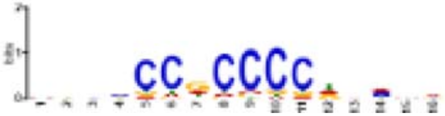 | uniprobe mouse | UP00043_2 | Bcl6b_secondary | 1.82e-10 | 1.08e-7 |

| | | | | | |
|---|---|---|---|---|---|
| 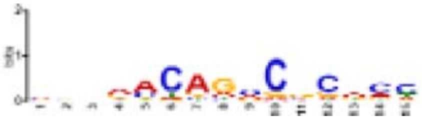 | uniprobe mouse | UP00036_2 | Myf6_secondary | 2.05e-10 | 1.21e-7 |
| 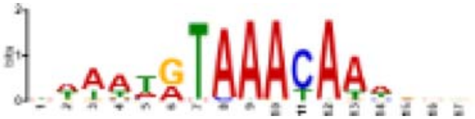 | uniprobe mouse | UP00061_1 | Foxl1_primary | 2.23e-10 | 1.32e-7 |
| 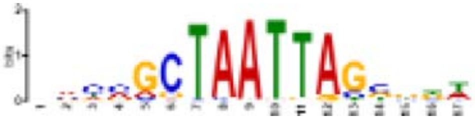 | uniprobe mouse | UP00132_1 | Evx2_2645.3 | 3.47e-10 | 2.05e-7 |
| 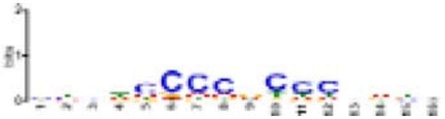 | uniprobe mouse | UP00099_2 | Ascl2_secondary | 5.37e-10 | 3.17e-7 |
| 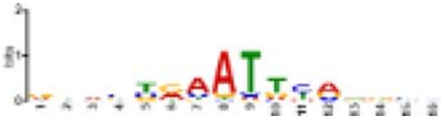 | uniprobe mouse | UP00064_2 | Sox18_secondary | 1.43e-9 | 8.48e-7 |
| 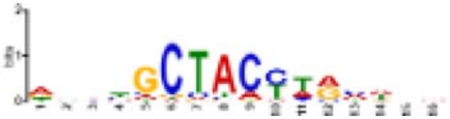 | uniprobe mouse | UP00052_2 | Osr2_secondary | 2.05e-9 | 1.21e-6 |
| 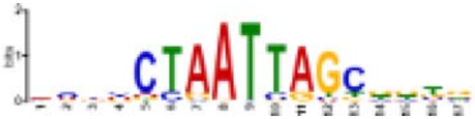 | uniprobe mouse | UP00184_1 | Lhx8_2247.2 | 2.57e-9 | 1.52e-6 |
| 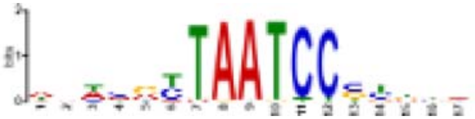 | uniprobe mouse | UP00112_1 | Gsc_2327.3 | 2.97e-9 | 1.76e-6 |
| 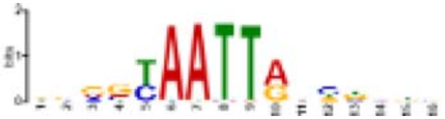 | uniprobe mouse | UP00230_1 | Dlx5_3419.2 | 3.97e-9 | 2.34e-6 |
| 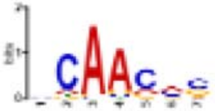 | JASPAR CORE 2014 vertebrates | MA0133.1 | BRCA1 | 4.06e-9 | 2.40e-6 |
| 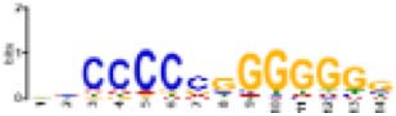 | uniprobe mouse | UP00102_1 | Zic1_primary | 4.51e-9 | 2.66e-6 |
| 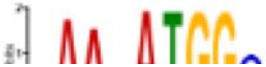 | JASPAR CORE 2014 | MA0095.2 | YY1 | 5.22e- | 3.08e-6 |

| | | | | | |
|---|---|---|---|---|---|
| 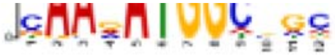 | vertebrates | [illegible] | [illegible] | [illegible] 9 | [illegible] |
| 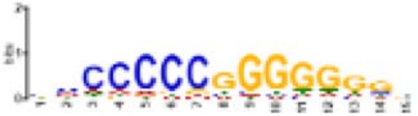 | uniprobe mouse | UP00006_1 | Zic3_primary | 6.33e-9 | 3.74e-6 |
| 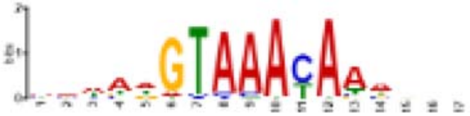 | uniprobe mouse | UP00073_1 | Foxa2_primary | 6.58e-9 | 3.89e-6 |
| 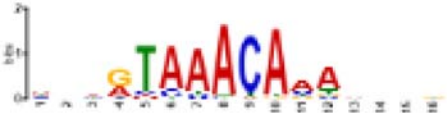 | uniprobe mouse | UP00041_1 | Foxj1_primary | 7.98e-9 | 4.72e-6 |
| 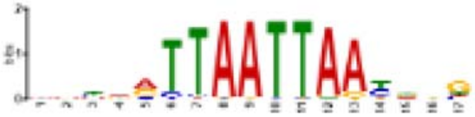 | uniprobe mouse | UP00237_1 | Otp_3496.1 | 1.05e-8 | 6.23e-6 |
| 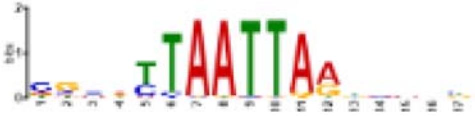 | uniprobe mouse | UP00187_1 | Alx4_1744.1 | 1.15e-8 | 6.80e-6 |
| 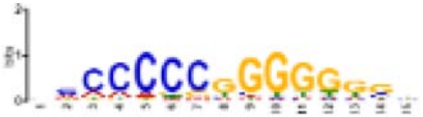 | uniprobe mouse | UP00057_1 | Zic2_primary | 1.24e-8 | 7.34e-6 |
| 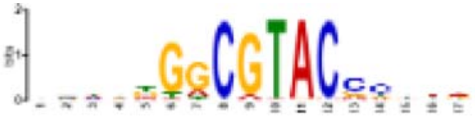 | uniprobe mouse | UP00094_1 | Zfp128_primary | 1.60e-8 | 9.45e-6 |
| 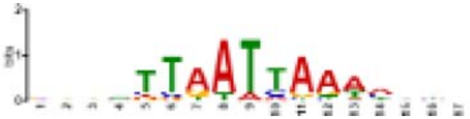 | uniprobe mouse | UP00078_1 | Arid3a_primary | 1.65e-8 | 9.76e-6 |
| 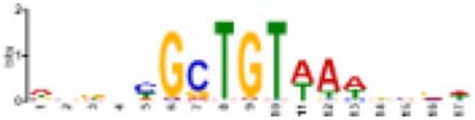 | uniprobe mouse | UP00193_1 | Rhox11_1765.2 | 3.31e-8 | 1.95e-5 |
| 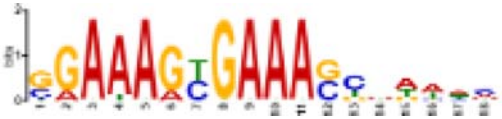 | JASPAR CORE 2014 vertebrates | MA0051.1 | IRF2 | 3.88e-8 | 2.29e-5 |
| 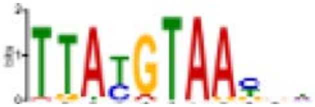 | JASPAR CORE 2014 vertebrates | MA0025.1 | NFIL3 | 4.21e-8 | 2.49e-5 |

| | | | | | |
|---|---|---|---|---|---|
| 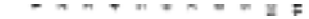 | | | | | |
| 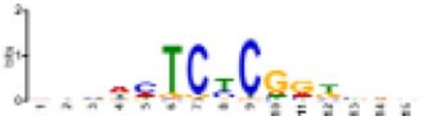 | uniprobe mouse | UP00011_2 | Irf6_secondary | 6.44e-8 | 3.80e-5 |
| 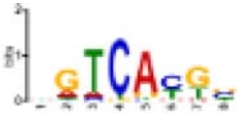 | JASPAR CORE 2014 vertebrates | MA0067.1 | Pax2 | 7.27e-8 | 4.30e-5 |
| 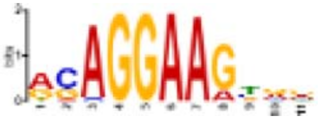 | JASPAR CORE 2014 vertebrates | MA0474.1 | Erg | 8.33e-8 | 4.92e-5 |
| 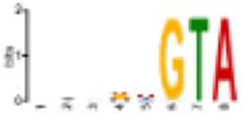 | JASPAR CORE 2014 vertebrates | MA0032.1 | FOXC1 | 1.22e-7 | 7.18e-5 |
| 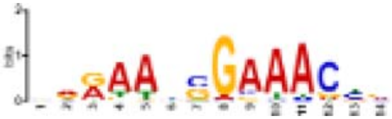 | uniprobe mouse | UP00086_1 | Irf3_primary | 1.27e-7 | 7.52e-5 |
| 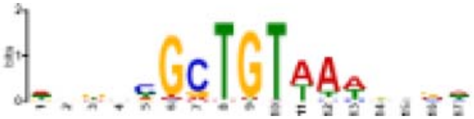 | uniprobe mouse | UP00193_2 | Rhox11_2205.1 | 1.37e-7 | 8.09e-5 |
| 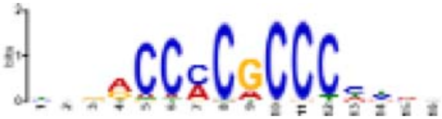 | uniprobe mouse | UP00093_1 | Klf7_primary | 1.47e-7 | 8.68e-5 |
| 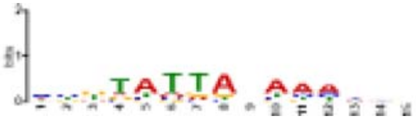 | uniprobe mouse | UP00054_2 | Tcf7_secondary | 1.59e-7 | 9.39e-5 |
| 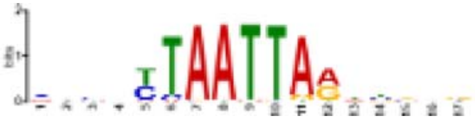 | uniprobe mouse | UP00257_1 | Shox2_2641.2 | 1.82e-7 | 1.07e-4 |
| 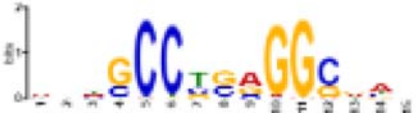 | uniprobe mouse | UP00028_1 | Tcfap2e_primary | 2.19e-7 | 1.29e-4 |
| 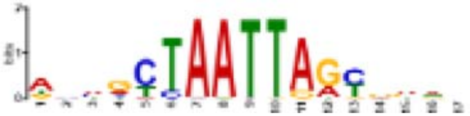 | uniprobe mouse | UP00136_1 | Prrx2_3072.1 | 2.67e-7 | 1.58e-4 |
| 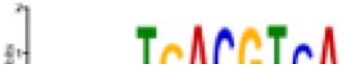 | | | | 2.89e- | |

| | | | | | |
|---|---|---|---|---|---|
| 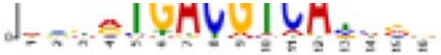 | uniprobe mouse | UP00020_1 | Atf1_primary | [illegible]-7 | 1.71e-4 |
| 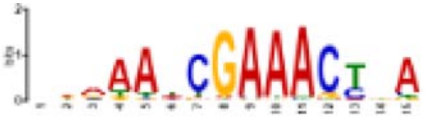 | uniprobe mouse | UP00074_1 | Isgf3g_primary | 2.99e-7 | 1.76e-4 |
| 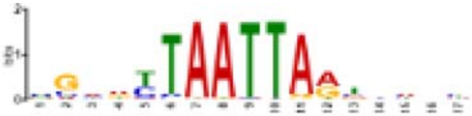 | uniprobe mouse | UP00209_1 | Cart1_0997.1 | 3.24e-7 | 1.91e-4 |
| 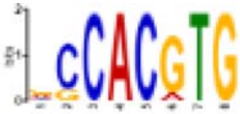 | JASPAR CORE 2014 vertebrates | MA0104.3 | Mycn | 3.77e-7 | 2.23e-4 |
| 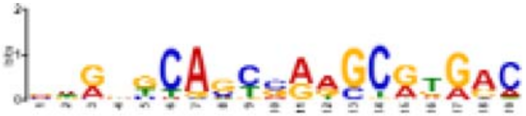 | JASPAR CORE 2014 vertebrates | MA0014.2 | PAX5 | 5.08e-7 | 3.00e-4 |
| 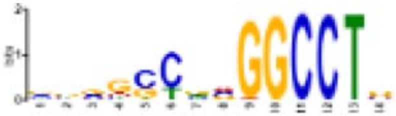 | JASPAR CORE 2014 vertebrates | MA0146.2 | Zfx | 5.16e-7 | 3.05e-4 |
| 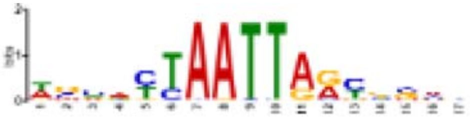 | uniprobe mouse | UP00253_1 | Rax_3443.1 | 9.14e-7 | 5.40e-4 |
| 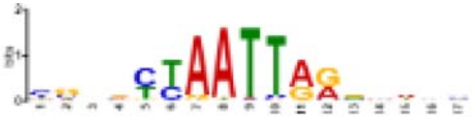 | uniprobe mouse | UP00178_1 | Og2x_3719.1 | 1.15e-6 | 6.80e-4 |
| 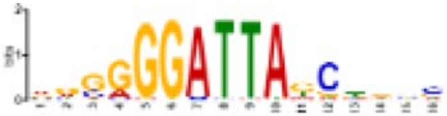 | uniprobe mouse | UP00265_1 | Pitx3_3497.2 | 1.44e-6 | 8.53e-4 |
| 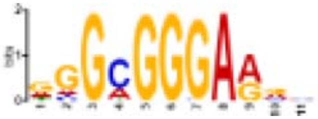 | JASPAR CORE 2014 vertebrates | MA0471.1 | E2F6 | 1.75e-6 | 1.04e-3 |
| 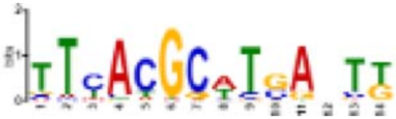 | JASPAR CORE 2014 vertebrates | MA0069.1 | Pax6 | 1.87e-6 | 1.10e-3 |
| 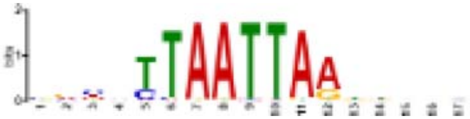 | uniprobe mouse | UP00209_2 | Cart1_1275.1 | 2.61e-6 | 1.54e-3 |

| | | | | | |
|---|---|---|---|---|---|
| 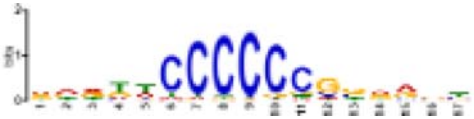 | uniprobe mouse | UP00022_2 | Zfp740_secondary | 2.64e-6 | 1.56e-3 |
| 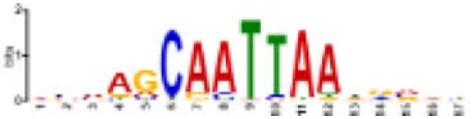 | uniprobe mouse | UP00104_1 | Hmx1_3423.1 | 3.20e-6 | 1.89e-3 |
| 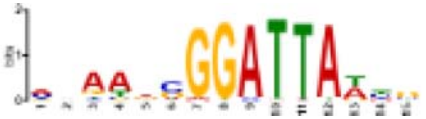 | uniprobe mouse | UP00109_1 | Obox6_3440.2 | 3.56e-6 | 2.10e-3 |
| 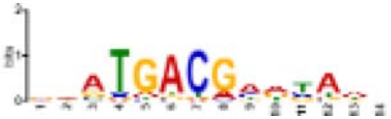 | uniprobe mouse | UP00020_2 | Atf1_secondary | 3.77e-6 | 2.22e-3 |
| 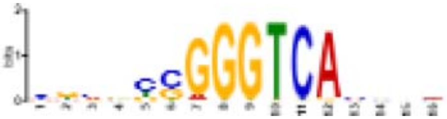 | uniprobe mouse | UP00009_2 | Nr2f2_secondary | 4.37e-6 | 2.58e-3 |
| 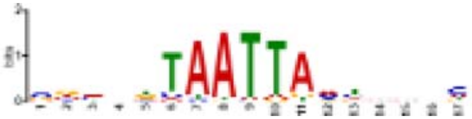 | uniprobe mouse | UP00164_1 | Hoxa7_2668.2 | 4.37e-6 | 2.58e-3 |
| 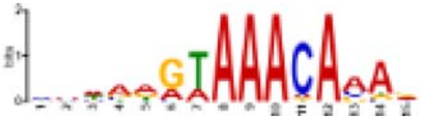 | JASPAR CORE 2014 vertebrates | MA0481.1 | FOXP1 | 5.15e-6 | 3.04e-3 |
| 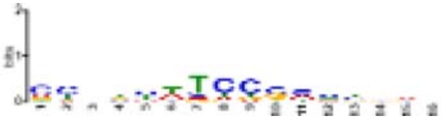 | uniprobe mouse | UP00013_2 | Gabpa_secondary | 5.21e-6 | 3.07e-3 |
| 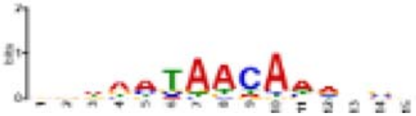 | uniprobe mouse | UP00073_2 | Foxa2_secondary | 6.17e-6 | 3.64e-3 |
| 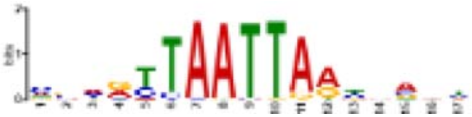 | uniprobe mouse | UP00141_1 | Vsx1_1728.1 | 8.71e-6 | 5.14e-3 |
| 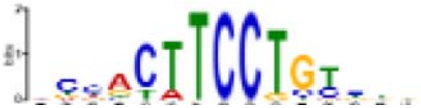 | JASPAR CORE 2014 vertebrates | MA0098.2 | Ets1 | 1.00e-5 | 5.89e-3 |

| | | | | | |
|---|---|---|---|---|---|
| 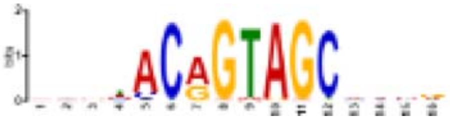 | uniprobe mouse | UP00052_1 | Osr2_primary | 1.11e-5 | 6.56e-3 |
| 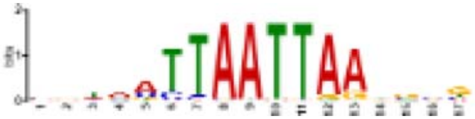 | uniprobe mouse | UP00142_1 | Uncx4.1_2281.2 | 1.29e-5 | 7.62e-3 |
| 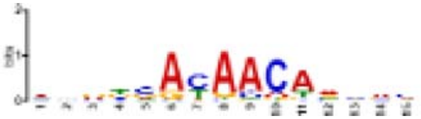 | uniprobe mouse | UP00041_2 | Foxj1_secondary | 1.58e-5 | 9.27e-3 |
| 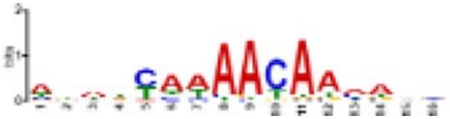 | uniprobe mouse | UP00061_2 | Foxl1_secondary | 1.69e-5 | 9.96e-3 |
| 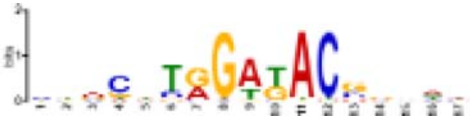 | uniprobe mouse | UP00076_2 | Rfxdc2_secondary | 1.96e-5 | 1.15e-2 |
| 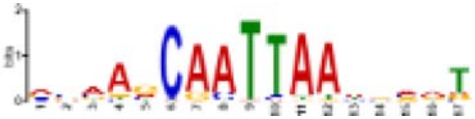 | uniprobe mouse | UP00155_1 | Hmx2_3424.3 | 2.03e-5 | 1.19e-2 |
| 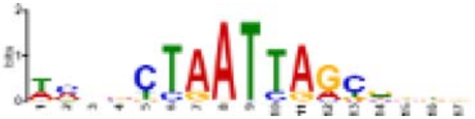 | uniprobe mouse | UP00256_2 | Lhx6_3432.1 | 2.52e-5 | 1.48e-2 |
| 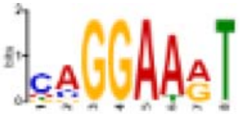 | JASPAR CORE 2014 vertebrates | MA0156.1 | FEV | 2.91e-5 | 1.70e-2 |
| 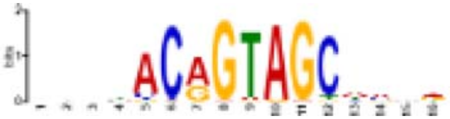 | uniprobe mouse | UP00027_1 | Osr1_primary | 4.33e-5 | 2.53e-2 |
| 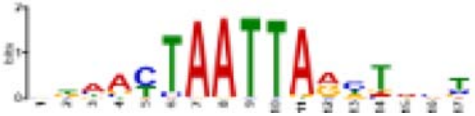 | uniprobe mouse | UP00266_1 | Prrx1_3442.1 | 4.44e-5 | 2.59e-2 |

#### Input Files

#### Sequences

| Primary Sequences | Number | Control Sequences | Number | | | | |
|---|---|---|---|---|---|---|---|
| Control Sequences | Number | | | | | | |
| ud-300expr.fa | 6525 | ud-300nonexpr.fa | 15480 | ud-300expr.fa | 6525 | ud-300nonexpr.fa | 15480 |

#### Motifs

| Database | Source | Motif Count |
|---|---|---|
| JASPAR CORE 2014 vertebrates | db/JASPAR/JASPAR_CORE_2014_vertebrates.meme | 205 |
| uniprobe mouse | db/MOUSE/uniprobe_mouse.meme | 386 |
| JASPAR CORE 2014 vertebrates | db/JASPAR/JASPAR_CORE_2014_vertebrates.meme | 205 |
| uniprobe mouse | db/MOUSE/uniprobe_mouse.meme | 386 |

**AME version**
4.11.24.11.2 (Release date: Thu May 05 14:58:55 2016 -0700Thu May 05 14:58:55 2016 -0700)
Copyright © Robert McLeay & Timothy Bailey, 2009.

**Command line summary**

```
/home/meme/meme_4.11.2/bin/ame --verbose 1 --oc . --control ud-300nonexpr.fa --bgformat 2 --bgfile background.model --scoring avg --method ranksum --pvalue-report-threshold 0.05 ud-300expr.fa db/JASPAR/JASPAR_CORE_2014_vertebrates.meme db/MOUSE/uniprobe_mouse.meme
```

Down upstream

### AME

Analysis of Motif Enrichment

For further information on how to interpret these results or to get a copy of the MEME software please access http://meme-suite.org.

If you use AME in your research, please cite the following paper:
Robert McLeay and Timothy L. Bailey, "Motif Enrichment Analysis: A unified framework and method evaluation", *BMC Bioinformatics*, **11**:165, 2010, doi:10.1186/1471-2105-11-165. [full text]

Enriched Motifs | Input Files | Program information

#### Enriched Motifs

Fixed partition size: number of primary sequences (2188)

Sequence motif score: avg_odds
Background model source: file background.model
Background model frequencies: 0.291,0.209,0.209,0.291
Total pseudocount added to a motif column: 0.25

Statistical test: Wilcoxon rank-sum test
Ranksum method: quick
Threshold *p*-value for reporting results: 0.05
Number of multiple tests for Bonferroni correction: #Motifs × #PartitionsTested = 591 × 1 = 591Fixed partition size: number of primary sequences (2188)

Sequence motif score: avg_odds
Background model source: file background.model
Background model frequencies: 0.291,0.209,0.209,0.291
Total pseudocount added to a motif column: 0.25

Statistical test: Wilcoxon rank-sum test
Ranksum method: quick
Threshold *p*-value for reporting results: 0.05
Number of multiple tests for Bonferroni correction: #Motifs × #PartitionsTested = 591 × 1 = 591

| Logo | Database | ID | Name | *p*-value | Adjusted *p*-value |
|---|---|---|---|---|---|
| 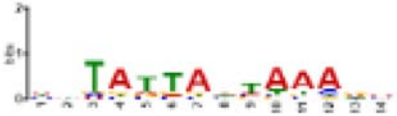 | uniprobe mouse | UP00024_2 | Glis2_secondary | 3.45e-7 | 2.04e-4 |

| | | | | | |
|---|---|---|---|---|---|
| 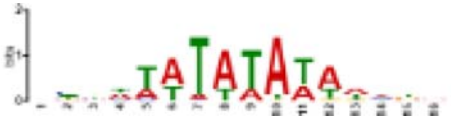 | uniprobe mouse | UP00029_1 | Tbp_primary | 2.47e-6 | 1.46e-3 |
| 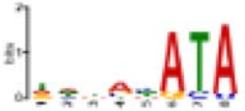 | JASPAR CORE 2014 vertebrates | MA0033.1 | FOXL1 | 8.40e-6 | 4.95e-3 |
| 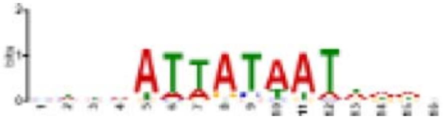 | uniprobe mouse | UP00071_1 | Sox21_primary | 1.21e-5 | 7.14e-3 |
| 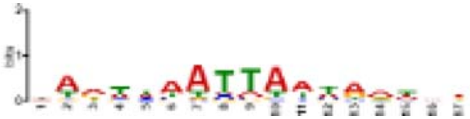 | uniprobe mouse | UP00255_1 | Dbx1_3486.1 | 2.03e-5 | 1.20e-2 |
| 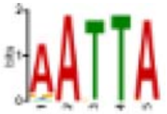 | JASPAR CORE 2014 vertebrates | MA0075.1 | Prrx2 | 2.31e-5 | 1.35e-2 |
| 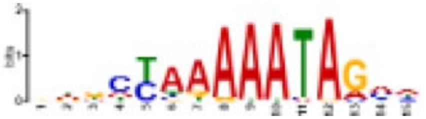 | JASPAR CORE 2014 vertebrates | MA0497.1 | MEF2C | 2.72e-5 | 1.59e-2 |
| 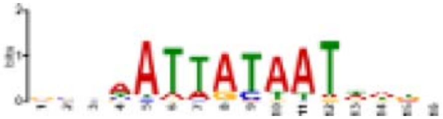 | uniprobe mouse | UP00004_1 | Sox14_primary | 3.36e-5 | 1.97e-2 |
| 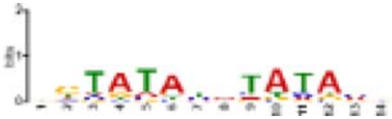 | uniprobe mouse | UP00094_2 | Zfp128_secondary | 4.50e-5 | 2.62e-2 |
| 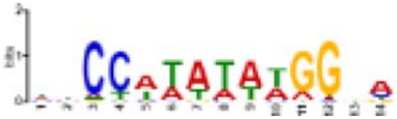 | uniprobe mouse | UP00077_1 | Srf_primary | 5.77e-5 | 3.35e-2 |
| 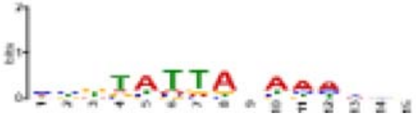 | uniprobe mouse | UP00054_2 | Tcf7_secondary | 5.86e-5 | 3.40e-2 |
| 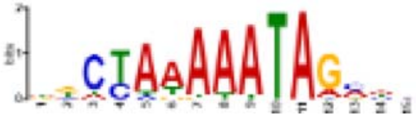 | JASPAR CORE 2014 vertebrates | MA0052.2 | MEF2A | 8.00e-5 | 4.62e-2 |

### Input Files

#### Sequences

| Primary Sequences | Number | Control Sequences | Number | | | | |
|---|---|---|---|---|---|---|---|
| Control Sequences | Number | | | | | | |
| ud300down.fa | 2188 | ud300restdown.fa | 4337 | ud300down.fa | 2188 | ud300restdown.fa | 4337 |

#### Motifs

| Database | Source | Motif Count |
|---|---|---|
| JASPAR CORE 2014 vertebrates | db/JASPAR/JASPAR_CORE_2014_vertebrates.meme | 205 |
| uniprobe mouse | db/MOUSE/uniprobe_mouse.meme | 386 |
| JASPAR CORE 2014 vertebrates | db/JASPAR/JASPAR_CORE_2014_vertebrates.meme | 205 |
| uniprobe mouse | db/MOUSE/uniprobe_mouse.meme | 386 |

**AME version**
4.11.24.11.2 (Release date: Thu May 05 14:58:55 2016 -0700Thu May 05 14:58:55 2016 -0700)
Copyright © Robert McLeay & Timothy Bailey, 2009.

**Command line summary**

```
/home/meme/meme_4.11.2/bin/ame --verbose 1 --oc . --control ud300restdown.fa --bgformat 2 --bgfile background.model --scoring avg --method ranksum --pvalue-report-threshold 0.05 ud300down.fa db/JASPAR/JASPAR_CORE_2014_vertebrates.meme db/MOUSE/uniprobe_mouse.meme
```

### AME
Analysis of Motif Enrichment



For further information on how to interpret these results or to get a copy of the MEME software please access http://meme-suite.org.

If you use AME in your research, please cite the following paper:
Robert McLeay and Timothy L. Bailey, "Motif Enrichment Analysis: A unified framework and method evaluation", *BMC Bioinformatics*, **11**:165, 2010, doi:10.1186/1471-2105-11-165. [full text]

Enriched Motifs | Input Files | Program information

#### Enriched Motifs

Fixed partition size: number of primary sequences (2188)

Sequence motif score: avg_odds
Background model source: file background.model
Background model frequencies: 0.291,0.209,0.209,0.291
Total pseudocount added to a motif column: 0.25

Statistical test: Wilcoxon rank-sum test
Ranksum method: quick
Threshold *p*-value for reporting results: 0.05
Number of multiple tests for Bonferroni correction: #Motifs × #PartitionsTested = 591 × 1 = 591Fixed partition size: number of primary sequences (2188)

Sequence motif score: avg_odds
Background model source: file background.model
Background model frequencies: 0.291,0.209,0.209,0.291
Total pseudocount added to a motif column: 0.25

Statistical test: Wilcoxon rank-sum test
Ranksum method: quick
Threshold *p*-value for reporting results: 0.05
Number of multiple tests for Bonferroni correction: #Motifs × #PartitionsTested = 591 × 1 = 591

| Logo | Database | ID | Name | *p*-value | Adjusted *p*-value |
|---|---|---|---|---|---|
| 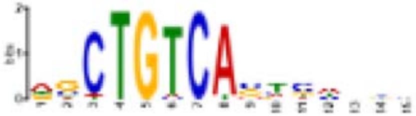 | JASPAR CORE 2014 vertebrates | MA0498.1 | Meis1 | 2.03e-15 | 1.20e-12 |

| | | | | | |
|---|---|---|---|---|---|
| 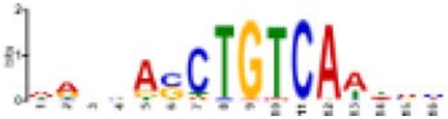 | uniprobe mouse | UP00210_1 | Mrg2_2302.1 | 6.92e-14 | 4.09e-11 |
| 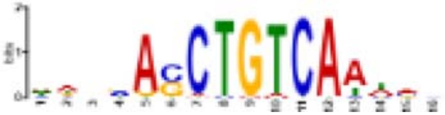 | uniprobe mouse | UP00226_1 | Mrg1_2246.2 | 9.68e-12 | 5.72e-9 |
| 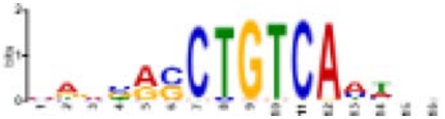 | uniprobe mouse | UP00205_1 | Pknox2_3077.2 | 1.04e-11 | 6.14e-9 |
| 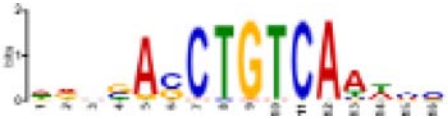 | uniprobe mouse | UP00203_1 | Pknox1_2364.2 | 1.44e-11 | 8.53e-9 |
| 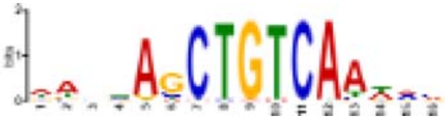 | uniprobe mouse | UP00258_1 | Tgif2_3451.1 | 2.30e-11 | 1.36e-8 |
| 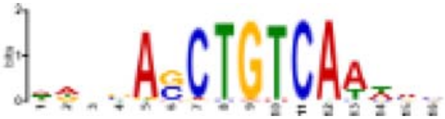 | uniprobe mouse | UP00186_1 | Meis1_2335.1 | 6.47e-11 | 3.82e-8 |
| 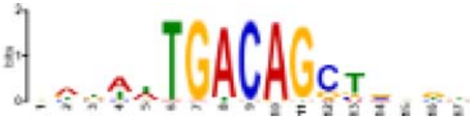 | uniprobe mouse | UP00122_1 | Tgif1_2342.2 | 9.09e-11 | 5.37e-8 |
| 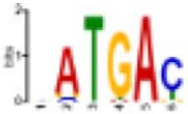 | JASPAR CORE 2014 vertebrates | MA0089.1 | NFE2L1::MafG | 9.31e-7 | 5.50e-4 |
| 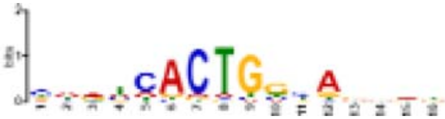 | uniprobe mouse | UP00031_2 | Zbtb3_secondary | 8.06e-6 | 4.75e-3 |
| 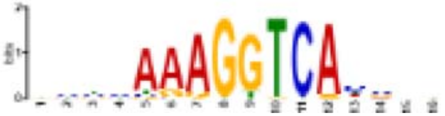 | uniprobe mouse | UP00048_1 | Rara_primary | 8.08e-5 | 4.66e-2 |

#### Input Files

#### Sequences

| Primary Sequences | Number | Control Sequences | Number | | | | |
|---|---|---|---|---|---|---|---|
| Control Sequences | Number | | | | | | |
| ud-300down.fa | 2188 | ud-300restdown.fa | 4337 | ud-300down.fa | 2188 | ud-300restdown.fa | 4337 |

#### Motifs

| Database | Source | Motif Count |
|---|---|---|
| JASPAR CORE 2014 vertebrates | db/JASPAR/JASPAR_CORE_2014_vertebrates.meme | 205 |
| uniprobe mouse | db/MOUSE/uniprobe_mouse.meme | 386 |
| JASPAR CORE 2014 vertebrates | db/JASPAR/JASPAR_CORE_2014_vertebrates.meme | 205 |
| uniprobe mouse | db/MOUSE/uniprobe_mouse.meme | 386 |

**AME version**
4.11.24.11.2 (Release date: Thu May 05 14:58:55 2016 -0700Thu May 05 14:58:55 2016 -0700)
Copyright © Robert McLeay & Timothy Bailey, 2009.

**Command line summary**

```
/home/meme/meme_4.11.2/bin/ame --verbose 1 --oc . --control ud-300restdown.fa --bgformat 2 --bgfile background.model --scoring avg --
method ranksum --pvalue-report-threshold 0.05 ud-300down.fa db/JASPAR/JASPAR_CORE_2014_vertebrates.meme db/MOUSE/uniprobe_mouse.meme
```

### AME
Analysis of Motif Enrichment

For further information on how to interpret these results or to get a copy of the MEME software please access [http://meme-suite.org](http://meme-suite.org).

If you use AME in your research, please cite the following paper:
Robert McLeay and Timothy L. Bailey, "Motif Enrichment Analysis: A unified framework and method evaluation", *BMC Bioinformatics*, **11**:165, 2010, doi:10.1186/1471-2105-11-165. [full text]

Enriched Motifs | Input Files | Program information

#### Enriched Motifs

Fixed partition size: number of primary sequences (2083)

Sequence motif score: avg_odds
Background model source: file background.model
Background model frequencies: 0.291,0.209,0.209,0.291
Total pseudocount added to a motif column: 0.25

Statistical test: Wilcoxon rank-sum test
Ranksum method: quick
Threshold *p*-value for reporting results: 0.05
Number of multiple tests for Bonferroni correction: #Motifs × #PartitionsTested = 591 × 1 = 591Fixed partition size: number of primary sequences (2083)

Sequence motif score: avg_odds
Background model source: file background.model
Background model frequencies: 0.291,0.209,0.209,0.291
Total pseudocount added to a motif column: 0.25

Statistical test: Wilcoxon rank-sum test
Ranksum method: quick
Threshold *p*-value for reporting results: 0.05
Number of multiple tests for Bonferroni correction: #Motifs × #PartitionsTested = 591 × 1 = 591

| Logo | Database | ID | Name | *p*-value | Adjusted *p*-value |
|---|---|---|---|---|---|
| 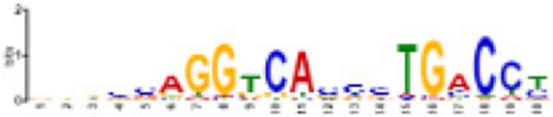 | JASPAR CORE 2014 vertebrates | MA0112.2 | ESR1 | 6.27e-8 | 3.70e-5 |

| | | | | | |
|---|---|---|---|---|---|
| 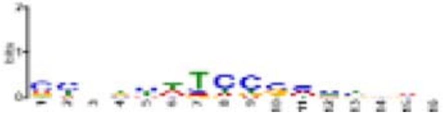 | uniprobe mouse | UP00013_2 | Gabpa_secondary | 1.34e-7 | 7.90e-5 |
| 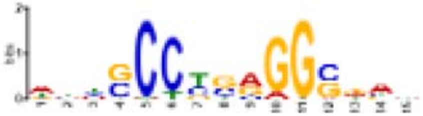 | uniprobe mouse | UP00087_1 | Tcfap2c_primary | 3.17e-7 | 1.87e-4 |
| 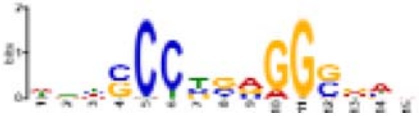 | uniprobe mouse | UP00005_1 | Tcfap2a_primary | 3.83e-7 | 2.26e-4 |
| 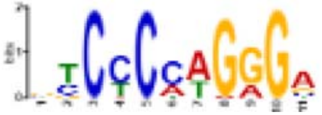 | JASPAR CORE 2014 vertebrates | MA0154.2 | EBF1 | 4.21e-7 | 2.49e-4 |
| 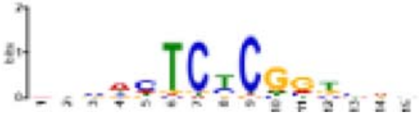 | uniprobe mouse | UP00011_2 | Irf6_secondary | 1.36e-6 | 8.02e-4 |
| 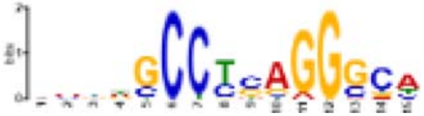 | JASPAR CORE 2014 vertebrates | MA0003.2 | TFAP2A | 1.56e-6 | 9.21e-4 |
| 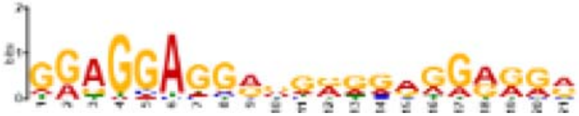 | JASPAR CORE 2014 vertebrates | MA0528.1 | ZNF263 | 2.24e-6 | 1.32e-3 |
| 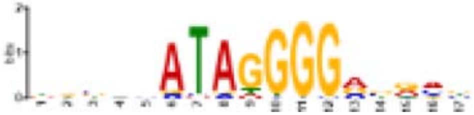 | uniprobe mouse | UP00070_2 | Gcm1_secondary | 2.29e-6 | 1.35e-3 |
| 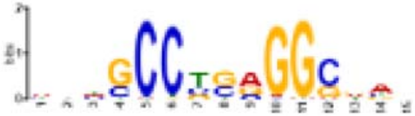 | uniprobe mouse | UP00028_1 | Tcfap2e_primary | 3.02e-6 | 1.78e-3 |
| 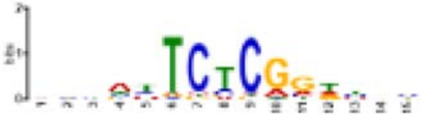 | uniprobe mouse | UP00018_2 | Irf4_secondary | 4.91e-6 | 2.90e-3 |
| 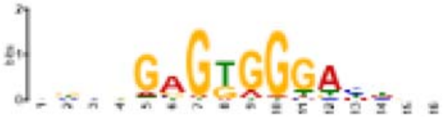 | uniprobe mouse | UP00007_2 | Egr1_secondary | 5.02e-6 | 2.96e-3 |

| | | | | | |
|---|---|---|---|---|---|
| 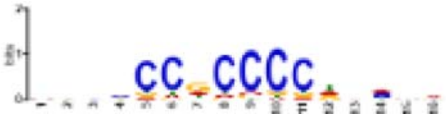 | uniprobe mouse | UP00043_2 | Bcl6b_secondary | 5.07e-6 | 2.99e-3 |
| 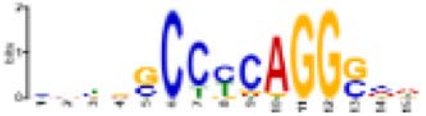 | JASPAR CORE 2014 vertebrates | MA0524.1 | TFAP2C | 5.78e-6 | 3.41e-3 |
| 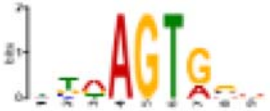 | JASPAR CORE 2014 vertebrates | MA0122.1 | Nkx3-2 | 5.86e-6 | 3.46e-3 |
| 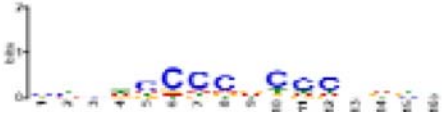 | uniprobe mouse | UP00099_2 | Ascl2_secondary | 9.15e-6 | 5.39e-3 |
| 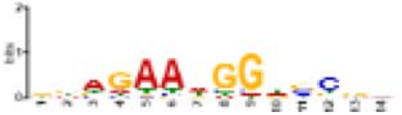 | uniprobe mouse | UP00086_2 | Irf3_secondary | 1.27e-5 | 7.51e-3 |
| 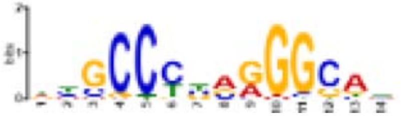 | uniprobe mouse | UP00010_1 | Tcfap2b_primary | 1.46e-5 | 8.61e-3 |
| 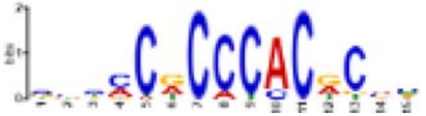 | JASPAR CORE 2014 vertebrates | MA0472.1 | EGR2 | 1.53e-5 | 9.01e-3 |
| 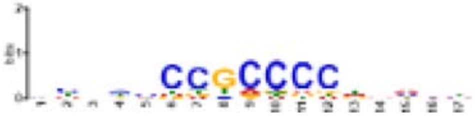 | uniprobe mouse | UP00033_2 | Zfp410_secondary | 2.52e-5 | 1.48e-2 |
| 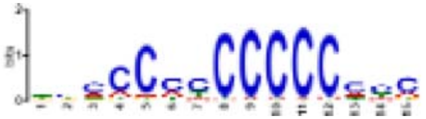 | uniprobe mouse | UP00021_1 | Zfp281_primary | 3.02e-5 | 1.77e-2 |
| 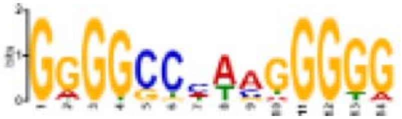 | JASPAR CORE 2014 vertebrates | MA0163.1 | PLAG1 | 6.11e-5 | 3.54e-2 |
| 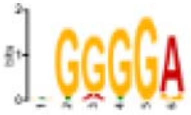 | JASPAR CORE 2014 vertebrates | MA0056.1 | MZF1_1-4 | 6.19e-5 | 3.59e-2 |

| | | | | | |
|---|---|---|---|---|---|
| 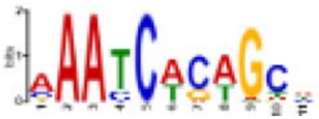 | JASPAR CORE 2014 vertebrates | MA0483.1 | Gfi1b | 7.18e-5 | 4.15e-2 |
| 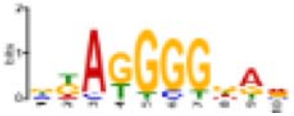 | JASPAR CORE 2014 vertebrates | MA0057.1 | MZF1_5-13 | 7.47e-5 | 4.32e-2 |
| 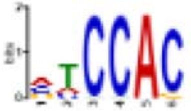 | JASPAR CORE 2014 vertebrates | MA0130.1 | ZNF354C | 7.78e-5 | 4.50e-2 |
| 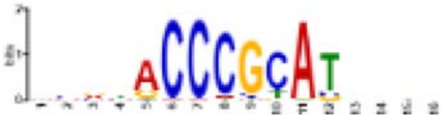 | uniprobe mouse | UP00070_1 | Gcm1_primary | 8.07e-5 | 4.66e-2 |

#### Input Files

##### Sequences

| Primary Sequences | Number | Control Sequences | Number | | | | |
|---|---|---|---|---|---|---|---|
| Control Sequences | Number | | | | | | |
| ud300up.fa | 2083 | ud300restup.fa | 4442 | ud300up.fa | 2083 | ud300restup.fa | 4442 |

##### Motifs

| Database | Source | Motif Count |
|---|---|---|
| JASPAR CORE 2014 vertebrates | db/JASPAR/JASPAR_CORE_2014_vertebrates.meme | 205 |
| uniprobe mouse | db/MOUSE/uniprobe_mouse.meme | 386 |
| JASPAR CORE 2014 vertebrates | db/JASPAR/JASPAR_CORE_2014_vertebrates.meme | 205 |
| uniprobe mouse | db/MOUSE/uniprobe_mouse.meme | 386 |

**AME version**
4.11.24.11.2 (Release date: Thu May 05 14:58:55 2016 -0700Thu May 05 14:58:55 2016 -0700)
Copyright © Robert McLeay & Timothy Bailey, 2009.

**Command line summary**

```
/home/meme/meme_4.11.2/bin/ame --verbose 1 --oc . --control ud300restup.fa --bgformat 2 --bgfile background.model --scoring avg --method ranksum --pvalue-report-threshold 0.05 ud300up.fa db/JASPAR/JASPAR_CORE_2014_vertebrates.meme db/MOUSE/uniprobe_mouse.meme
```

Type 1 downstream

### AME
Analysis of Motif Enrichment

For further information on how to interpret these results or to get a copy of the MEME software please access http://meme-suite.org.

If you use AME in your research, please cite the following paper:
Robert McLeay and Timothy L. Bailey, "Motif Enrichment Analysis: A unified framework and method evaluation", *BMC Bioinformatics*, **11**:165, 2010, doi:10.1186/1471-2105-11-165. [full text]

ENRICHED MOTIFS | INPUT FILES | PROGRAM INFORMATION

#### ENRICHED MOTIFS

Fixed partition size: number of primary sequences (250)

Sequence motif score: avg_odds
Background model source: file background.model
Background model frequencies: 0.291,0.209,0.209,0.291
Total pseudocount added to a motif column: 0.25

Statistical test: Wilcoxon rank-sum test
Ranksum method: quick
Threshold *p*-value for reporting results: 0.05
Number of multiple tests for Bonferroni correction: #Motifs × #PartitionsTested = 591 × 1 = 591Fixed partition size: number of primary sequences (250)

Sequence motif score: avg_odds
Background model source: file background.model
Background model frequencies: 0.291,0.209,0.209,0.291
Total pseudocount added to a motif column: 0.25

Statistical test: Wilcoxon rank-sum test
Ranksum method: quick
Threshold *p*-value for reporting results: 0.05
Number of multiple tests for Bonferroni correction: #Motifs × #PartitionsTested = 591 × 1 = 591

| Logo | Database | ID | Name | *p*-value | Adjusted *p*-value |
|---|---|---|---|---|---|
| 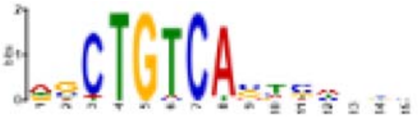 | JASPAR CORE 2014 vertebrates | MA0498.1 | Meis1 | 2.98e-7 | 1.76e-4 |

| | | | | | |
|---|---|---|---|---|---|
| 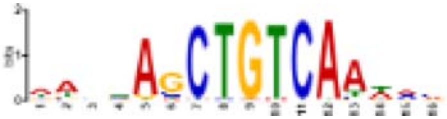 | uniprobe mouse | UP00258_1 | Tgif2_3451.1 | 5.61e-7 | 3.32e-4 |
| 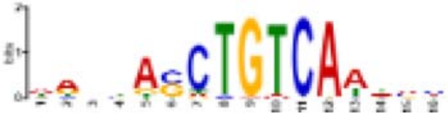 | uniprobe mouse | UP00210_1 | Mrg2_2302.1 | 5.07e-6 | 2.99e-3 |
| 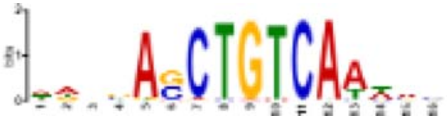 | uniprobe mouse | UP00186_1 | Meis1_2335.1 | 1.60e-5 | 9.39e-3 |
| 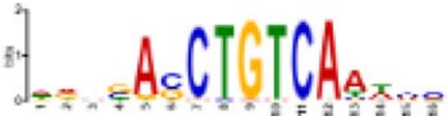 | uniprobe mouse | UP00203_1 | Pknox1_2364.2 | 2.16e-5 | 1.27e-2 |
| 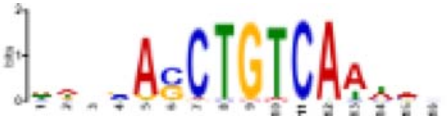 | uniprobe mouse | UP00226_1 | Mrg1_2246.2 | 2.48e-5 | 1.45e-2 |
| 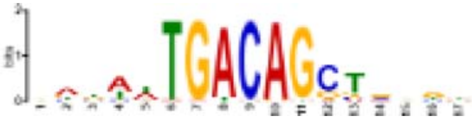 | uniprobe mouse | UP00122_1 | Tgif1_2342.2 | 5.99e-5 | 3.48e-2 |

#### Input Files

##### Sequences

| Primary Sequences | Number | Control Sequences | Number | | | | |
|---|---|---|---|---|---|---|---|
| Control Sequences | Number | | | | | | |
| ud-300type1.fa | 250 | ud-300rest1.fa | 6275 | ud-300type1.fa | 250 | ud-300rest1.fa | 6275 |

##### Motifs

| Database | Source | Motif Count |
|---|---|---|
| JASPAR CORE 2014 vertebrates | db/JASPAR/JASPAR_CORE_2014_vertebrates.meme | 205 |
| uniprobe mouse | db/MOUSE/uniprobe_mouse.meme | 386 |
| JASPAR CORE 2014 vertebrates | db/JASPAR/JASPAR_CORE_2014_vertebrates.meme | 205 |
| uniprobe mouse | db/MOUSE/uniprobe_mouse.meme | 386 |

**AME version**

4.11.24.11.2 (Release date: Thu May 05 14:58:55 2016 -0700Thu May 05 14:58:55 2016 -0700)

Copyright © Robert McLeay & Timothy Bailey, 2009.

**Command line summary**

```
/home/meme/meme_4.11.2/bin/ame --verbose 1 --oc . --control ud-300rest1.fa --bgformat 2 --bgfile background.model --scoring avg --method ranksum --pvalue-report-threshold 0.05 ud-300type1.fa db/JASPAR/JASPAR_CORE_2014_vertebrates.meme db/MOUSE/uniprobe_mouse.meme
```

Type 2 downstream

### AME
Analysis of Motif Enrichment

For further information on how to interpret these results or to get a copy of the MEME software please access http://meme-suite.org.

If you use AME in your research, please cite the following paper:
Robert McLeay and Timothy L. Bailey, "Motif Enrichment Analysis: A unified framework and method evaluation", *BMC Bioinformatics*, **11**:165, 2010, doi:10.1186/1471-2105-11-165. [full text]

ENRICHED MOTIFS | INPUT FILES | PROGRAM INFORMATION

#### ENRICHED MOTIFS

Fixed partition size: number of primary sequences (596)

Sequence motif score: avg_odds
Background model source: file background.model
Background model frequencies: 0.291,0.209,0.209,0.291
Total pseudocount added to a motif column: 0.25

Statistical test: Wilcoxon rank-sum test
Ranksum method: quick
Threshold *p*-value for reporting results: 0.05
Number of multiple tests for Bonferroni correction: #Motifs × #PartitionsTested = 591 × 1 = 591Fixed partition size: number of primary sequences (596)

Sequence motif score: avg_odds
Background model source: file background.model
Background model frequencies: 0.291,0.209,0.209,0.291
Total pseudocount added to a motif column: 0.25

Statistical test: Wilcoxon rank-sum test
Ranksum method: quick
Threshold *p*-value for reporting results: 0.05
Number of multiple tests for Bonferroni correction: #Motifs × #PartitionsTested = 591 × 1 = 591

| Logo | Database | ID | Name | *p*-value | Adjusted *p*-value |
|---|---|---|---|---|---|
| 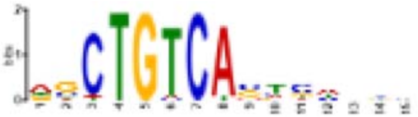 | JASPAR CORE 2014 vertebrates | MA0498.1 | Meis1 | 4.22e-6 | 2.49e-3 |

#### Input Files

##### Sequences

| Primary Sequences | Number | Control Sequences | Number | | | | |
|---|---|---|---|---|---|---|---|
| Control Sequences | Number | | | | | | |
| ud-300type2.fa | 596 | ud-300rest2.fa | 5929 | ud-300type2.fa | 596 | ud-300rest2.fa | 5929 |

##### Motifs

| Database | Source | Motif Count |
|---|---|---|
| JASPAR CORE 2014 vertebrates | db/JASPAR/JASPAR_CORE_2014_vertebrates.meme | 205 |
| uniprobe mouse | db/MOUSE/uniprobe_mouse.meme | 386 |
| JASPAR CORE 2014 vertebrates | db/JASPAR/JASPAR_CORE_2014_vertebrates.meme | 205 |
| uniprobe mouse | db/MOUSE/uniprobe_mouse.meme | 386 |

**AME version**
4.11.24.11.2 (Release date: Thu May 05 14:58:55 2016 -0700Thu May 05 14:58:55 2016 -0700)
Copyright © Robert McLeay & Timothy Bailey, 2009.

**Command line summary**

```
/home/meme/meme_4.11.2/bin/ame --verbose 1 --oc . --control ud-300rest2.fa --bgformat 2 --bgfile background.model --scoring avg --method ranksum --pvalue-report-threshold 0.05 ud-300type2.fa db/JASPAR/JASPAR_CORE_2014_vertebrates.meme db/MOUSE/uniprobe_mouse.meme
```

### AME
Analysis of Motif Enrichment

For further information on how to interpret these results or to get a copy of the MEME software please access http://meme-suite.org.

If you use AME in your research, please cite the following paper:
Robert McLeay and Timothy L. Bailey, "Motif Enrichment Analysis: A unified framework and method evaluation", *BMC Bioinformatics*, **11**:165, 2010, doi:10.1186/1471-2105-11-165. [full text]

Enriched Motifs | Input Files | Program information

#### Enriched Motifs

Fixed partition size: number of primary sequences (329)

Sequence motif score: avg_odds
Background model source: file background.model
Background model frequencies: 0.291,0.209,0.209,0.291
Total pseudocount added to a motif column: 0.25

Statistical test: Wilcoxon rank-sum test
Ranksum method: quick
Threshold *p*-value for reporting results: 0.05
Number of multiple tests for Bonferroni correction: #Motifs × #PartitionsTested = 591 × 1 = 591Fixed partition size: number of primary sequences (329)

Sequence motif score: avg_odds
Background model source: file background.model
Background model frequencies: 0.291,0.209,0.209,0.291
Total pseudocount added to a motif column: 0.25

Statistical test: Wilcoxon rank-sum test
Ranksum method: quick
Threshold *p*-value for reporting results: 0.05
Number of multiple tests for Bonferroni correction: #Motifs × #PartitionsTested = 591 × 1 = 591

| Logo | Database | ID | Name | *p*-value | Adjusted *p*-value |
|---|---|---|---|---|---|
| 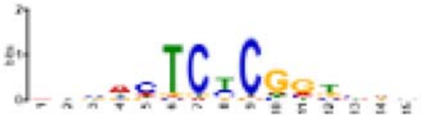 | uniprobe mouse | UP00011_2 | Irf6_secondary | 2.43e-5 | 1.42e-2 |

#### INPUT FILES

##### Sequences

| Primary Sequences | Number | Control Sequences | Number | | | | |
|---|---|---|---|---|---|---|---|
| Control Sequences | Number | | | | | | |
| ud-300type4.fa | 329 | ud-300rest4.fa | 6196 | ud-300type4.fa | 329 | ud-300rest4.fa | 6196 |

##### Motifs

| Database | Source | Motif Count |
|---|---|---|
| JASPAR CORE 2014 vertebrates | db/JASPAR/JASPAR_CORE_2014_vertebrates.meme | 205 |
| uniprobe mouse | db/MOUSE/uniprobe_mouse.meme | 386 |
| JASPAR CORE 2014 vertebrates | db/JASPAR/JASPAR_CORE_2014_vertebrates.meme | 205 |
| uniprobe mouse | db/MOUSE/uniprobe_mouse.meme | 386 |

**AME version**
4.11.24.11.2 (Release date: Thu May 05 14:58:55 2016 -0700Thu May 05 14:58:55 2016 -0700)
Copyright © Robert McLeay & Timothy Bailey, 2009.

**Command line summary**

```
/home/meme/meme_4.11.2/bin/ame --verbose 1 --oc . --control ud-300rest4.fa --bgformat 2 --bgfile background.model --scoring avg --method ranksum --pvalue-report-threshold 0.05 ud-300type4.fa db/JASPAR/JASPAR_CORE_2014_vertebrates.meme db/MOUSE/uniprobe_mouse.meme
```

### AME
Analysis of Motif Enrichment

For further information on how to interpret these results or to get a copy of the MEME software please access http://meme-suite.org.

If you use AME in your research, please cite the following paper:
Robert McLeay and Timothy L. Bailey, "Motif Enrichment Analysis: A unified framework and method evaluation", *BMC Bioinformatics*, **11**:165, 2010, doi:10.1186/1471-2105-11-165. [full text]

ENRICHED MOTIFS | INPUT FILES | PROGRAM INFORMATION

#### ENRICHED MOTIFS

Fixed partition size: number of primary sequences (166)

Sequence motif score: avg_odds
Background model source: file background.model
Background model frequencies: 0.291,0.209,0.209,0.291
Total pseudocount added to a motif column: 0.25

Statistical test: Wilcoxon rank-sum test
Ranksum method: quick
Threshold *p*-value for reporting results: 0.05
Number of multiple tests for Bonferroni correction: #Motifs × #PartitionsTested = 591 × 1 = 591Fixed partition size: number of primary sequences (166)

Sequence motif score: avg_odds
Background model source: file background.model
Background model frequencies: 0.291,0.209,0.209,0.291
Total pseudocount added to a motif column: 0.25

Statistical test: Wilcoxon rank-sum test
Ranksum method: quick
Threshold *p*-value for reporting results: 0.05
Number of multiple tests for Bonferroni correction: #Motifs × #PartitionsTested = 591 × 1 = 591

| Logo | Database | ID | Name | *p*-value | Adjusted *p*-value |
|---|---|---|---|---|---|
| 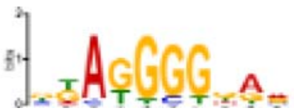 | JASPAR CORE 2014 vertebrates | MA0057.1 | MZF1_5-13 | 3.51e-7 | 2.08e-4 |

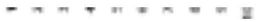

| | | | | | |
|---|---|---|---|---|---|
| 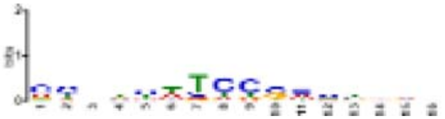 | uniprobe mouse | UP00013_2 | Gabpa_secondary | 9.17e-6 | 5.41e-3 |
| 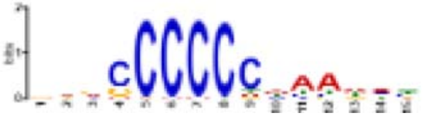 | uniprobe mouse | UP00047_1 | Zbtb7b_primary | 9.35e-6 | 5.51e-3 |
| 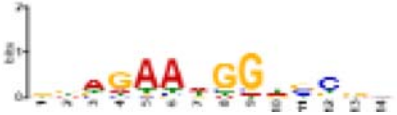 | uniprobe mouse | UP00086_2 | Irf3_secondary | 1.14e-5 | 6.74e-3 |
| 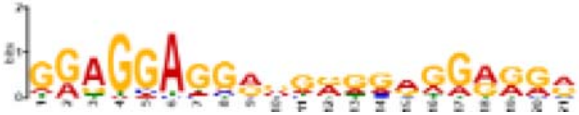 | JASPAR CORE 2014 vertebrates | MA0528.1 | ZNF263 | 2.66e-5 | 1.56e-2 |
| 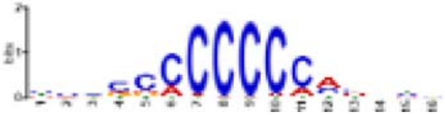 | uniprobe mouse | UP00022_1 | Zfp740_primary | 2.67e-5 | 1.56e-2 |
| 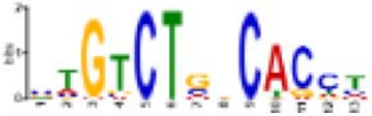 | JASPAR CORE 2014 vertebrates | MA0513.1 | SMAD2::SMAD3::SMAD4 | 3.19e-5 | 1.87e-2 |
| 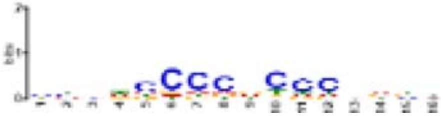 | uniprobe mouse | UP00099_2 | Ascl2_secondary | 3.65e-5 | 2.14e-2 |

#### Input Files

##### Sequences

| Primary Sequences | Number | Control Sequences | Number | | | | |
|---|---|---|---|---|---|---|---|
| **Control Sequences** | **Number** | | | | | | |
| ud300type6.fa | 166 | ud300rest6.fa | 6359 | ud300type6.fa | 166 | ud300rest6.fa | 6359 |

##### Motifs

| Database | Source | Motif Count |
|---|---|---|
| JASPAR CORE 2014 vertebrates | db/JASPAR/JASPAR_CORE_2014_vertebrates.meme | 205 |
| uniprobe mouse | db/MOUSE/uniprobe_mouse.meme | 386 |
| JASPAR CORE 2014 vertebrates | db/JASPAR/JASPAR_CORE_2014_vertebrates.meme | 205 |
| uniprobe mouse | db/MOUSE/uniprobe_mouse.meme | 386 |

**AME version**
4.11.24.11.2 (Release date: Thu May 05 14:58:55 2016 -0700Thu May 05 14:58:55 2016 -0700)
Copyright © Robert McLeay & Timothy Bailey, 2009.

**Command line summary**

```
/home/meme/meme_4.11.2/bin/ame --verbose 1 --oc . --control ud300rest6.fa --bgformat 2 --bgfile background.model --scoring avg --method ranksum --pvalue-report-threshold 0.05 ud300type6.fa db/JASPAR/JASPAR_CORE_2014_vertebrates.meme db/MOUSE/uniprobe_mouse.meme
```

### AME

Analysis of Motif Enrichment

For further information on how to interpret these results or to get a copy of the MEME software please access http://meme-suite.org.

If you use AME in your research, please cite the following paper:
Robert McLeay and Timothy L. Bailey, "Motif Enrichment Analysis: A unified framework and method evaluation", *BMC Bioinformatics*, **11**:165, 2010, doi:10.1186/1471-2105-11-165. [full text]

Enriched Motifs | Input Files | Program information

#### Enriched Motifs

Fixed partition size: number of primary sequences (1240)

Sequence motif score: avg_odds
Background model source: file background.model
Background model frequencies: 0.291,0.209,0.209,0.291
Total pseudocount added to a motif column: 0.25

Statistical test: Wilcoxon rank-sum test
Ranksum method: quick
Threshold *p*-value for reporting results: 0.05
Number of multiple tests for Bonferroni correction: #Motifs × #PartitionsTested = 591 × 1 = 591Fixed partition size: number of primary sequences (1240)

Sequence motif score: avg_odds
Background model source: file background.model
Background model frequencies: 0.291,0.209,0.209,0.291
Total pseudocount added to a motif column: 0.25

Statistical test: Wilcoxon rank-sum test
Ranksum method: quick
Threshold *p*-value for reporting results: 0.05
Number of multiple tests for Bonferroni correction: #Motifs × #PartitionsTested = 591 × 1 = 591

| Logo | Database | ID | Name | *p*-value | Adjusted *p*-value |
|---|---|---|---|---|---|
| 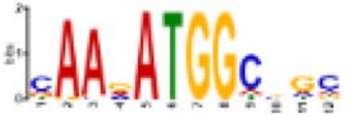 | JASPAR CORE 2014 vertebrates | MA0095.2 | YY1 | 2.56e-5 | 1.50e-2 |

#### Input Files

##### Sequences

| Primary Sequences | Number | Control Sequences | Number | | | | |
|---|---|---|---|---|---|---|---|
| Control Sequences | Number | | | | | | |
| ud-300type7.fa | 1240 | ud-300rest7.fa | 5285 | ud-300type7.fa | 1240 | ud-300rest7.fa | 5285 |

##### Motifs

| Database | Source | Motif Count |
|---|---|---|
| JASPAR CORE 2014 vertebrates | db/JASPAR/JASPAR_CORE_2014_vertebrates.meme | 205 |
| uniprobe mouse | db/MOUSE/uniprobe_mouse.meme | 386 |
| JASPAR CORE 2014 vertebrates | db/JASPAR/JASPAR_CORE_2014_vertebrates.meme | 205 |
| uniprobe mouse | db/MOUSE/uniprobe_mouse.meme | 386 |

**AME version**
4.11.24.11.2 (Release date: Thu May 05 14:58:55 2016 -0700Thu May 05 14:58:55 2016 -0700)
Copyright © Robert McLeay & Timothy Bailey, 2009.

**Command line summary**

```
/home/meme/meme_4.11.2/bin/ame --verbose 1 --oc . --control ud-300rest7.fa --bgformat 2 --bgfile background.model --scoring avg --method ranksum --pvalue-report-threshold 0.05 ud-300type7.fa db/JASPAR/JASPAR_CORE_2014_vertebrates.meme db/MOUSE/uniprobe_mouse.meme
```

Type 8 upstream

### AME

Analysis of Motif Enrichment

For further information on how to interpret these results or to get a copy of the MEME software please access http://meme-suite.org.

If you use AME in your research, please cite the following paper:
Robert McLeay and Timothy L. Bailey, "Motif Enrichment Analysis: A unified framework and method evaluation", *BMC Bioinformatics*, **11**:165, 2010, doi:10.1186/1471-2105-11-165. [full text]

Enriched Motifs | Input Files | Program information

#### Enriched Motifs

Fixed partition size: number of primary sequences (527)

Sequence motif score: avg_odds
Background model source: file background.model
Background model frequencies: 0.291,0.209,0.209,0.291
Total pseudocount added to a motif column: 0.25

Statistical test: Wilcoxon rank-sum test
Ranksum method: quick
Threshold *p*-value for reporting results: 0.05
Number of multiple tests for Bonferroni correction: #Motifs × #PartitionsTested = 591 × 1 = 591Fixed partition size: number of primary sequences (527)

Sequence motif score: avg_odds
Background model source: file background.model
Background model frequencies: 0.291,0.209,0.209,0.291
Total pseudocount added to a motif column: 0.25

Statistical test: Wilcoxon rank-sum test
Ranksum method: quick
Threshold *p*-value for reporting results: 0.05
Number of multiple tests for Bonferroni correction: #Motifs × #PartitionsTested = 591 × 1 = 591

| Logo | Database | ID | Name | *p*-value | Adjusted *p*-value |
|---|---|---|---|---|---|
| 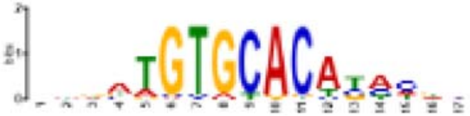 | uniprobe mouse | UP00042_1 | Gm397_primary | 5.35e-6 | 3.16e-3 |

| | | | | | |
|---|---|---|---|---|---|
| 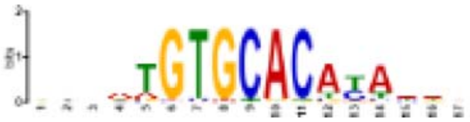 | uniprobe mouse | UP00026_1 | Zscan4_primary | 2.97e-5 | 1.74e-2 |

#### Input Files

##### Sequences

| Primary Sequences | Number | Control Sequences | Number | | | | |
|---|---|---|---|---|---|---|---|
| **Control Sequences** | **Number** | | | | | | |
| ud300type8.fa | 527 | ud300rest8.fa | 5998 | ud300type8.fa | 527 | ud300rest8.fa | 5998 |

##### Motifs

| Database | Source | Motif Count |
|---|---|---|
| JASPAR CORE 2014 vertebrates | db/JASPAR/JASPAR_CORE_2014_vertebrates.meme | 205 |
| uniprobe mouse | db/MOUSE/uniprobe_mouse.meme | 386 |
| JASPAR CORE 2014 vertebrates | db/JASPAR/JASPAR_CORE_2014_vertebrates.meme | 205 |
| uniprobe mouse | db/MOUSE/uniprobe_mouse.meme | 386 |

**AME version**
4.11.24.11.2 (Release date: Thu May 05 14:58:55 2016 -0700Thu May 05 14:58:55 2016 -0700)
Copyright © Robert McLeay & Timothy Bailey, 2009.

**Command line summary**

```
/home/meme/meme_4.11.2/bin/ame --verbose 1 --oc . --control ud300rest8.fa --bgformat 2 --bgfile background.model --scoring avg --method
ranksum --pvalue-report-threshold 0.05 ud300type8.fa db/JASPAR/JASPAR_CORE_2014_vertebrates.meme db/MOUSE/uniprobe_mouse.meme
```

Type 9 upstream

### AME
Analysis of Motif Enrichment

For further information on how to interpret these results or to get a copy of the MEME software please access http://meme-suite.org.

If you use AME in your research, please cite the following paper:
Robert McLeay and Timothy L. Bailey, "Motif Enrichment Analysis: A unified framework and method evaluation", *BMC Bioinformatics*, **11**:165, 2010, doi:10.1186/1471-2105-11-165. [full text]

ENRICHED MOTIFS | INPUT FILES | PROGRAM INFORMATION

#### ENRICHED MOTIFS

Fixed partition size: number of primary sequences (150)

Sequence motif score: avg_odds
Background model source: file background.model
Background model frequencies: 0.291,0.209,0.209,0.291
Total pseudocount added to a motif column: 0.25

Statistical test: Wilcoxon rank-sum test
Ranksum method: quick
Threshold *p*-value for reporting results: 0.05
Number of multiple tests for Bonferroni correction: #Motifs × #PartitionsTested = 591 × 1 = 591Fixed partition size: number of primary sequences (150)

Sequence motif score: avg_odds
Background model source: file background.model
Background model frequencies: 0.291,0.209,0.209,0.291
Total pseudocount added to a motif column: 0.25

Statistical test: Wilcoxon rank-sum test
Ranksum method: quick
Threshold *p*-value for reporting results: 0.05
Number of multiple tests for Bonferroni correction: #Motifs × #PartitionsTested = 591 × 1 = 591

| Logo | Database | ID | Name | *p*-value | Adjusted *p*-value |
|---|---|---|---|---|---|
| 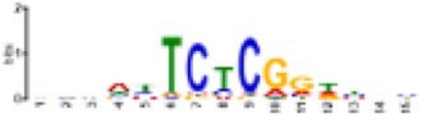 | uniprobe mouse | UP00018_2 | Irf4_secondary | 5.82e-5 | 3.38e-2 |

| | | | | | |
|---|---|---|---|---|---|
| 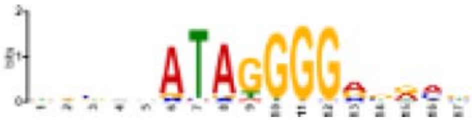 | uniprobe mouse | UP00070_2 | Gcm1_secondary | 7.34e-5 | 4.25e-2 |

#### Input Files

##### Sequences

| Primary Sequences | Number | Control Sequences | Number | | | | |
|---|---|---|---|---|---|---|---|
| Control Sequences | Number | | | | | | |
| ud300type9.fa | 150 | ud300rest9.fa | 6375 | ud300type9.fa | 150 | ud300rest9.fa | 6375 |

##### Motifs

| Database | Source | Motif Count |
|---|---|---|
| JASPAR CORE 2014 vertebrates | db/JASPAR/JASPAR_CORE_2014_vertebrates.meme | 205 |
| uniprobe mouse | db/MOUSE/uniprobe_mouse.meme | 386 |
| JASPAR CORE 2014 vertebrates | db/JASPAR/JASPAR_CORE_2014_vertebrates.meme | 205 |
| uniprobe mouse | db/MOUSE/uniprobe_mouse.meme | 386 |

**AME version**
4.11.24.11.2 (Release date: Thu May 05 14:58:55 2016 -0700Thu May 05 14:58:55 2016 -0700)
Copyright © Robert McLeay & Timothy Bailey, 2009.

**Command line summary**

```
/home/meme/meme_4.11.2/bin/ame --verbose 1 --oc . --control ud300rest9.fa --bgformat 2 --bgfile background.model --scoring avg --method ranksum --pvalue-report-threshold 0.05 ud300type9.fa db/JASPAR/JASPAR_CORE_2014_vertebrates.meme db/MOUSE/uniprobe_mouse.meme
```
