## Supplementary figures and images for "Cellular Factors Involved in Transcriptome Dynamics in Early Zebrafish Embryogenesis"

### Supplemental File S9

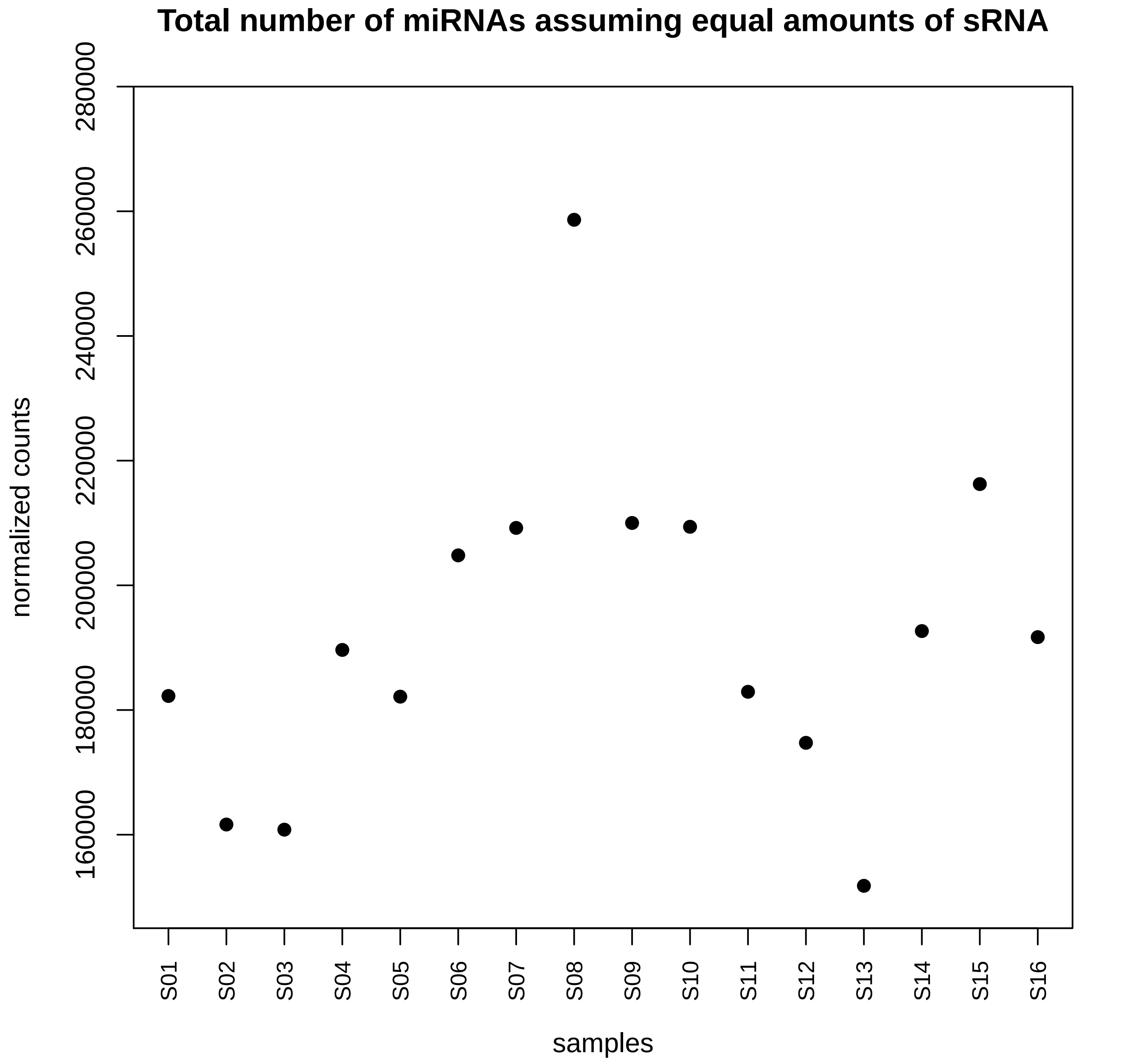

### Supplemental File S12

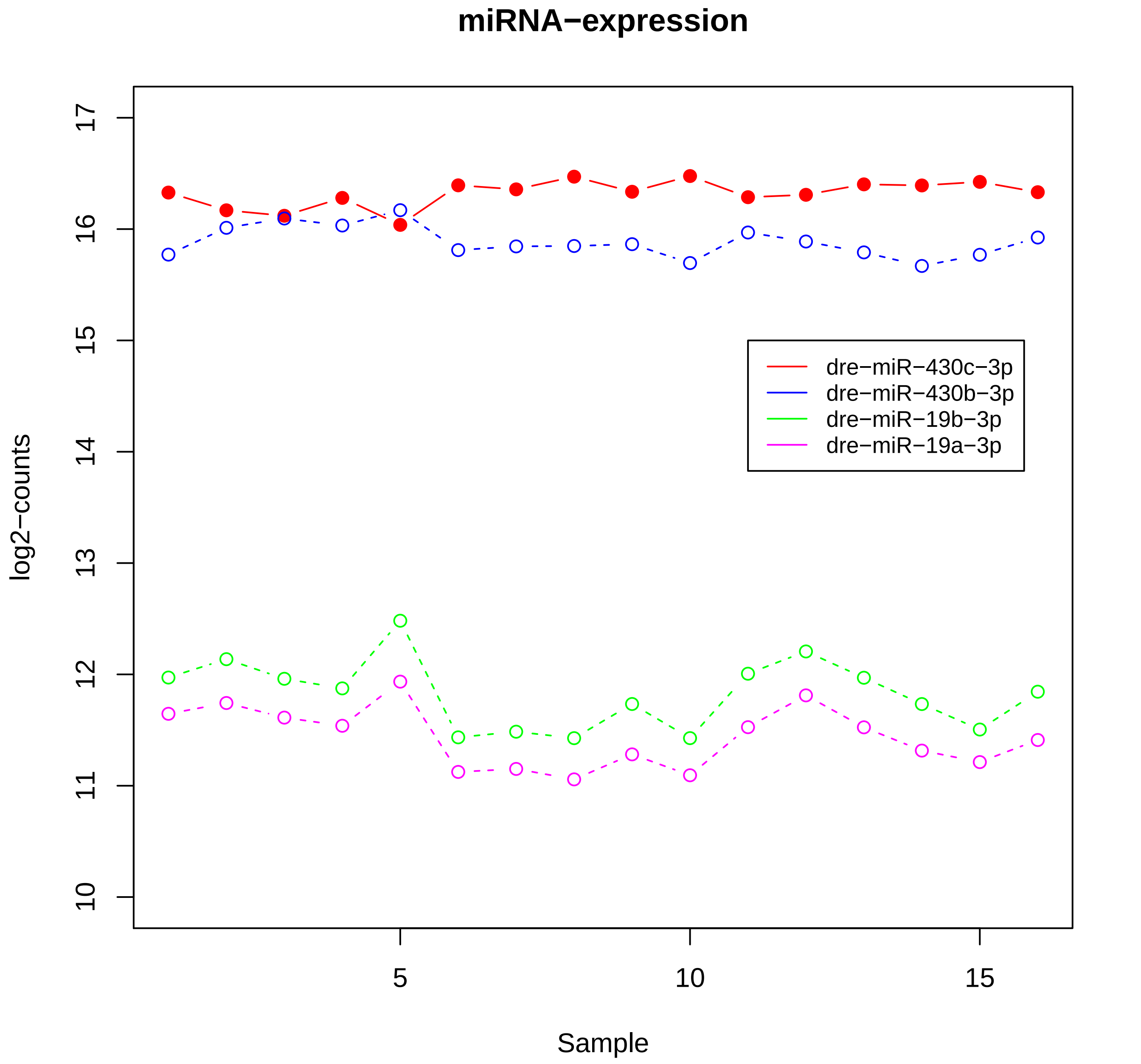

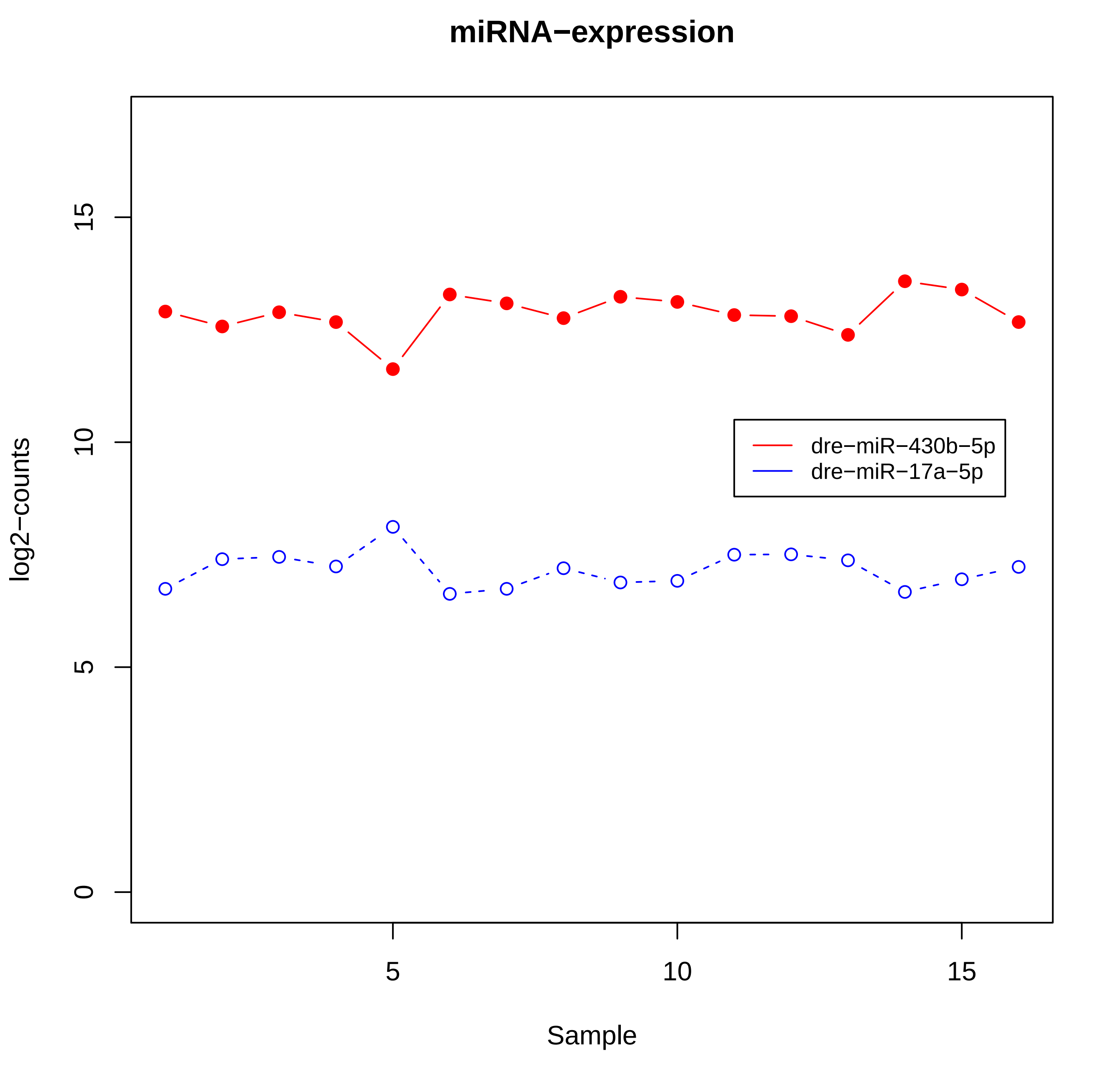

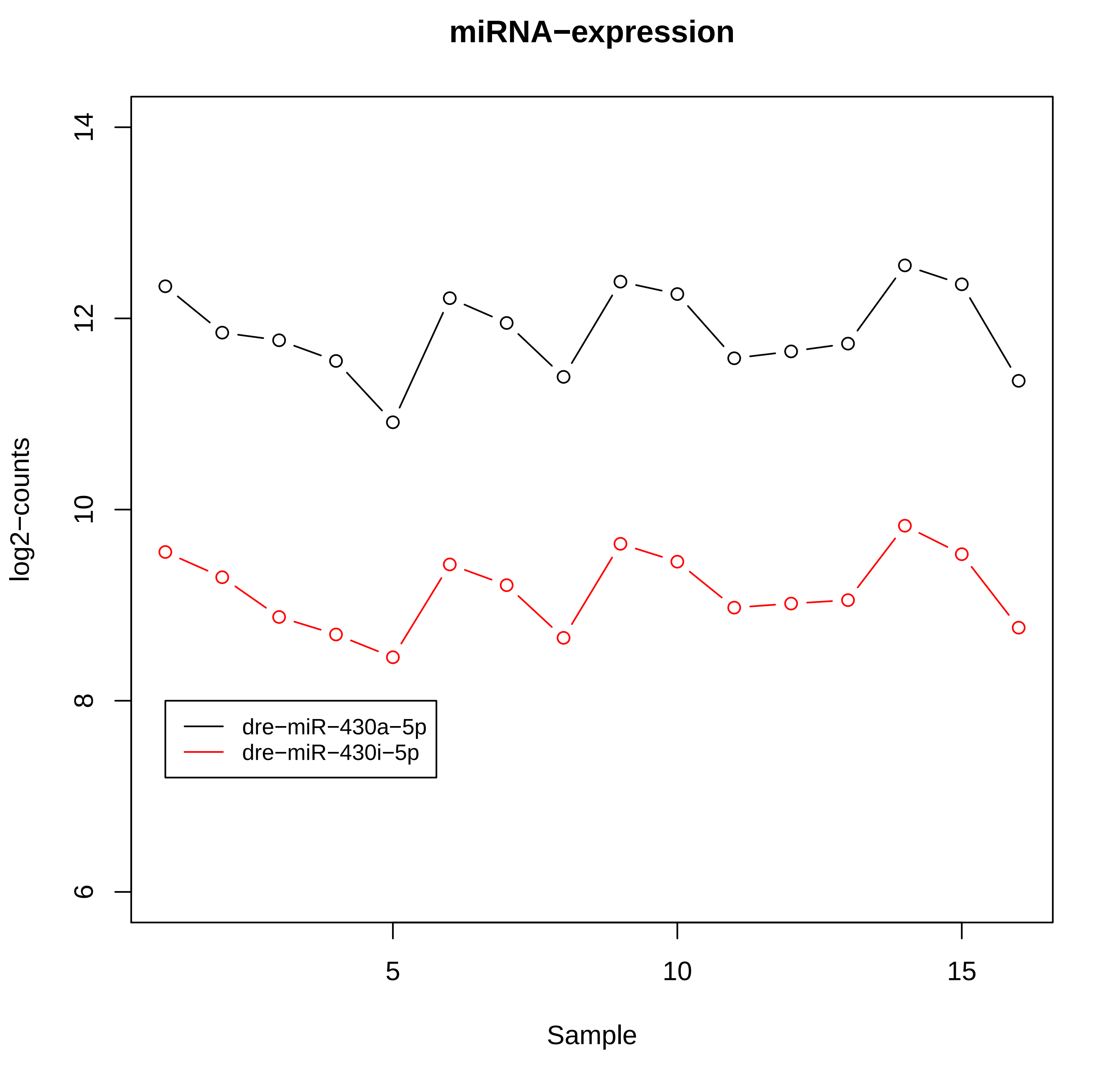

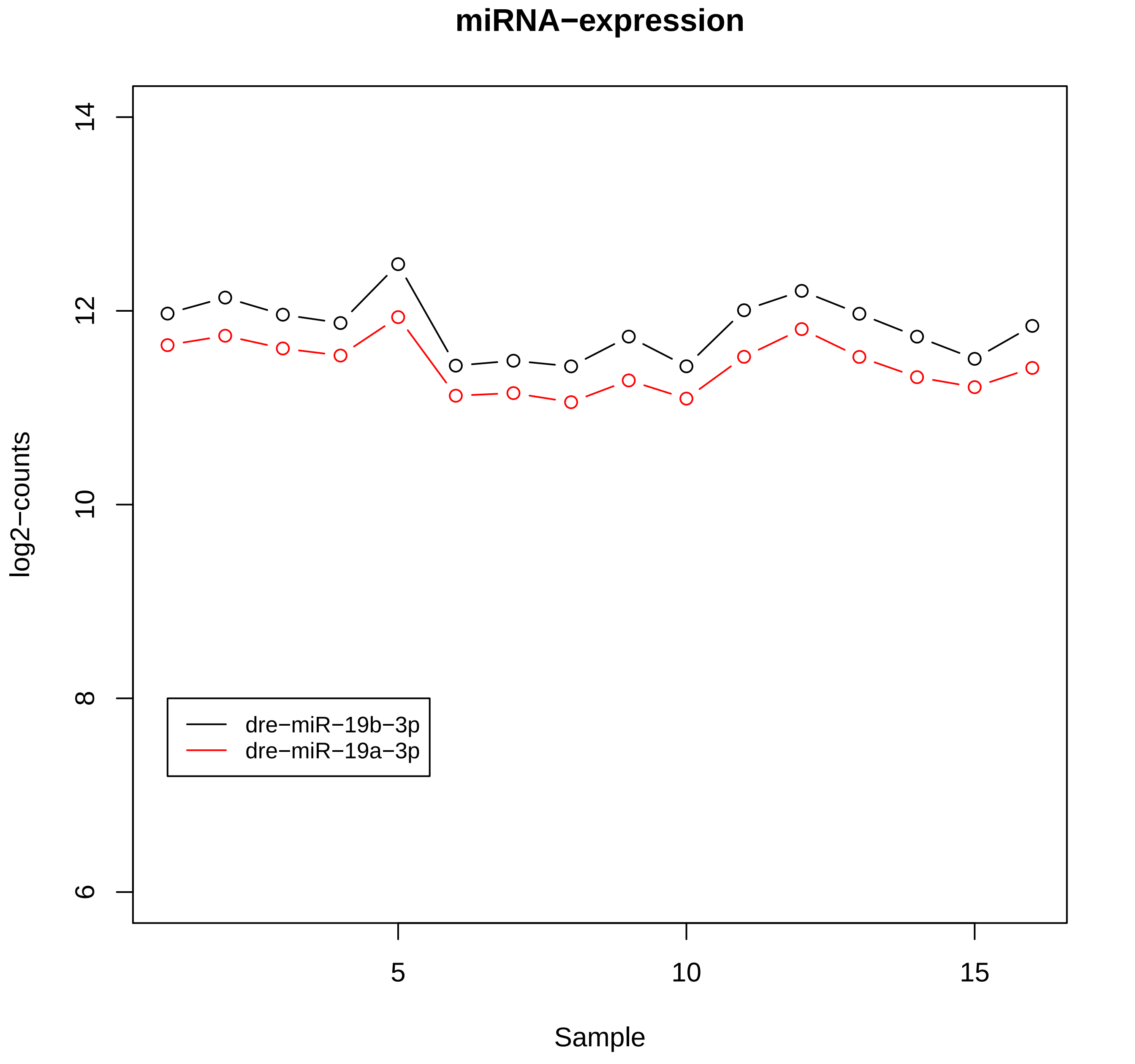

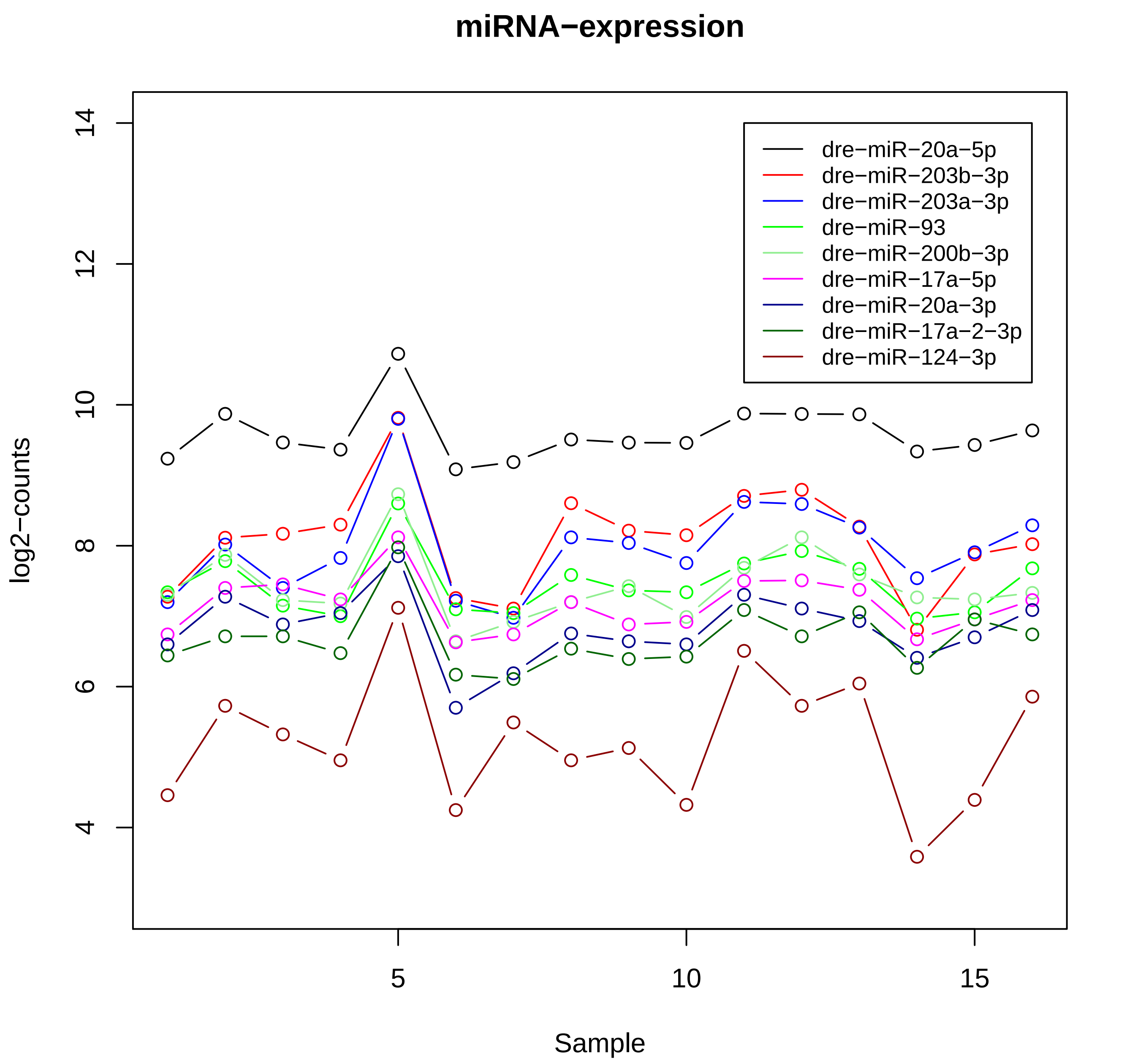

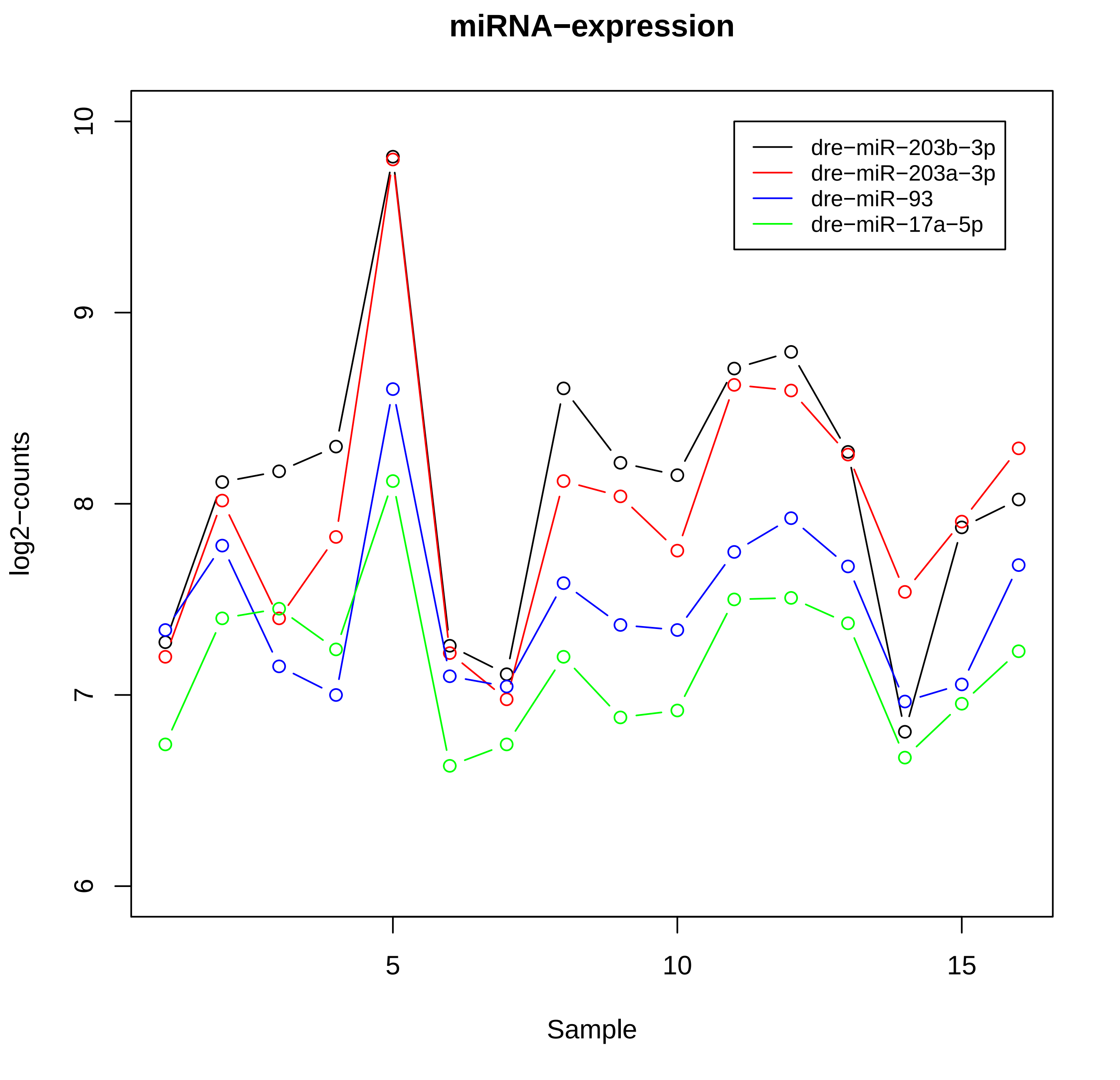

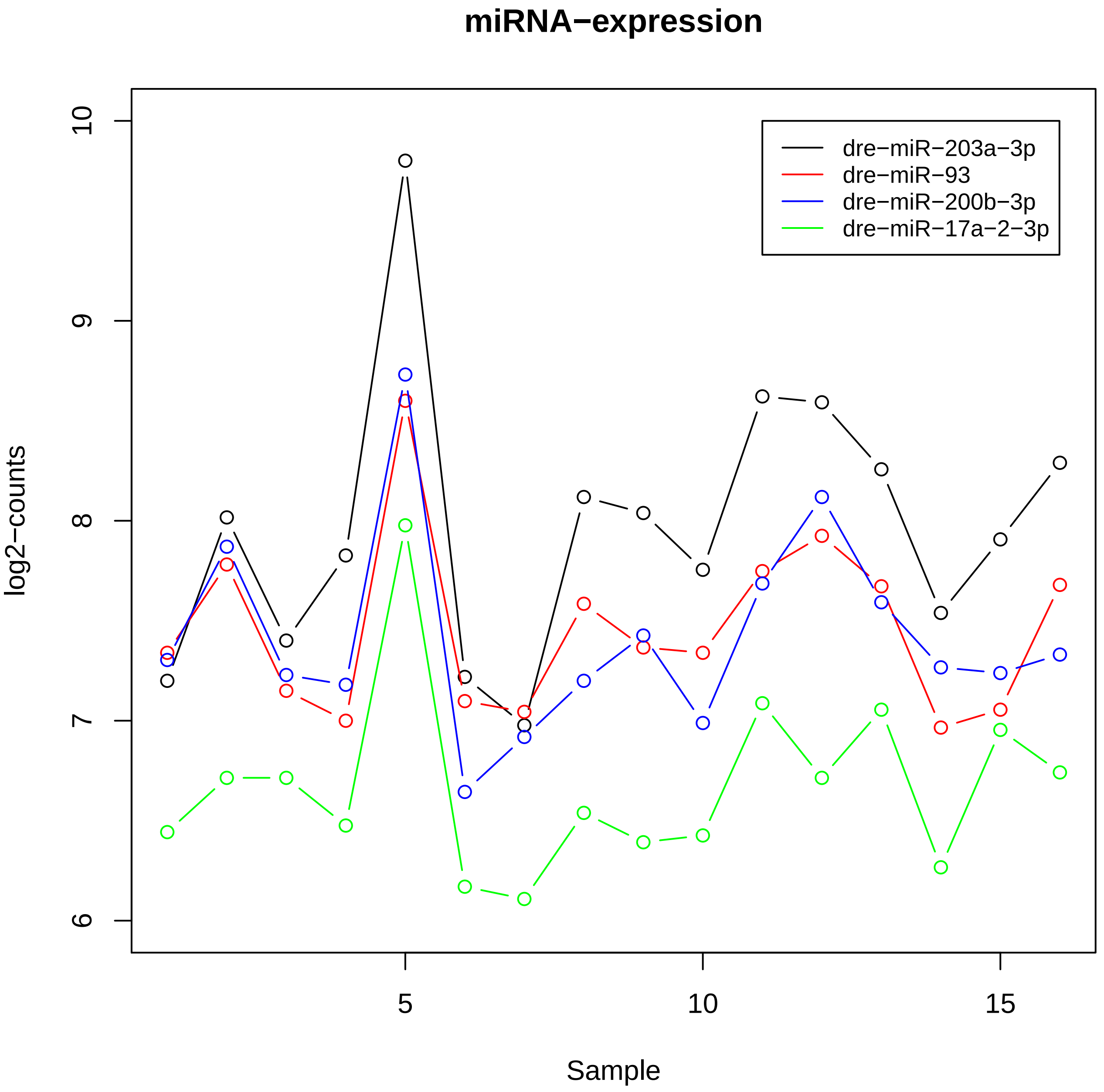

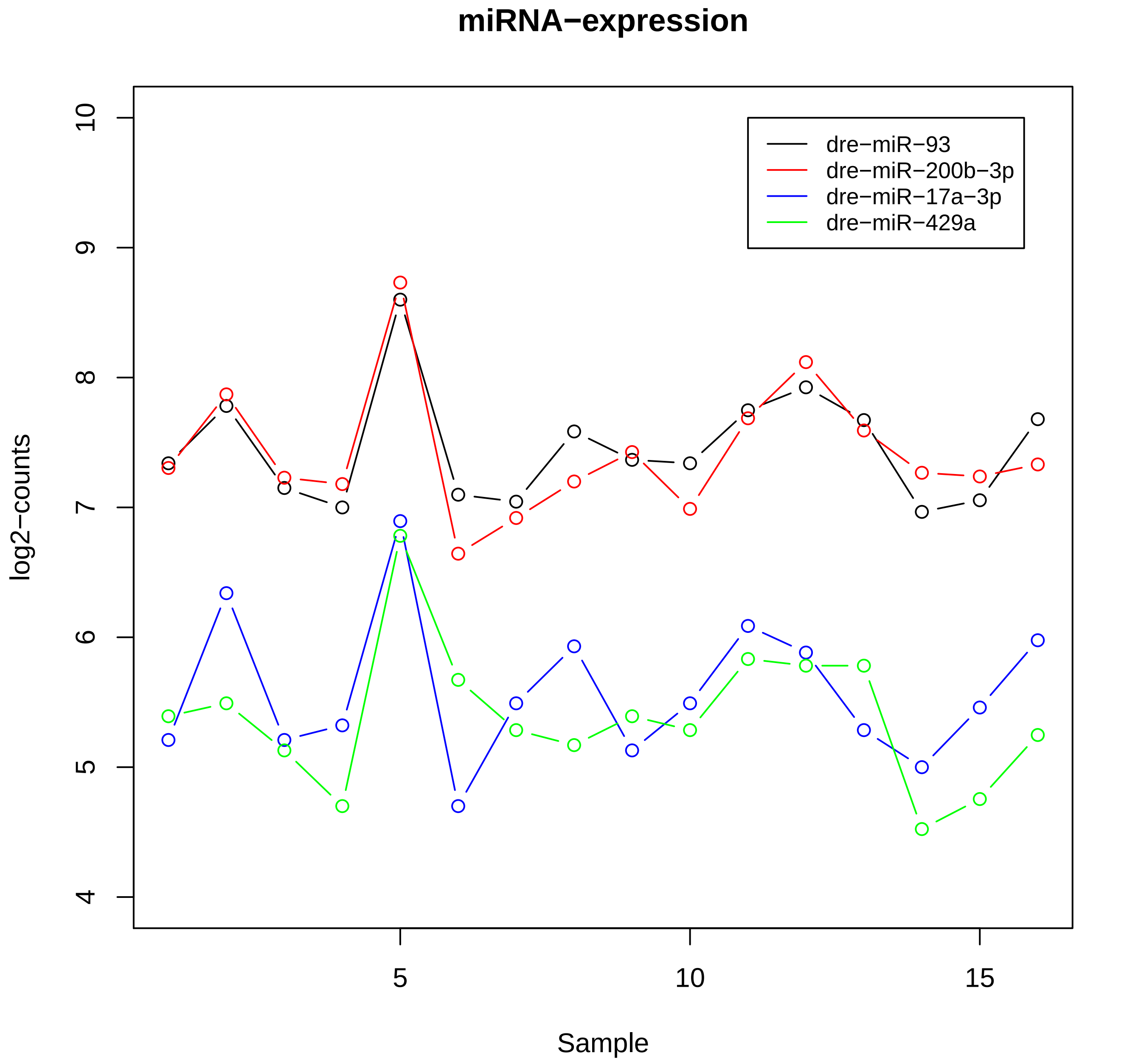

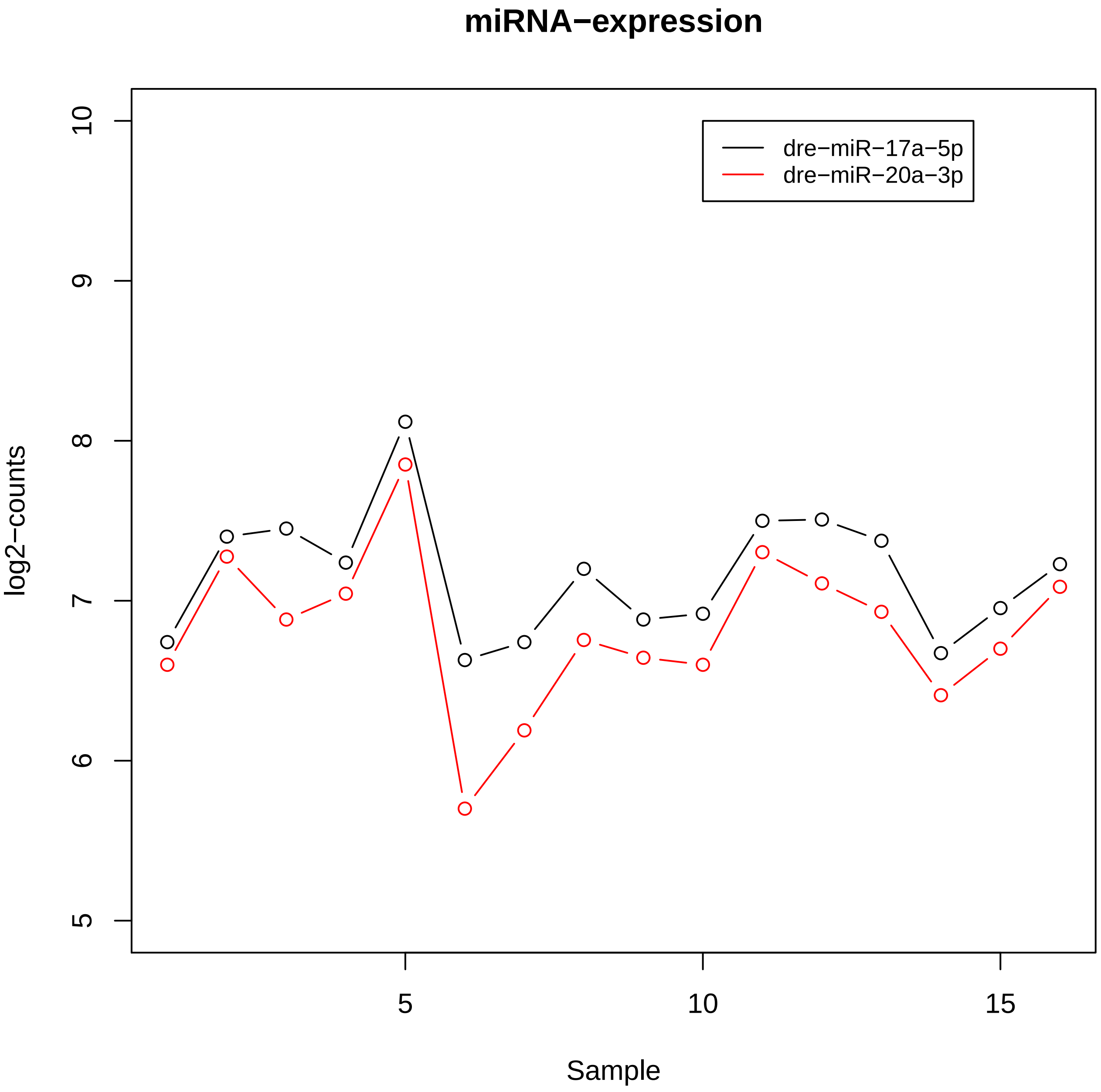

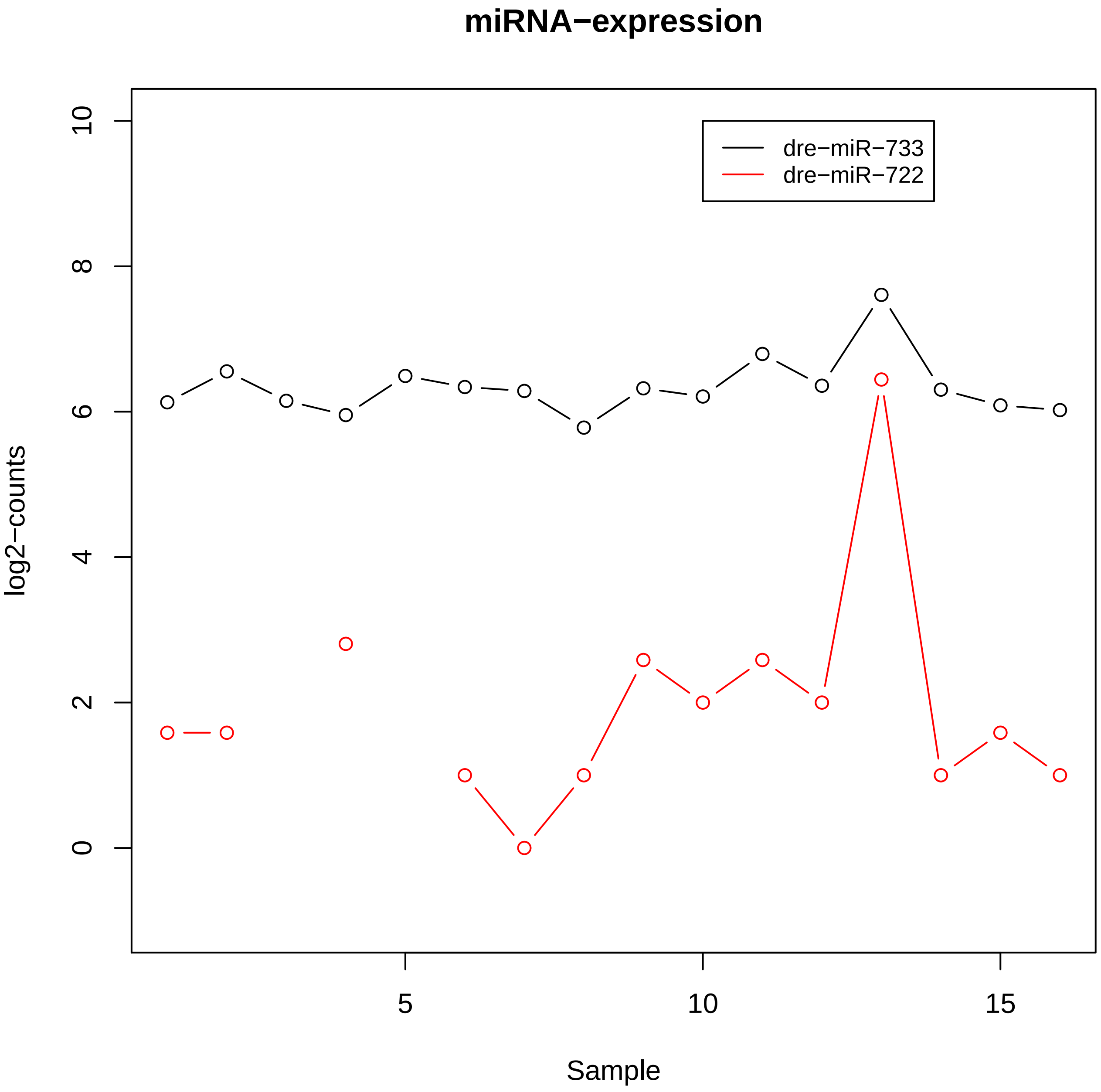
